# Multi-omic stratification of the missense variant cysteinome

**DOI:** 10.1101/2023.08.12.553095

**Authors:** Heta Desai, Samuel Ofori, Lisa Boatner, Fengchao Yu, Miranda Villanueva, Nicholas Ung, Alexey I. Nesvizhskii, Keriann Backus

## Abstract

Cancer genomes are rife with genetic variants; one key outcome of this variation is gain-of-cysteine, which is the most frequently acquired amino acid due to missense variants in COSMIC. Acquired cysteines are both driver mutations and sites targeted by precision therapies. However, despite their ubiquity, nearly all acquired cysteines remain uncharacterized. Here, we pair cysteine chemoproteomics—a technique that enables proteome-wide pinpointing of functional, redox sensitive, and potentially druggable residues—with genomics to reveal the hidden landscape of cysteine acquisition. For both cancer and healthy genomes, we find that cysteine acquisition is a ubiquitous consequence of genetic variation that is further elevated in the context of decreased DNA repair. Our chemoproteogenomics platform integrates chemoproteomic, whole exome, and RNA-seq data, with a customized 2-stage false discovery rate (FDR) error controlled proteomic search, further enhanced with a user-friendly FragPipe interface. Integration of CADD predictions of deleteriousness revealed marked enrichment for likely damaging variants that result in acquisition of cysteine. By deploying chemoproteogenomics across eleven cell lines, we identify 116 gain-of-cysteines, of which 10 were liganded by electrophilic druglike molecules. Reference cysteines proximal to missense variants were also found to be pervasive, 791 in total, supporting heretofore untapped opportunities for proteoform-specific chemical probe development campaigns. As chemoproteogenomics is further distinguished by sample-matched combinatorial variant databases and compatible with redox proteomics and small molecule screening, we expect widespread utility in guiding proteoform-specific biology and therapeutic discovery.

## INTRODUCTION

The average human genome is rife with sequence variation and differs from the reference at roughly 3.5 million sites^1^. This profound genetic variation gives rise to human diversity and disease. While the fraction of single nucleotide variants (SNVs) that occur in protein-coding make up a small fraction of all known variants, most known disease-causing mutations are found in protein coding sequences. Nearly all (>98%) of nonsynonymous protein-coding SNVs are missense variants that result in the substitution of single amino acids^2^. There are over 2 million coding mutations that have been identified in human cancers (Catalogue of Somatic Mutations [COSMIC] database), of which >90% are missense variants^3,4^. However, only a tiny fraction of these genetic variants (∼3,400) have been identified as putative missense driver mutations^5^ that confer selective growth advantages to cancer cells with the remaining mutations acting as “passengers.”

Quite surprisingly given the relative rarity of cysteine (2.3% of all residues in a human reference proteome)^6^, cysteine is the most commonly acquired amino acid due to somatic mutations in human cancers^7^. Given the unique chemistry of the cysteine thiol, including its nucleophilicity and sensitivity to oxidative stress, a subset of these residues almost unquestionably have a substantial impact on protein function. Exemplifying this paradigm, a number of driver mutations are gained cysteines, including Gly12Cys KRAS Tyr279Cys SHP2, Ser249Cys FGFR, and Arg132Cys IDH1^8–12^. A likely reason for the ubiquity of cysteine acquisition is the comparative instability of CpG motifs; C-T transitions are nearly ten times more common than other missense mutations in cancer^13^, and these transitions should favor gain-of-cysteine codons.

Due to its nucleophilicity and sensitivity to alkylation, cysteine residues have emerged as attractive sites to target with chemical probes. Covalent compounds can access small and poorly defined binding sites and can efficiently block high-affinity interactions (e.g. protein-protein interactions) or compete with high concentrations of endogenous biomolecules (e.g. ATP). There are numerous examples of cysteine-reactive clinical candidates and drugs, including the blockbuster covalent kinase inhibitors (e g. Afatinib and Ibrutinib^14–16^) and covalent compound that react with the Gly12Cys mutated oncogenic form of the GTPase KRAS (e.g. ARS-1620 and sotorasib^9,17–19^), a protein previously thought to be undruggable.

Mass spectrometry-based chemical proteomic methods, including those developed by our lab, have begun to unlock the therapeutic potential of the cysteinome. By capturing and enriching cysteines using highly reactive chemical probes, such as iodoacetamide alkyne (IAA) and iodoacetamide desthiobiotin, the studies have assayed the ligandability of upwards of 25% of all cysteines in the human proteome^20–29^. Cysteine chemoproteomics has even enabled the discovery of new lead molecules that target specific cysteines, including JAK^30^, SARM1^31^, PPP2R1A^32^, XRCC5^33^, NRB01^34^, and pro-CASP8^29^. Several new strategies have made substantial inroads into stratifying cysteine functionality to achieve function-first readouts of the likelihood of a covalent modification altering the labeled protein, including quantifying intrinsic cysteine nucleophilicity^25^, by pairing of chemoproteomics with CRISPR-base editing^35^, by performing proteomic stratification of covalent-modification induced altered protein complexes^36^, and our own work combining computational predictions of genetic pathogenicity with cysteine chemoproteomics^27^.

Single amino acid variants (SAAVs) encoded by missense mutations, including those that result in acquisition of cysteine, are almost universally missed by chemoproteomic studies. A key reason for this gap is that most genetic variants are not found in reference protein sequence databases used to identify peptides from acquired tandem mass spectrometry (MS/MS) data^20–29^. Understanding whether a genetic variant is translated into protein is a critical step for characterizing the functional impact and therapeutic relevance of genomic variation. Proteogenomic studies that implement custom variant-containing sequence databases for search have enabled proteome-wide detection of protein coding variants, including SAAVs and splice variants^37–43^. When compared to variant calling at the genomic level, the coverage of these studies remains comparatively small, spanning tens to hundreds of peptides, with the exception of recent studies employing ultra deep fractionation^44,45^ resulting in thousands of identified variants. These studies all share general data processing pipelines. Variant calling is performed on next-gen sequencing (NGS) data, then customized databases featuring both canonical protein sequences and sequences encoding SAAV-, insertion/deletions (indels)-, or splice variant-proteins are generated, using customized tools, such as Spritz^46^, CustomProDB^47^, Galaxy-P^48^, and sapFinder^49^. While targeted proteomics methods, such as parallel reaction monitoring (PRM) have enabled focused monitoring of high value variant-containing peptides^50^, including encoding driver mutations, the broader landscape of translated SAAVs remains to be fully explored.

There are two central complexities to these pipelines that have only recently begun to be addressed. The first challenge is that, by relying on exome-only sequencing and short read sequencing, the relative proximity of two or more variants in the same gene (whether they are on the same or opposite chromosomes) is not typically apparent. A notable exception is the recent integration of long read sequencing for de-novo database construction with sample-specific proteomics to characterize novel protein isoforms^51^. Consequently, multi-variant peptides are typically not detected by most proteogenomics workflows that rely on databases featuring either single-each or all-in-one SAAV-containing proteins. Such search strategies also introduce higher chances of false positive identification^52^. All possible cancer-derived aberrant peptide sequences, reflecting increased genetic complexity of tumor genomes, increases the size of the custom databases and thus search spaces. One solution to the false discovery rate (FDR) challenge is to calculate a class-specific FDR (separating the FDR calculations for the variant-containing peptides and reference peptides)^52^. An alternative strategy to ensure class-specific FDR control is to perform a 2-stage database search^53^. In this strategy, the first first search of acquired MS/MS spectra is performed against a reference database of canonical protein sequences. Subsequently, peptide to spectrum (PSM) matches identified with a certain high level of confidence (e.g. passing 1% FDR) are removed, and the remaining spectra are then searched against a variant-containing database. While implementation of such strategies in prior proteogenomic studies highlights the importance of rigorous statistical validation of identified variant-containing peptides^53–55^, the requirement for customized pipelines has so far limited widespread adoption.

Here we develop and deploy chemoproteogenomics as an integrated platform tailored to capture the missense variant cysteinome. Chemoproteogenomics unites a missense-variant focused proteogenomic pipeline with mass spectrometry-based cysteine chemoproteomics. By mining publically available datasets, including COSMIC, dbSNP, and ClinVar, we reveal that gain-of-cysteine variants are a ubiquitous consequence of genetic variation. We further reveal that DNA repair deficient cell lines are particularly enriched for acquired cysteines, together with a general high burden of rare and predicted deleterious variants. Guided by these discoveries, we generate combinatorial cell-specific custom databases built from whole exome and RNA-Seq data for eleven cell lines. Chemoproteogenomic analysis with a user-friendly FragPipe computational platform, extended to support 2-stage database search and FDR estimation, identified >1,400 total unique variants, including 629 chemoproteomic enriched variant-proximal cysteines and 103 gain-of-cysteines. Chemoproteogenomics also robustly identifies ligandable SAAVs that alter cysteine oxidation state and outperforms bulk proteogenomic analysis for capture of SAAVs with lower variant allele frequency. The utility of chemoproteogenomics is further showcased through our identification of iodoacetamide-labeled Cys67 (Cys91) in the highly variable peptide binding-groove of HLA-B. In sum, chemoproteogenomics sets the stage for enhanced global understanding of the functional and therapeutic relevance of the missense variant proteome.

## RESULTS

### High missense burden cancer cell lines are rich in acquired cysteines, including in census genes

Our first step to realize variant-directed chemoproteomics was to mine existing publicly available missense repositories to assess the scope of acquired cysteines present in cancer genomes (COSMIC) and healthy genomes (dbSNP) (**Figure 1A**). By doing so, we sought to achieve three goals: (1) validate prior reports of high cysteine acquisition in cancer ^7,56,57^ (2) determine whether cysteine acquisition is a privileged feature of cancer genomes, and (3) establish a panel of variant rich cell lines. We analyzed publicly available sequencing data of 1,020 cell lines, found in the Catalogue of Somatic Mutations in Cancer Cell Lines Project database^58,59^ (COSMIC-CLP, release v96), to establish a panel of high mutational burden tumor cell lines; our hypothesis was high missense burden cell lines would be enriched for acquired cysteine SAAVs, including those found in Census genes^60^ and residues that are driver mutations. The top 15 cell lines with the highest mutational burden (**Figure 1B, S1A, Table S1**) encode 77,693 total unique missense variants, which represents ∼18% of all unique missense variants in COSMIC-CLP.

**Figure 1.**
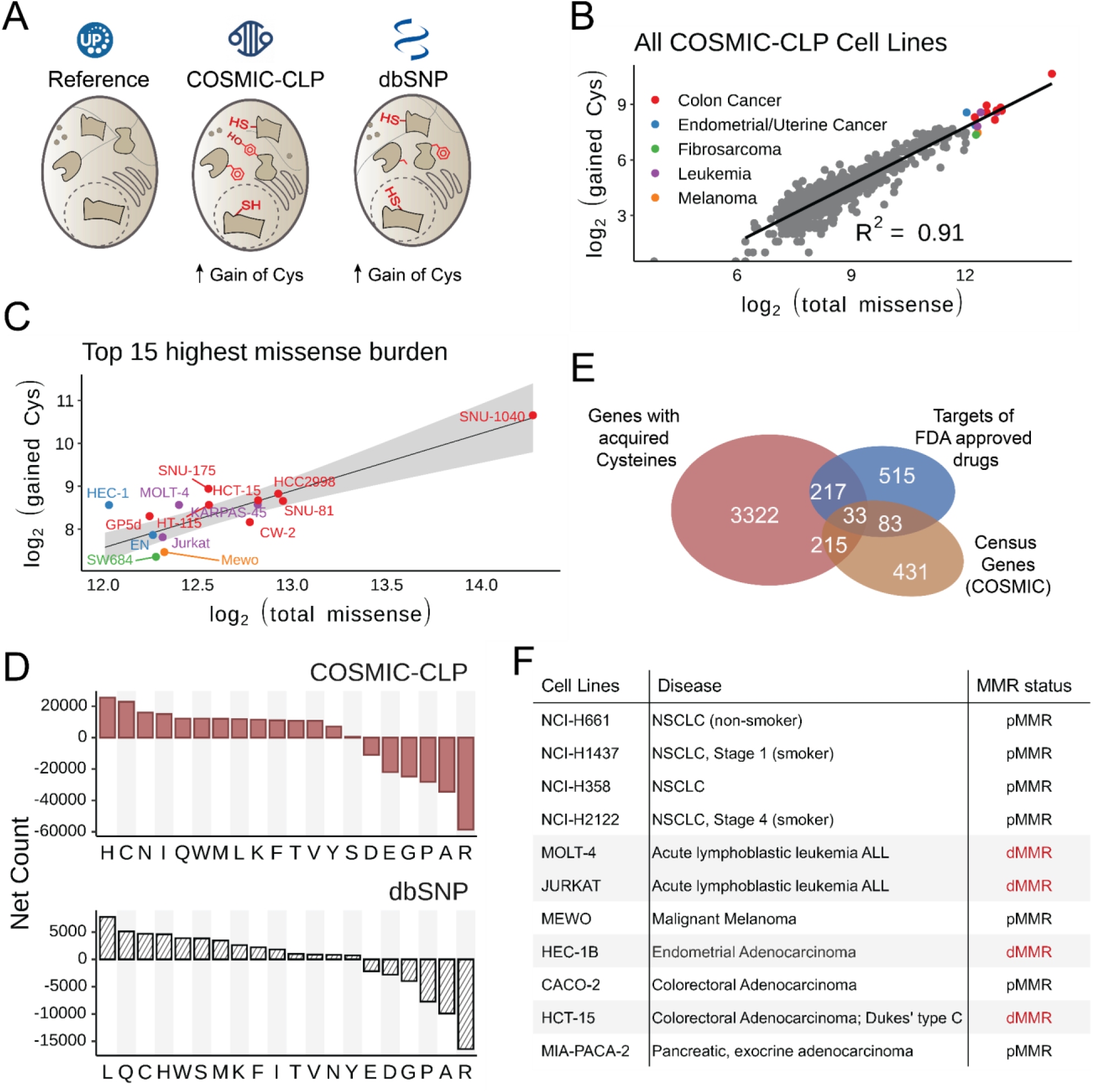
Acquired cysteines are prevalent across cancer genomes, particularly for high missense burden cell lines. A) The full scope of acquired cysteines in the COSMIC Cell Lines Project (COSMIC-CLP, cancer.sanger.ac.uk/cell_lines) (v96)^58,59^ and dbSNP (4-23-18)^73^ were analyzed. B) 1,020 cell lines stratified by number of gained cysteines and total missense mutations; color indicates cancer type for top 15 highest missense count cell lines. C) Top 15 cell lines with highest missense burden from panel B; linear regression and 95% confidence interval shaded in gray. D) Net missense mutations (gained-lost) from COSMIC-CLP (v96) and common SNPs (dbSNP 4-23-18). E) Overlap of genes with acquired cysteines in top 15 subset from panel B with Census genes and targets of FDA approved drugs. F) Panel of cell lines used in this study with MMR status (dMMR= deficient mismatch repair, pMMR=proficient mismatch repair). Data found in **Table S1**.

We next evaluated whether these identified missense-rich cell line genomes were similarly enriched for gained cysteine SAAVs. We calculated the net gain amino acid changes (total gained minus total lost) encoded by all coding missense variants in this cell line panel (**Figure S2**), which revealed a marked enrichment for acquired histidines and cysteines together with loss of arginine, both for the aggregate cell line panel and for individually analyzed cell line datasets (**Figure S3**). As calculations of net gain can fail to distinguish high versus low missense burden cell lines, we also further stratified these cell lines based on total gained and total lost amino acids (**Figure S1B, S4, S5**), which further substantiated the enrichment for gain-of-cysteine across all of the top 15 missense variant burden cell lines analyzed (**Figure 1C, S1B**). This marked cysteine enrichment in cancer cell line genomes is consistent with previously reported aggregate analysis, not stratified by cell line, of all available COSMIC missense data^7,56,57^. Our own analysis of all COSMIC-CLP mutations shows cysteine as the second most gained residue (**Figure 1D**). The genomes of the top 15 missense cell lines encoded 4,725 total gained cysteines, found in 3,688 genes. Showcasing the potential therapeutic relevance of this set, <10% of these identified genes have been targeted by FDA approved drugs^20,61^ (**Figure 1E**). Notably, 219/738 Census genes (v98) were found to harbor one or more gained cysteines, including NRAS (G12C), which is found in the Molt-4 cell line; TP53 (R273C) found in the KARPAS-45 cell line; GNAS (R218C) in the CW-2, SNU-175, and HT-115 cell lines; FBXW7 (R505C) in Jurkat and KARPAS-45 cell line; ASXL1 (W796C) found in HCT-15 cell line, and KEAP1 (Y33C) found in the Hec-1 cell line (**Table S1**).

### dMMR cell lines are enriched for SAAVs, including acquired cysteines

Cancer genomes display characteristic patterns of mutations, or signatures, that have developed from biological processes specific to the course of the cancer^62,63^. Endogenous and exogenous sources of DNA damage, left uncorrected due to faulty repair pathways, often lead to high tumor mutational burdens. Microsatellite instability (MSI) is a hypermutable phenotype caused by deficiency in mismatch repair (dMMR). High MSI tumors have higher mutational burdens; the converse is not true as high mutational burden tumors do not always display MSI^64^. Eight out of fifteen of the top missense burden cell lines reported in COSMIC were observed to be derived from colorectal carcinoma (CRC) (**Figure 1B, S3**). As ∼15% of CRCs are reported to have elevated MSI^65–67^, this high CRC missense burden is to be expected^64,68^. While Jurkat, Molt-4 and Hec-1B cells are not CRC, both have previously been reported as dMMR with mutations in mismatch repair machinery^69,70^. Unexpectedly, MeWo cells, which are derived from metastasized melanoma and reported to be microsatellite stable (MSS)^71^, also exhibited a high burden of missense mutations. The majority of missense rich cell lines, including the dMMR lines were observed to encode between 200 and 500 acquired cysteine SAAVs (**Figure S1B**). However, a significant depletion of gained cysteines relative to total variant burden was observed for MeWo and SW684 (**Figure 1C**).

### Acquired cysteines are ubiquitous in both healthy and diseased genomes

We next asked whether this marked enrichment for gained cysteines was specific to cancer genomes or a more universal consequence of human genetic variation, with the overarching goal of facilitating efforts to pinpointing acquired cysteines with therapeutic relevance. Complicating matters, gain-of-cysteine missense variants are also expected to be ubiquitous in healthy genomes, due to the comparative instability of CpG–a key consequence of this instability is the frequent loss of arginine codons (4/6 CG dinucleotides)^72^. We aggregated and quantified the amino acid changes resulting from common missense variants reported by dbSNP^73^, a repository of single nucleotide polymorphisms and ClinVar^74^, a repository of variants with reported pathogenicity. We find that cysteine acquisition is the third most common consequence of missense variants identified in dbSNP (**Figure 1D, Table S1**) for common variants—common variants are defined by NCBI as of germline origin and/or with a minor allele frequency (MAF) of >=0.01 in at least one major population, with at least two unrelated individuals having the minor allele. Analogous stratification of variants reported by ClinVar also revealed a preponderance of gained cysteines compared with lost cysteines, albeit to a more modest degree than that observed for cancer genomes (**Figure S6 and Table S1**). For the pathogenic variant subset of ClinVar, both gain– and loss-of-cysteine and gain-of-proline were frequently observed (**Figure S6**).

### An expanded cell line panel incorporates high value acquired cysteines

Across the >2 million missense variants reported in COSMIC, 52 acquired cysteines are reported as putative driver mutations (dN/dS values)^75^ in the Cancer Mutation Census (**Table S1**). Consequently, nearly all acquired cysteine SAAVs are of uncertain functional significance for tumor cell growth and survival. Given that one of our key objectives is to enable rapid proteomic identification and subsequent electrophilic compound screening of functional variants, we next stratified the top missense variant cell lines based on known driver mutations and damaging variants. We find that top missense cell lines that are readily available for purchase encode NRAS G12C, KRAS G12D, PIK3CA E545K, and TP53 R248Q variants among other known driver mutations (**Table S1**). Given the considerable interest in targeting G12C KRAS, we opted to add several KRAS mutated cell lines to our panel (MIA-PACA-2, H2122, and H358) in order to favor detection of the G12C peptide. Notably, the smoking-associated mutational signature is C→A/G→T^76^, which should favor gain-of-cysteines. Therefore, we additionally sought to test whether smoking associated NSCLC-derived H2122 and H1437 adenocarcinoma cell lines would be enriched for acquired cysteines when compared to other proficient mismatch repair (pMMR) cell lines, including lung cancer cell lines (H358 NSCLC and H661 metastatic large cell undifferentiated carcinoma (LCUC) lung cancer cell lines). Lastly, we opted to include CACO-2 cells, an MSS CRC cell line, to test the feasibility of capturing driver mutations located proximal to chemoproteomics detectable cysteines—Caco-2 cells express mutant SMAD4 (D351H), a variant implicated in blocking SMAD homo– and hetero-oligomerization^77^ and located proximal to two previously chemoproteomics detected cysteines (C345 and C363)^21,22^. Our prioritized cell line panel features 11 cells lines in total (2 female and 9 male) spanning 6 tumor types and encoding 22,559 somatic variants and 1,296 somatic acquired cysteines, as annotated by COSMIC-CLP (**Figure 1F, Table S1-S2**), with aggregate enrichment for gained cysteines observed for the entire panel (**Figure S7, S8**). Of the proteins that harbor gained cysteines, 486 are Census genes and 5% are targeted by FDA approved drugs (**Table S1**).

### dMMR cell lines are enriched for rare predicted missense changes, including acquired cysteines

Given the preponderance of acquired cysteine SAAVs observed across COSMIC, ClinVar, and dbSNP, we postulated that cancer genomes would be enriched for both rare and common gain-of-cysteine mutations. To both test this hypothesis and enable the building of sequence databases for proteogenomics search, we sequenced exomes and RNA of our cell lines and subjected NGS reads to variant-calling (**Figure 2A, Figure S9**). For all 11 cell lines sequenced, we identified on average 82% of the variants reported in COSMIC-CLP and 70% of missense mutations reported by Cancer Cell Line Encyclopedia (CCLE)^71^ databases (**Table S2**). Driver mutations (CMC significant, dN/dS q-values) identified include KRAS G12C for MIA-PACA-2, H358, and H2122 cell lines, PIK3CA E545K in HCT-15, and FBXW7 R505C in Jurkat cells (**Table S2**). 9,190 total rare variants were identified that had been not previously reported in COSMIC-CLP, including 435 variants encoding acquired cysteines (**Table S2**).

**Figure 2.**
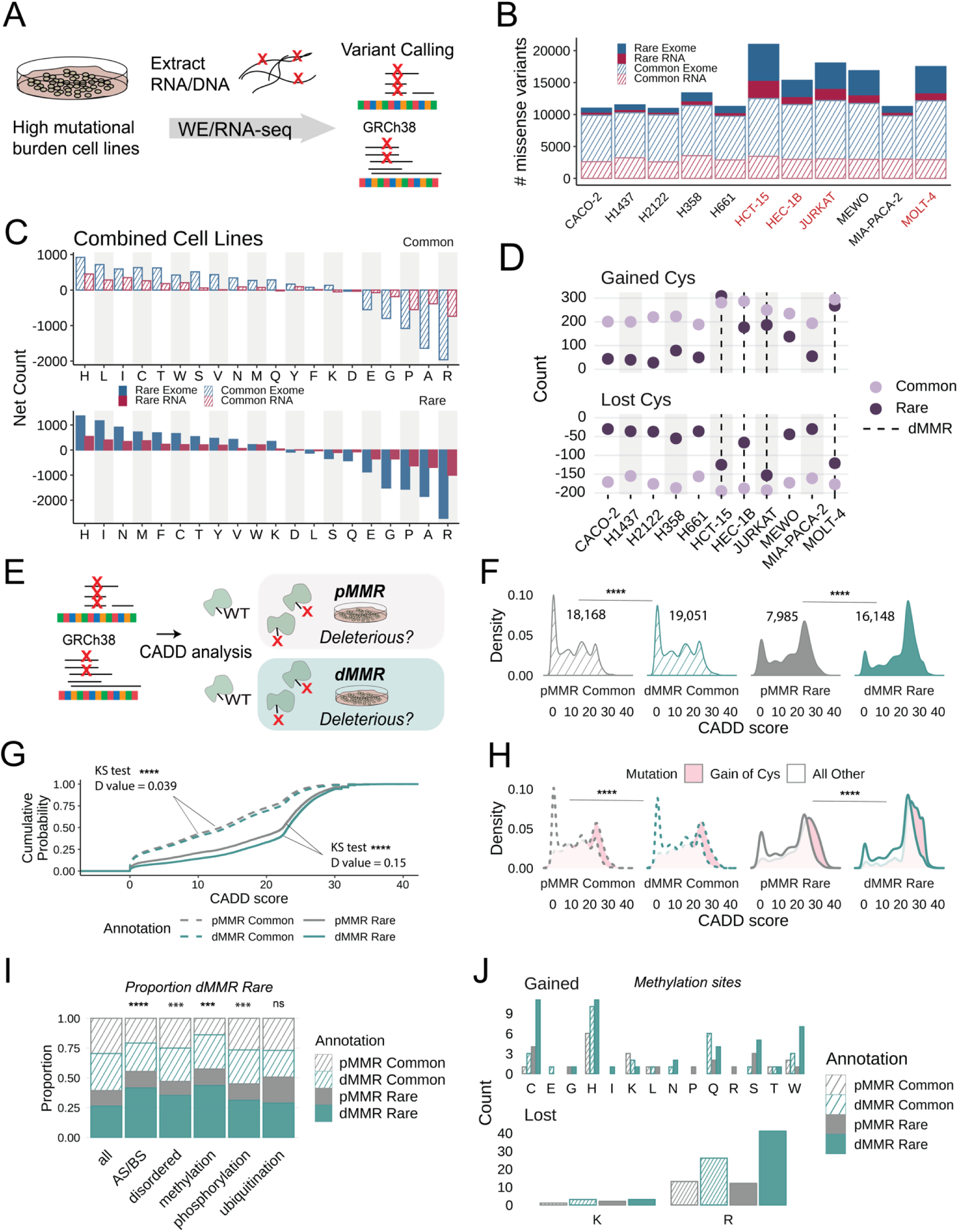
dMMR cell lines are enriched for rare, predicted deleterious gain-of-cysteine mutations. A) Sequencing portion of the ‘chemoproteogenomic’ workflow to identify chemoproteomic detected variants– extracted genomic DNA or RNA from cell lines undergo sequencing followed by variant calling using Platypus (v0.8.1)^83^ and GATK-Haplotype Caller (v4.1.8.1)^84^ for RNA and exomes respectively and predicted missense changes were computed. B) Total numbers of missense mutations identified from either RNA-seq or WE-seq; stripe vs solid denotes common and rare variants, red text indicate dMMR cell lines. C) Net amino acid changes for all cell lines combined. D) Totals of gained and lost cysteine in each cell line separated by rare and common variants, dashed line indicates dMMR cell lines. E) Scheme of CADD score analysis for two dMMR and non-dMMR cell lines. F) Distribution of CADD scores for indicated variant grouping; statistical significance was calculated using Mann-Whitney U test, **** p < 0.0001. G) Empirical cumulative distributions (ECDF) were computed for CADD scores with indicated grouping; statistical significance was calculated using two-sample Kolmogorov-Smirnov test, **** p < 0.0001. H) CADD score distributions for cysteine gained amino acid indicated separated by grouping; statistical significance between gained Cys values was calculated using Mann-Whitney U test, **** p < 0.0001. I) Proportion of variants belonging to the indicated sites; AS/BS = in or near active site/binding site as annotated by UniProtKB or Phosphosite; statistical significant calculated using two-sample test of proportions, *** p < 0.001, **** p < 0.0001, ns p > 0.05. J) Amino acid changes at protein methylation sites as identified by Phosphosite. Data found in **Table S2**.

As with our analysis COSMIC-CLP (**Figure 1B**), we detected a high missense burden for the dMMR cell lines compared to the pMMR cell lines. MeWo cells were an exception, with a missense burden comparable to that of the dMMR cell lines (**Figure 2B)**. Analysis of DNA damage repair-associated genes revealed specific mutations (**Table S2**), including DDB2 R313* in MeWo cells, which provide an explanation for the previously unreported high missense burden— inactivating mutations in DDB2 are implicated in deficient nucleotide excision repair^78^.

We next subsetted the data into rare and common variant categories, using dbSNP common variants (04-23-2018 00-common_all.vcf.gz) (**Table S2**)^73^. The dMMR cell lines, together with the MeWo cells, have proportionally more rare variants compared to common variants (**Figure 2B**), irrespective of sequencing coverage (**Figure S10**). Further SAAV analysis revealed net gain of histidine, isoleucine, and cysteine as the most frequent amino acids gained across the common and rare subsets (**Figure 2C**). We find that cysteine acquisition is a more frequent consequence of common variants detected in pMMR cell lines (**Figure 2D**).

In contrast with the common variants, the net gained SAAV signatures encoded by rare variants differed markedly between dMMR and pMMR cell lines (**Figure 2D**, **S11-13).** No significant difference between the number of gained cysteines was observed for the smoking-associated lung cancer cell lines (**Figure S14**). By contrast, in the dMMR cell lines, we detected a sizable increase, when compared to the pMMR cell lines, of acquired rare SNVs encoding Cys, along with His, Ile, Asn, Tyr, and Tryp (**Figure 2D, Figure S11**). Beyond cysteine acquisition, the SAAV signature for MeWo cells was observed to be distinct, with pronounced gain-of rare Phe and Lys detected (**Figure S11-13**), consistent with UV radiation induced pyrimidine dimers, which result in gain-of F and K (**Figure S15**, **Table S2**). These findings together with our analysis of the top missense cell lines in COSMIC-CLP indicate that previously reported widespread cysteine acquisition in cancer genomes is predominated by mismatch repair deficient cell lines.

### Rare gained cysteines in dMMR cell lines are enriched for high CADD scores

With the overarching goal of facilitating identification of likely functional variants, we next stratified the predicted deleteriousness of the identified missense variants (**Figure 2E, Table S2**). We focused on the Combined Annotation Dependent Depletion (CADD) score, due to its high reported specificity and sensitivity^79^ and our prior findings that showed strong association between cysteine functionality and high CADD score^27^. Unsurprisingly, our analysis revealed higher CADD scores for rare variants compared to common variants, across the cell line panel (**Figure 2E, Table S2**). More unexpectedly, we observed a more marked increase in the predicted pathogenicity of the rare variants detected in dMMR cell lines compared with pMMR cell lines (the top 1% most predicted deleterious mutations have CADD phred-scaled scores > 20) (**Figure 2F-G, S16-17)**. This enrichment for high CADD score rare variants held true for the MeWo cells. Further stratification by specific gained or lost amino acids (**Figure 2H, Figure S18-21**), revealed that gained cysteine missense are the most significantly enriched for high predicted deleterious scores across all pMMR and dMMR cell lines (**Figure S19**, **Table S2**)—a notable exception are the MeWo cell line variants for which gain-of Phe, Lys, and Leu codons are the most high CADD scoring variants (**Figure S22**).

As only a small fraction of the acquired cysteines are known driver mutations, we next restricted our analysis to include only the 388 total variants localized to hotspot mutations, as annotated by CCLE and The Cancer Genome Atlas (TCGA). We find that gain of cysteine within TCGA hotspot mutations is markedly enriched for high CADD score variants (**Figure S21**). Notable high CADD score hotspot acquired cysteines include the tumor suppressor FBXW7 R505C in Jurkat cells, the metalloprotease ADAMTS1 R604C in Molt-4 cells, and exrin-associated protein SCYL3 R61C in MeWo cells. 98% (50/51) of these cysteines are gained due to loss of arginine, which aligns with the observed parallel enrichment for high CADD scores at loss of arginine hotspot variants (**Figure S23**).

### dMMR rare variants are enriched for proximity to known functional sites

To further broaden our understanding of the functional landscape of cysteine acquisition, we also analyzed proximity to known functional sites and sites of post translational modification (**Table S2**). We find that the dMMR rare variant set is enriched for known proximal active site/binding site residues (**Figure 2I**). Intriguingly, analysis of known PTM modified sites reported by Phosphosite^80^ revealed a significant association between arginine methylation sites and rare variants in dMMR cell lines (**Figure 2I**). These findings are consistent with loss of arginine as a frequent consequence of exonic CpG mutability^72,81^ together with roles of MMR in protecting against CpG associated deamination^82^. As 60% of the gained cysteines in our data resulted from loss of arginine (**Figure S24**), we expected that many of these variants will result in altered PTM status (**Figure 2J**).

### Variant peptide identification enabled by MSFragger 2-stage database search and false discovery rate (FDR) estimation

To enable chemoproteomic detection of acquired cysteine SAAV-containing peptides and SAAVs found in peptides with canonical cysteines, we next established a customized proteogenomics pipeline **(Figure 3A, B**). Motivated by the prior report^38^ that demonstrated proteogenomic sample searches performed with sample-specific databases both improved coverage (∼45% more variants) and decreased rates of SAAV peptide false discovery, we generated cell line-specific variant peptide databases from HEK293T RNA-seq data (**Figure 3A, Table S3**). Next, to afford a reduction to the likelihood that a variant peptide will be mismatched to wild-type spectra^53^, we established a 2-stage database search and FDR control scheme (**Figure 3B**), usingMSFragger (v3.5)/Philosopher^85,86^ command line pipeline within FragPipe computational platform (detailed in Methods).

**Figure 3.**
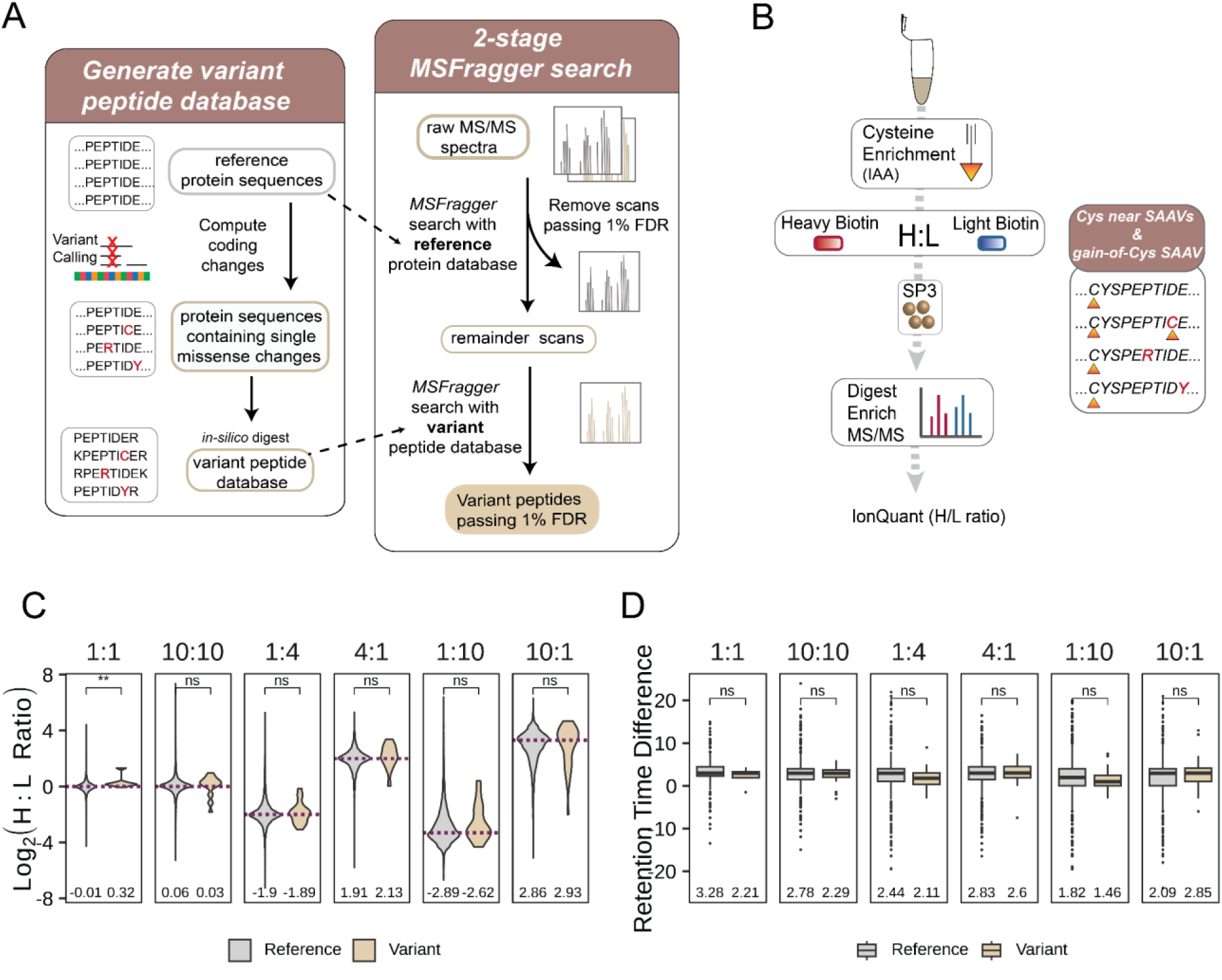
Variant peptide identification implementing an MSFragger-search pipeline. A) 2-stage MSFragger-enabled variant searches–variant databases are generated from non-redundant reference protein sequences that are *in-silico* mutated to incorporate sequencing-derived missense variants followed by 2-stage MSFragger/PeptideProphet search to identify confident variant-containing peptides. First, raw spectra are searched against a normal reference protein database, confidently matched spectra (passing 1% FDR) are removed and remainder spectra are searched with a variant tryptic database. B) Chemoproteomics workflow to validate heavy and light biotin^87^. HEK293T cell lysates were labeled with pan-reactive iodoacetamide alkyne (IAA) followed by ‘click’ conjugation onto heavy or light biotin azide enrichment handles in known ratios. Following neutravidin enrichment, samples are digested and subjected to MS/MS analysis. C) Heavy to light ratios (H:L) from triplicate datasets comparing identifications from reference and variant searches; mean ratio value indicated, *dashed lines* indicate ground-truth log2 ratio, statistical significance was calculated using Mann-Whitney U test, ** p < 0.01, ns p > 0.05. D) Retention time difference for heavy and light identified peptides for reference and variant-searches; mean value indicated, statistical significance was calculated using Mann-Whitney U test, ns p > 0.05. Data found in **Table S3**.

We then subjected our chemoproteogenomics pipeline to benchmarking by generating a set of high coverage cysteine chemoproteomics datasets (**Figure 3B**) in which cell lysates labeled with iodoacetamide alkyne (IAA)^25^ and conjugated isotopically labeled ‘light’ (^1^H_6_) or’ heavy’ (^2^H_6_) biotin-azide reagents^87^ (+6 Da mass difference between the reagents) were combined pairwise in biological triplicate at different H/L ratios (1:1,10:10, 1:4, 4:1, 1:10, and 10:1). By searching these datasets using our 2-stage search, we sought to validate the accuracy of variant identification. Peptide quantification using IonQuant^88,89^, following the workflow shown in **Figure 3A**, revealed MS1 intensity ratios for both canonical and variant peptide sequences that matched closely with the expected values (**Figure 3C, Table S3**). We also compared the retention times of the heavy– and light-peptides and observed an ∼2-3 sec shift for the deuterated heavy sequences for both the variant and canonical peptide sequences (**Figure 3D, Table S3**). These retention time shifts are consistent with our previous study^87^ and with prior reports^90,91^. Analogous to studies that utilize isotopically enriched synthetic peptide standards to validate novel peptide sequences^92–94^, the observed co-elution of both heavy and light variant peptides provides further evidence to support the low FDR of our data processing pipeline. Lastly, the high concordance between observed and expected MS1 ratios provides compelling support for the use of the heavy and light biotin azide reagents in competitive cysteine-reactive compound screens, in which elevated MS1 intensity ratios are indicative of a compound modified cysteine.

### Chemoproteomics with combinatorial databases improves coverage of acquired cysteines and proximal variants

We next set out to apply our validated search scheme for chemoproteogenomic variant detection (**Figure 4A**). Inspired by the recent report^95^ of combinatorial databases to improve detection of proximal SAAVs—we expect such variants to be prevalent in heterogeneous cell populations, such as a mismatch repair deficient tumor cell line— we established an algorithm (**Figure S27**) to generate all combinations of SAAVs derived from both RNA/WE-seq data within 30 amino acids flanking the variant site. These combinations were then converted into a peptide FASTA database containing two tryptic sites flanking each variant site (**Figure 4B**). On average, >4,500 total multi-variant peptide sequences were generated per cell line. Our approach differs from most prior custom database generators, which offer ‘Single-Each’^47,92,96,97^ or ‘All-in-One’ outputs^98,99^ for the former, all protein sequences harbor one SAAV each; for the latter, each protein harbors all SAAV detected. While establishing our combinatorial databases, we observed that a small number of highly polymorphic genes (**Table S4**) markedly increased database size—exemplifying this increased complexity, upwards of 1 billion combinations (*2^n –1*) are possible for protein sequences with 30 or more SAAVs. To determine the practical limit for the number of SAAVs/protein, we performed test searches where we limited the numbers of variants to combine (**Table S4**). We find that nearly all variants are retained with databases that include combinations for proteins with up to 25 variants (**Table S4**). For the small set of highly polymorphic protein sequences (e.g. HLA, MUC, and OBSCN, (**Table S4**), Single-Each sequences were searched (**Figure S27**).

**Figure 4.**
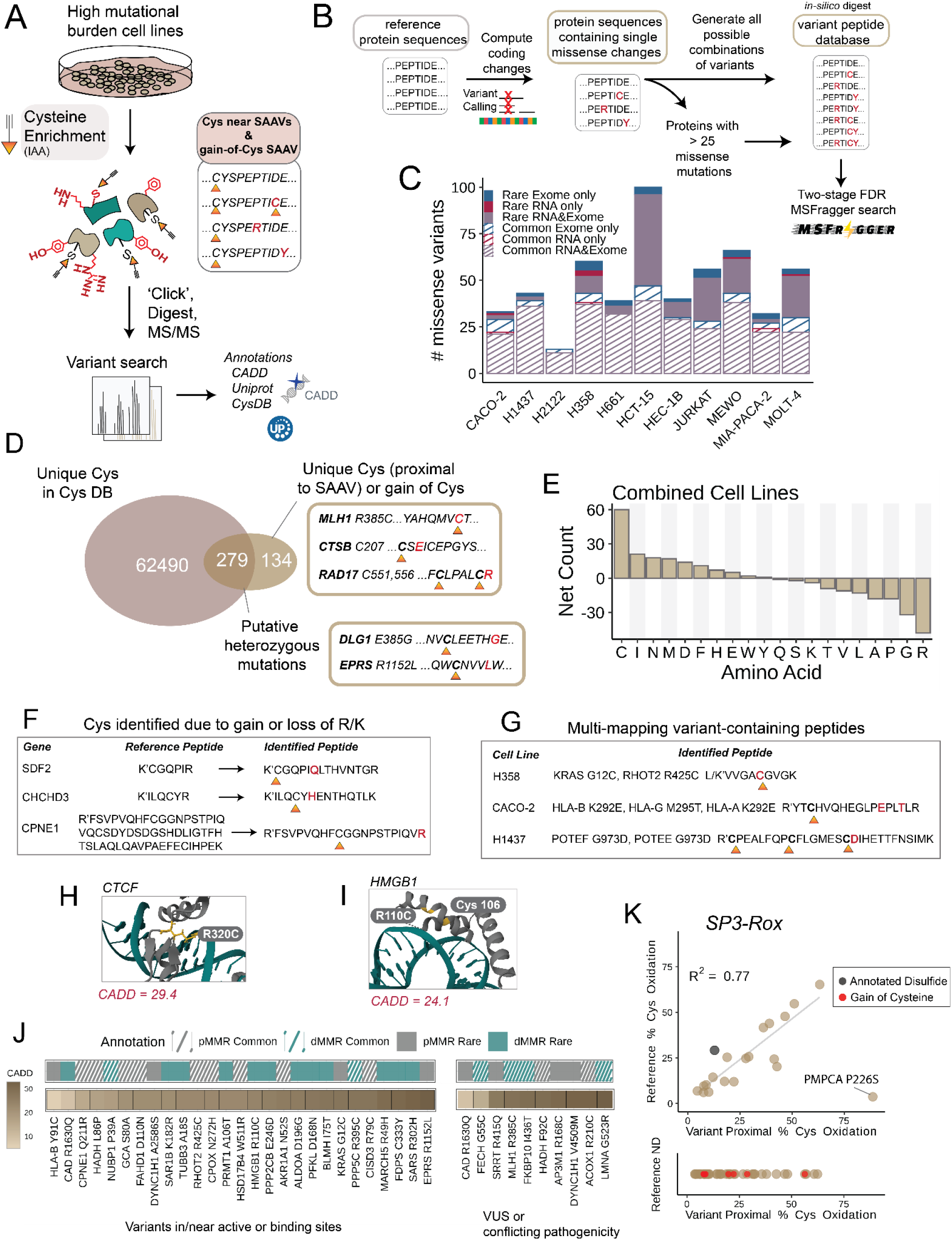
Variant peptide identification on tumor cell lines. A) Cell lysates were labeled with pan-reactive iodoacetamide alkyne (IAA) followed by ‘click’ conjugation onto biotin azide enrichment. Samples were prepared and acquired using our SP3-FAIMS chemoproteomic platform^22,23,107^ using single pot solid phase sample preparation (SP3)^108^ sample cleanup, neutravidin enrichment, sequence specific proteolysis, and LC-MS/MS analysis with field asymmetric ion mobility (FAIMS) device^109^. Experimental spectra are searched using the custom fasta for variant identification. Sample set includes both reanalysis of previously reported datasets from Yan et al. (Molt-4, Jurkat, Hec-1B, HCT-15, H661, and H2122 cell line) with newly acquired datasets (H1437, H358, Caco-2, Mia-PaCa-2 and MeWo cell lines). B) Non-synonymous changes are incorporated into reference protein sequences and combinations of variants are generated for proteins with less than 25 variant sites to make customized fasta databases. Details in methods. C) Total numbers of unique missense variants identified from either RNA-seq or WE-seq or both after using 2-stage MSFragger search and philosopher validation from duplicate datasets; stripe vs solid denotes common and rare variants, red text indicate dMMR cell lines Indicated is sequencing source and type of variant. D) Overlap of identified cysteines from variant searches with cysteines in CysDB database^20^. E) Net amino acid changes for all cell lines combined F) Example of cysteies identified from loss of R/K peptides G) Examples of multi-mapping variant sites. H) Crystal structure of CTCF indicating detected Cys320 (*yellow*) and DNA-binding site (PDB: 5T0U). I) Crystal structure of HMGB1 indicating detected Cys110 and nearby Cys106 (*yellow*) (PDB: 6CIL). J) Variants identified in or near active and binding sites with CADD score, common/rare, cell line dMMR/pMMR annotations. K) Re-analysis of SP3-Rox^106^ oxidation state data in Jurkat cells. Data found in **Table S4**.

Next, for all 11 sequenced cell lines (**Table S2**), we prepared and acquired a set of high coverage cysteine chemoproteomics datasets (**Figure 4A**). In aggregate, 32,638 total canonical cysteines were identified on 7,233 total proteins, with 9,349 cysteines unique to individual cell lines and 25,223 shared across the entire dataset (**Figure S25, Table S4**). 2,318 cysteines on 1,406 total proteins had not previously been reported in the CysDB database^20^ (**Figure S26**). 2-stage MSFragger search using our sample specific combinatorial databases identified a total of 59 gained cysteines and 302 SAAVs located proximal to 343 reference cysteines (**Figure 4C, Table S4**). 74 canonical sequence cysteines located proximal to variants and 60 acquired cysteines had not been previously reported in CysDB (**Figure 4D**)^20^. Notable examples of acquired cysteine variants not reported in CysDB include acquired cysteines KRAS G12C and PRKDC R2899C. Consistent with the aforementioned genomic data findings, we observe arginine as the most frequently lost out of detected Cys-proximal SAAVs (**Figure 4E**). We detect 15 total cysteines in peptides that harbor gain/loss of arginine that were previously too long or too short to be identified (**Figure 4F, Table S4**). For the cysteine protease cathepsin B (CTSB), we identify Cys207 in HCT-15 cells which was not identified in CysDB–a K209E mutation that creates a longer tryptic peptide sequence compared to reference sequence (‘CSK’ to ‘CSEICEPGYSPTYKQDK’). In the well-studied Jurkat proteome, we detect stromal cell derived factor 2 SDF2, Cys88, which is also not reported in CysDB, is found in a peptide harboring a proximal R93Q mutation that creates a longer, detectable peptide sequence (‘CGQPIR’ to ‘CGQPIQLTHVNTGR’). Showcasing the utility of the combinatorial exome and RNA-seq SAAV databases, we identify six multi variant-containing peptides (**Table S4**). One noteworthy example is the peptide L86P/F92C peptide from the mitochondrial enzyme HADH, which catalyzes beta-oxidation of fatty acyl-CoAs—two variants, one from RNA-seq and one from exome-seq were detected in this peptide. For the I105V, A114V peptide from enzyme GSTP1, the I105V variants were flagged as bad quality reads from RNA-seq data but passed filters from the exome-seq data (**Table S4**). Of these combination variants, two are exome-seq only derived variants that span exon boundaries.

### Chemoproteomic identified variants are in diverse functional sites across protein families

We next asked whether the chemoproteogenomic-identified SAAVs might be of functional significance. By stratifying the the CADD scores of identified SAAVs, we find that the enrichment of high CADD score missense variants in the dMMR rare variant subset was maintained for SAAVs identified by chemoproteogenomics, including for gain-of-cysteine SAAVs (**Figure S28, S29**).

As CADD scores only provide a prediction of deleteriousness, we also asked whether any of the identified variants are located in Census genes or have been reported in Clinvar. We identify 77 variants previously reported in ClinVar (**Table S4**), with nearly all annotated as benign. A total of 16 mutations and 7 putative driver mutations (dN/dS p-values) were identified in Census genes. One prevalent driver was KRAS G12C, which was identified in several of the cell lines known to harbor this variant as a driver mutation (MIA-PACA-2 and H358 but not H2122). As KRAS expression is known to vary across cell lines^71^, this data suggests both H358 and MIA-PACA-2 cell lines are suitable for chemoproteogenomic target engagement analysis of G12C-directed compounds. However, as a cautionary example in mapping peptides, we identify several SAAV-peptides that match to multiple protein sequences, including sequences in human leukocyte antigens (HLA) and POTE ankyrin domain family proteins (**Figure 4G**). Most notably, the RHOT2 R425C mitochondrial GTPase peptides in H358 cells have exact sequence similarity to KRAS G12C peptides; these half-tryptic peptides are also identified in H1437 cells that do not harbor the KRAS G12C variant.

Chemoproteogenomics failed to capture several key Census gene SAAVs that we detected on the genomic level (e.g. SMAD4 (D351H) in CaCo-2, FBXWY (R505C) in Jurkat and CDK6 (R220C) in Molt-4 cells). Several Census gene SAAVs did, however, stand out due both to their high CADD scores and proximity to known pathogenic mutation sites. These variants of interest include MLH1 R385C, RAD17 L557R (proximal Cys551/556), MSN R180C, HIF1A S790N (proximal Cys800) and CTCF R320C, a likely pathogenic position in this protein (CADD score = 29.4) (**Figure 4H, Table S4**).

Exemplifying the utility of the chemoproteogenomics to uncover new variants, we find that 20 of the identified SAAVs have not been previously reported in COSMIC, CCLE or ClinVar (**Table S4**). One variant of unknown significance, not reported in ClinVar, is HMGB1 R110C labeled in the Molt-4 cell line (**Figure 4I**) (CADD score = 24.1). Adjacent Cys106 is a cysteine under highly controlled redox state that is responsible for inactivating the immunostimulatory state of HMGB1^100^. We also identify SARS R302H (proximal Cys300;CADD = 32), a mutation in the ATP binding site of serine-tRNA ligase, which is a tRNA ligase involved in negative regulation of VEGFA expression^101^.

Given the comparatively limited set of variants at or proximal to known damaging sites, we next broadened our analysis to include SAAVs at or proximal to UniProtKB annotated active sites (AS) and binding sites (BS) (**Figure 4J**). We find that 27 SAAVs are located within the permissive range of 10 amino acids of a known functional residue, including 4 active sites and 24 binding sites. Specific examples of high value SAAVs include tRNA synthetase EPRS R1152 (proximal Cys1148; CADD = 33), a mutation known to cause complete loss of tRNA glutamate-proline ligase activity^102^. Interestingly, EPRS has mTORC-mediated roles in regulating fat metabolism^103^. We also capture a variant proximal to the active site of BLM hydrolase I75T (proximal Cys73,78; CADD = 27.6), a cysteine protease responsible for BLM anti-tumor drug resistance^104^. More broadly, analysis of SAAV location by protein domains, reveals no marked bias for variants located in specific domain types, with the ubiquitous P-loop NTPase domain as the most SAAV-rich domain (**Figure S30, Table S4**).

As cysteines play critical roles in protein structure via disulfide bond formation together with additional cysteine oxidative modifications^105^, we asked whether identified loss of cysteine variants (10 in total) were annotated as involved in disulfides. Likely due to the comparatively small number of loss-of-cys variants, none were observed with disulfide annotations. To further pinpoint whether any variants are sensitive to oxidative modification, we subjected our previously reported Jurkat cell redox chemoproteomics datasets to reanalysis^106^. In total, our reanalysis quantified 7 acquired cysteines and 54 variants proximal to acquired cysteines. For nearly all of the cysteines quantified both in our reference database searches and now also identified with proximal variants, we observed a high concordance between variant– and reference sequence oxidation (R^2^=0.77). One notable exception was the Mitochondrial-processing peptidase enzyme (PMPCA) Cys225, for which markedly different cysteine oxidation states were measured for the reference peptide Cys (∼3% oxidation) and variant peptide Cys (∼88% oxidation) (**Figure 4K**). These data provide evidence that the proximal P226S mutation profoundly impacts Cys225 sensitivity to oxidative modifiers.

### Assessing how differential expression impacts chemoproteogenomic detection

Our comparatively modest coverage of SAAVS achieved by chemoproteogenomics (particularly when compared to our genomics datasets) is on par with the coverage reported by most prior proteogenomics studies^41,43,53^. A notable exception is the recent study by Coon and colleagues that implemented ultra-deep fractionation to achieve more global coverage of variants^44^. Inspired by this work, we next sought to ask whether chemoproteogenomics, with its built in enrichment step, would enable sampling of variants not detectable by fractionation methods (**Figure 5A**). We subjected lysates from HCT-15 and Molt-4 cells, which were chosen based on high rare missense burden, to tryptic digest, off-line high pH fractionation, and LC-MS/MS analysis. In aggregate across both cell lines, we identified 8,435 proteins and 149,006 peptides, including 1,069 unique SAAVs found in 1,352 total peptides using our 2-stage MSFragger search (**Figure 5B**,**S31**,**Table S5**). 26 peptides were identified that contained multiple variants, including peptides that would only be detected by our combinatorial databases (**Figure 4B**) as well as those readily detected by combined ‘Single-Each’ and ‘All-in-One’ database searches (**Table S5**).

**Figure 5.**
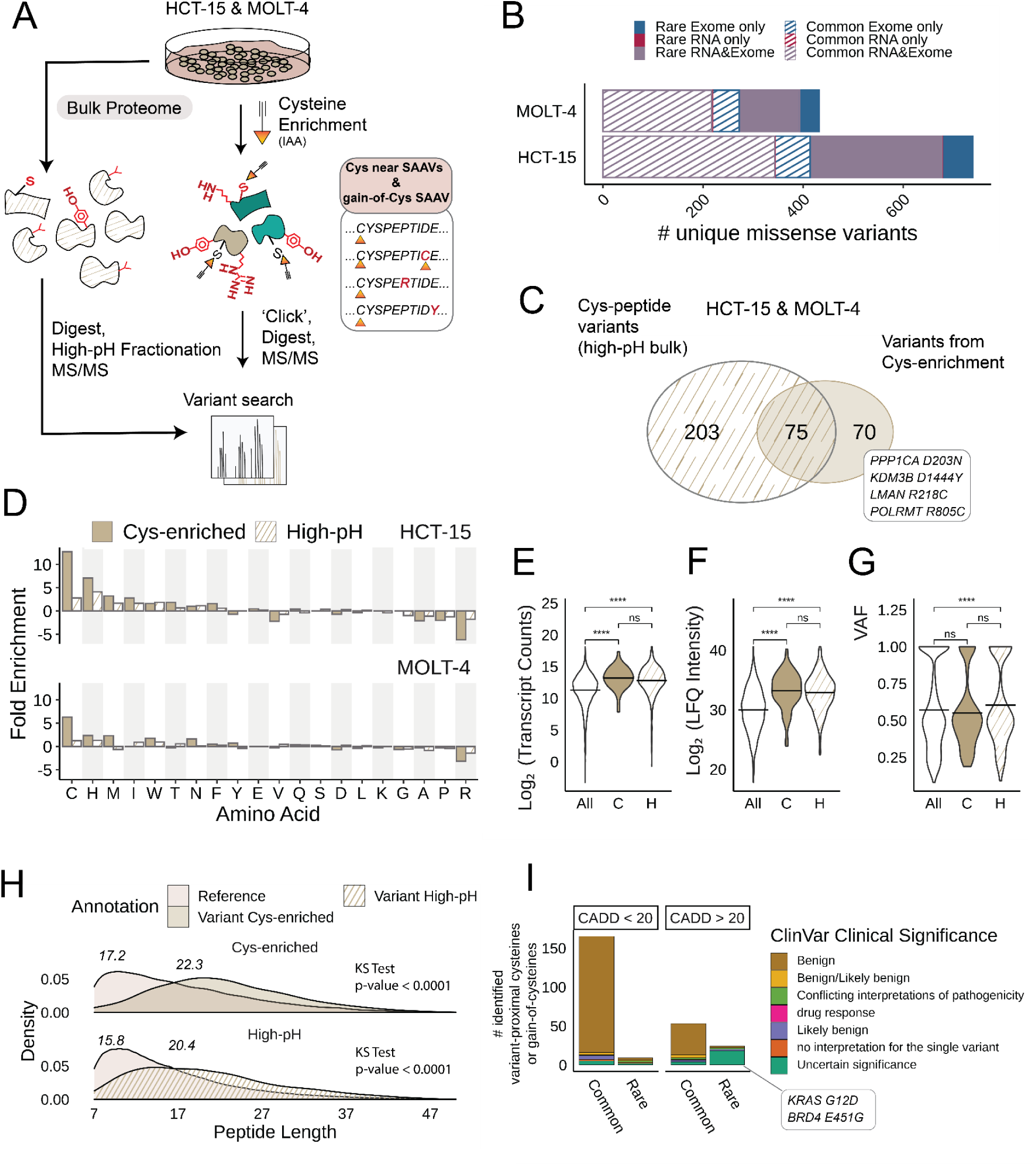
Comparison of variants identified from cysteine enrichment and bulk proteomics. A) Workflow for high-pH fractionation of lysates. Cell lysates are treated with DTT and iodoacetamide followed by digestion, high-pH fractionation, and LC-MS/MS analysis. Triplicate high-pH sets for HCT-15 and Molt-4 cells were used. B) Total numbers of unique missense variants identified from either RNA-seq or WE-seq or both after using 2-stage MSFragger search of high-pH datasets. C) Overlap of cysteine-containing peptide variants identified from bulk fractionation and cysteine enrichment datasets. D) Fold enrichment of amino acids as a ratio of the net amino acid frequency (gain minus loss) to the amino acid frequency in all missense-containing proteins detected in high-pH and cys-enriched datasets. E) DE-seq normalized transcript counts for all RNA variants ‘All’, variants detected from cys-enrichment ‘C’, and variants detected from high-pH fractionation ‘H’ in HCT-15 cells. F) Label free quantitation (LFQ) intensities for proteins matched to all RNA variants ‘All’, variants detected from cys-enrichment ‘C’, and variants detected from high-pH fractionation ‘H’ in HCT-15 cells. G) Variant allele frequencies (VAF) (total reads/total coverage per site) for RNA-seq variants called in HCT-15 and Molt-4 cells. E-G statistical significance was calculated using Mann-Whitney U test, **** p < 0.0001, ns p > 0.05. H) Peptide lengths of reference and variant peptides identified in dataset types. I) High-pH detected variants stratified by CADD score and ClinVar clinical significance. Data found in **Table S5**.

Comparison of this unenriched dataset to the chemoproteogenomic dataset for the matched HCT-15 and Molt-4 proteomes (145 total SAAVs identified by chemoproteogenomics for these two cell lines) revealed 70 SAAVs, including eight acquired cysteines, uniquely identified with chemoproteogenomics (**Table S4-S5**), (**Figure 5C).** Despite the lower numbers of total SAAVs in the chemoproteogenomics datasets, we find that chemoproteomic enrichment afforded a ∼5-fold boost in the relative fraction of acquired cysteines captured (**Figure 5D**). Further stratification of the net detected amino acid changes (**Figure S32-S33**) revealed that, again, cysteine was a top gainer and arginine was the most lost amino acid for both enriched and unenriched datasets.

We next asked whether protein or RNA abundance might rationalize the differences in SAAV coverage for each method. Comparison of normalized transcript counts for SAAV-matched genes identified either by chemoproteogenomics or in our bulk proteomic dataset, for HCT-15 cells analysis revealed no significant difference between measured transcript abundance between the sets (**Figure 5E, Table S5**). A notable subset of SAAVs (3,262 total, including PIK3CA E545K, TP53 S241F, SMARCA4 R885C TCGA hotspot mutations) with low abundance transcripts (less than 4000 normalized counts) were not detected in either the chemoproteogenomics or bulk proteogenomics. Providing further evidence that lower transcript abundance decreases the likelihood of detection, we find that an even more sizable fraction of cysteines found in reference protein sequences matched with low abundance genes are not detected, both for high-pH fractionated samples and chemoproteomics enriched samples (**Figure S34**).

Given the likely disconnect between transcript abundance and protein abundance^110–112^ for some SAAVs analyzed, we also extended these analyses to measures of protein abundance. Using label-free quantification (LFQ) analysis, we find that for proteins with proteomic-detectable SAAV peptides, the quantified protein intensities were significantly higher when compared to proteins for which the corresponding variants were only detected via genomic analysis. No difference was observed between the bulk fractionated samples and the chemoproteogenomic samples (**Figure 5F, Table S5**).

As both the transcript and protein abundance analyses do not delineate reference from variant-specific transcript/protein sequences, we also compared the variant allele frequencies (VAF) for SAAVs detected by each method. We find that high-pH variant allele frequencies (VAF) were significantly higher than the chemoproteogenomic detected SAAVs, which were comparable to the aggregate bulk RNA-seq VAFs (**Figure 5G, Table S5**). This enrichment for lower VAF for the chemoproteogenomic detected SAAVs extended to the acquired cysteine subset (**Figure S34**).

Given that cysteine chemoproteomics requires peptide derivatization, with a comparatively large (463 Da) biotin modification, we postulated that some differences in coverage might be ascribed to behavior of peptides during sample acquisition. Comparing the properties of the SAAV peptides detected by chemoproteogenomics versus proteogenomics we observed a more restricted charge state distribution for cysteine-enriched samples and no appreciable differences in the amino acid content beyond enrichment for cysteine (**Figure S35**). While we did not observe differences in the peptide lengths in our comparison of between the chemoproteomic-enriched and high pH detected SAAV peptides, a marked significant increase in SAAV peptide length (average 5AA longer) was observed compared to reference peptides in both datasets (**Figure 5H**). This increased peptide length is consistent with the ubiquity of loss-of-arginine SAAVs in both datasets, which are favored in the longer length peptides (**Figure S36**).

Protein families analysis revealed slight differences between the two datasets with enzymes making up a larger fraction of cys-enriched detected variant proteins. Significantly higher CADD scores were also observed for enrichment data (**Figure S37**). Notable high-CADD score variants identified only from enrichment include lysine demethylase KDM3B D1444Y, RNA polymerase POLRMT R805C, glycoprotein transporter LMAN2 R218C and Serine/threonine-protein phosphatase PP1-alpha catalytic subunit PPP1CA D203N (**Figure 5C**). Addition of the bulk proteomic analysis yielded coverage of 85 notable variants belonging to Census genes, including BRD4 E451G and KRAS G13D, and 26 rare and common variants of uncertain significance in ClinVar, including rare gain-of-cysteines ubiquitin hydrolase USP8 Y1040C and LMNA R298C (**Figure 5I**,**Table S5**).

### Chemoproteogenomics enables ligandability screening

As demonstrated by our previous studies, cysteine chemoproteomics platforms are capable of pinpointing small-molecule targetable cysteine residues^21,22,26,29^. Therefore, we next paired our 2-stage search method with cysteine-reactive small molecule ligandability analysis to establish a chemoproteogenomic small molecule screening platform (**Figure 6A**). We first opted to use the widely employed scout fragment **KB02**^29^ (**Figure 6B**) to compare the ligandable variant proteomes for three high variant burden dMMR cell lines (HCT-15, Jurkat, and Molt-4). For **KB02** treated samples, we identified 210 total variants. The high concordance for ratios detected for variant peptides with multiple alleles provides evidence of the robustness of our platform and hints that most cysteine proximal variants do not substantially alter cysteine ligandability (**Figure 6C)**.

**Figure 6.**
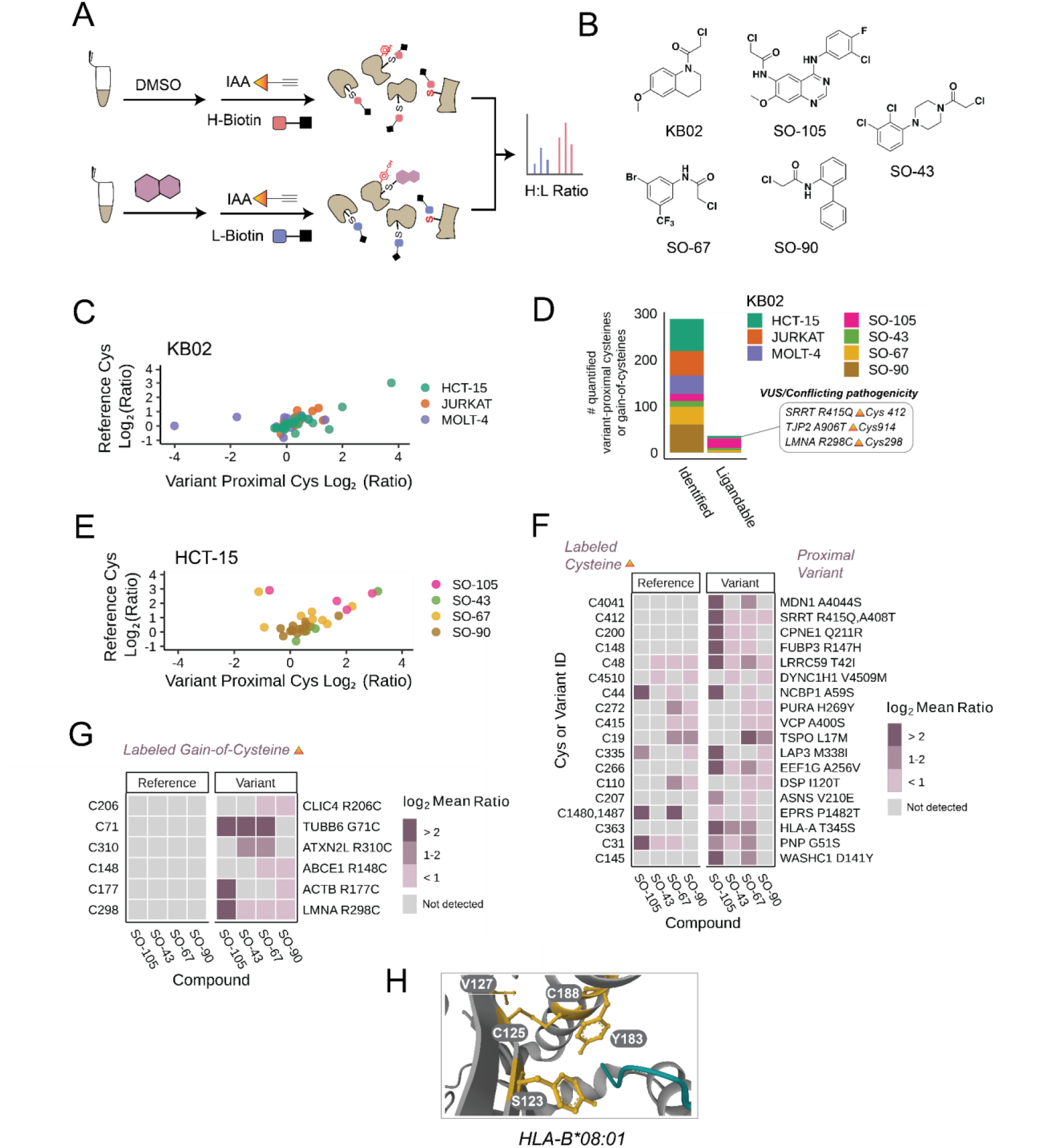
Assessing ligandability of variant proximal cysteines and gain-of-cysteines. A) Schematic of activity-based screening of Cys reactive compounds; cell lysates are labeled with compound or DMSO followed by chase with IAA and ‘click’ conjugation to heavy or light biotin click conjugation to our isotopically differentiated heavy and light biotin-azide reagents, tryptic digest, LC-MS/MS acquisition, and MSFragger analysis. B) Chloroacetamide compound library. C) Total quantified variants and total ligandable variants (Log_2_ Ratio > 2) identified stratified by cell line (KB02 data) or compound (HCT-15 cell line). D) Correlation of high-confidence variant containing and reference cysteine ratio values from KB02 data. E) Correlation of high-confidence variant containing and reference cysteine ratio values from SO compound data. F) Log_2_ heavy to light ratio values for variant containing and reference cysteine peptides. G) Subset of gain of cysteine peptide variant log_2_ ratios. H) Crystal structure of HLA-B*08:01 protein liganded Cys125, disulfide Cys188, and binding site residue Y183 as well as variant sites V127 and S123 (PDB: 3X13). Data provided in **Table S6**.

We next subjected the HCT-15 proteome to more in depth analysis using a small panel of custom electrophilic fragments (**Figure 6B**). We observed 27 total liganded variant peptides in 27 proteins in the HCT-15 proteome, which are labeled by one or more compounds (**Figure 6C**). As with the **KB02** cell line comparison, nearly all multi-allelic peptides showed comparable ratios (**Figure 6E**). Nucleotide analogue **SO-105** was observed to be more promiscuously reactive (**Figure 6F**) when compared to the less elaborate fragments.

In aggregate across all ligandability datasets, we identified 259 total variants found in 232 total proteins (**Figure 6D**). Of these variants, 57 were acquired cysteines, in 55 proteins; 22 were ligandable (Log2(HL) ratio > 2), variant-proximal cysteines and 10 were ligandable gain-of-cysteines (**Figure 6D**). Notable liganded sites we identify include Cullin-associated NEDD8-dissociated protein 1 (CAND1) G1069C–a site which mutated in the Arabadopisis ortholog reduces auxin response^113^ and Tubulin beta 6 (TUBB6) G71C (**Figure 6G**). Some sites with differing reference and variant ratios include EPRS P1482T–the mutated proline nearby Cys 1480 may be requisite for labeling by electrophilic fragments. We also identify 3 ligandable variants of uncertain significance or conflicting pathogenicity that we show may be modulated for study with small molecules and could act as potential starting points for biological analyses (**Figure 6C**). As multi-allelic acquired cysteine sites cannot be captured sans cysteine, no analogous ratio comparison could be performed for the 6 total quantified acquired cysteines (**Figure 6G**).

To understand functionality of the ligandable variant sites in 3D protein space, we analyzed active site and binding sites within 10 angstrom distance of the ligandable Cys residues and Cys-proxial variant sites (**Table S6**). We find three ligandable cysteines near or in active/binding sites including previously identified HMGB1 Cys106 (R110C) (**Figure 4I**), as well as Aldolase A ALDOA Cys178 (G196G) and HLA-B/C Cys125 (V127L/S123Y). Intriguingly HLA-B/C Cys125 (C101 post signal peptide cleavage), near peptide binding region sites Y183 is liganded by **KB02** in HCT-15 cells which harbor HLA-B*08:01 and HLA-B*35:01 (**Figure 6F**). This conserved cysteine plays important roles in HLA structure^114^. Ligandability of this site is unexpected as this site is known to be disulfided with C188 in cell surface HLA^115^; however, we find in our sequencing that HCT-15 cells harbor truncated beta-2-microglobulin (β2m) protein (B2M Y30*) (**Table S2**). β2m is known to stabilize this specific disulfide^115,116^, facilitating protein folding and translocation to the cell surface^117–119^. In HLA-B27 allelic variants, Cys125 is known to be exposed without β2m^120^.

### Expanding HLA cysteine peptide coverage and gel-based ABPP of HLA covalent labeling

Major Histocompatibility Complex (MHC) Class I molecules (known as HLA molecules in humans) present intracellularly derived protein fragments, either self-derived or from pathogens in the context of cross-presentation, on the cell surface for recognition by T cells and subsequent immune response; noncovalent assembly of a polymorphic heavy chain with a light chain (β2m) and peptide occurs in the endoplasmic reticulum (ER) followed by translocation via the Golgi to the cell surface^121^. Recent reports of allele-specific HLA-binding compounds, most notably abacivir HIV drug^122^, together with efforts to develop covalent modulators of MHC Class I and II complexes^123–125^ prompted us to assess the impact of chemoproteogenomics on achieving improved coverage of highly polymorphic genes (**Figure 7A**). 15,000 HLA alleles have been reported in the human population^126^. Exemplifying this impact on proteomic sequence coverage, our panel of cell lines alone harbor >25 HLA-A, B and C alleles (**Table S2**), while most protein reference databases only contain one copy of each MHC Class I and Class II molecule.

**Figure 7.**
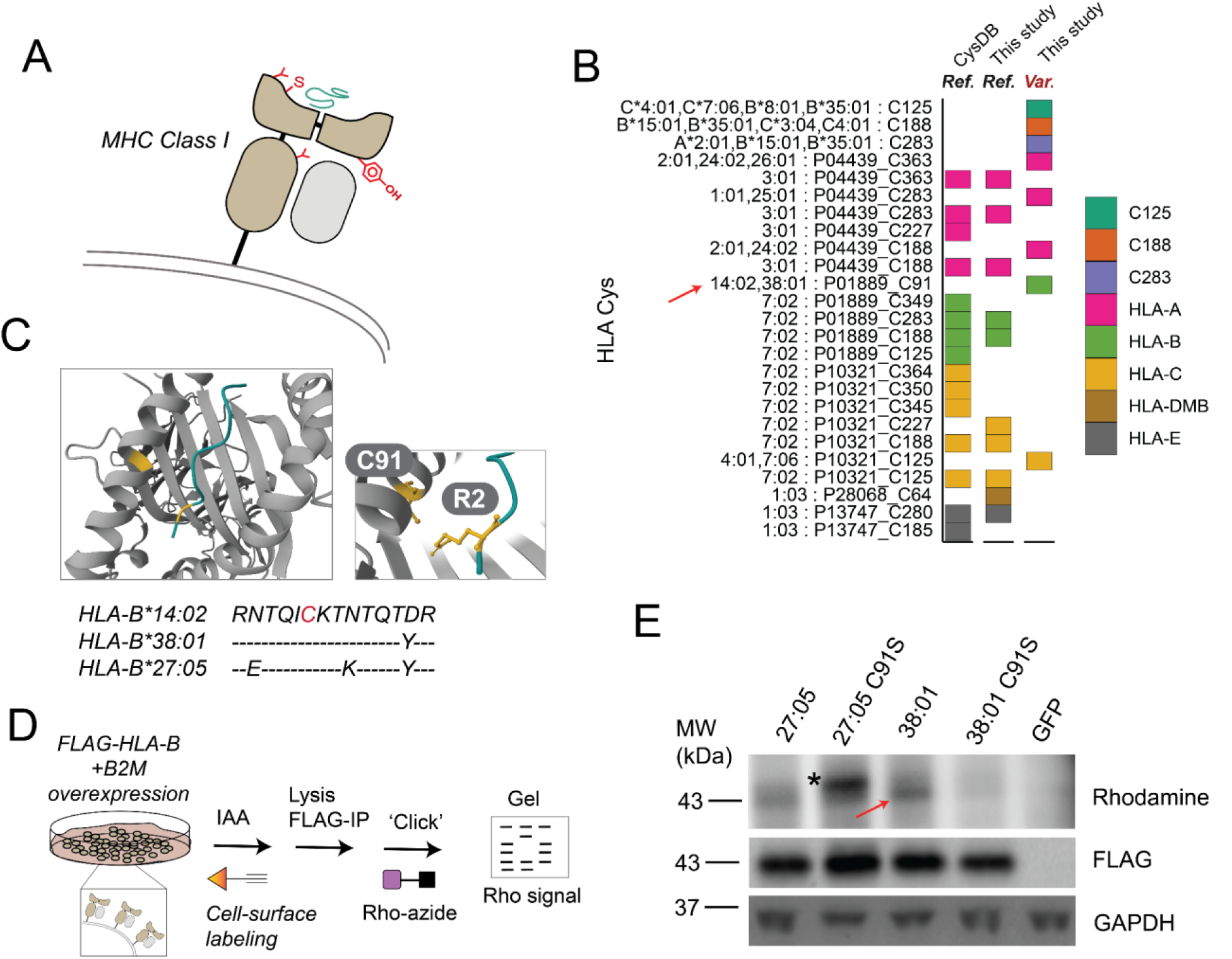
Expanding HLA cysteine peptide coverage and gel-based ABPP of HLA covalent labeling. A) Schematic of highly variable HLA binding pocket containing cysteine with bound peptide. B) Coverage of HLA cysteines from this study and in CysDB; color indicates HLA type or multi-mapped cysteines. C) Crystal structure of HLA-B 14:02 (PDB: 3BXN) with highlighted Cys67 and Arg P2 position of bound peptide; alignments of Cys91 regions of three HLA-B alleles. D) Workflow to visualize HLA cysteine labeling; first cells were harvested and treated with IAA followed by lysis, FLAG immunoprecipitation, and click onto rhodamine-azide. E) Cys-dependent cell surface labeling of HLA-B alleles with IAA, band indicated with red arrow and non-specific band represented with asterisk (representative of 2 two biological replicates). Data provided in **Table S7**.

Through search of sample-specific databases of both chemoproteomics and high pH fractionated samples, we achieved ∼50% more coverage of HLA-A sequence in comparison to reference searches (**Figure 7B and Figure S39**). A key finding of our analysis was detection of HLA-B Y91C (C67 post signal peptide cleavage), which lies in the extracellular peptide binding pocket of HLA-B and was identified as IAA-labeled in MeWo cells (**Figure 4J**). The MeWo cell line HLA alleles (HLA-B*14:02 and HLA-B*38:01) both harbor this comparatively rare Cys (**Figure 7C**). Notably this cysteine is also a key feature of the pathogenic ankylosing spondylitis associated allele HLA-B*27^127,128^. To test whether this cysteine was amenable to gel-based ABPP analysis and to determine whether this IAA labeling extends to HLA-B*27:05, we co-expressed c-terminal FLAG tagged HLA-B*38:01, HLA-B*27:05, HLA-B*38:01 C91S, and HLA-B*27:05 C91S with beta-2-microglobulin (β2m) and subjected cells to in situ IAA labeling followed by lysis, FLAG immunoprecipitation to enhance the detectability of the HLA cysteine, and click conjugation to rhodamine azide (**Figure 7D**). Gratifyingly, we observed a Cys67-specific rhodamine signal (**Figure 7E**), showcasing the utility of gel-based ABPP in visualizing HLA small molecule interactions. Notably IAA labeling was also observed for HLA-B27:05, although the presence of a strong co-migrating band in the HLA-B27:05 C67S immunoprecipitated sample complicates interpretation of the specificity of this labeling to Cys67. We were unable to observe comparable signal in lysate-based labeling studies, supporting enhanced accessibility of this cysteine to cell-based labeling (**Figure S40**).

### FragPipe graphical user interface with improved 2-stage MSFragger search and FDR estimation

Motivated by the multi-faceted uses of the 2-stage search pipeline, including those reported here and future envisioned applications, we also sought to facilitate the utilization of the 2-stage search strategy by the scientific community. Therefore, we enhanced FragPipe by establishing semi-automated execution of these searches while also providing an option to run MSBooster and Percolator (instead of PeptideProphet) to further improve the sensitivity of identification of variant peptides **(Figure 8A**).

**Figure 8.**
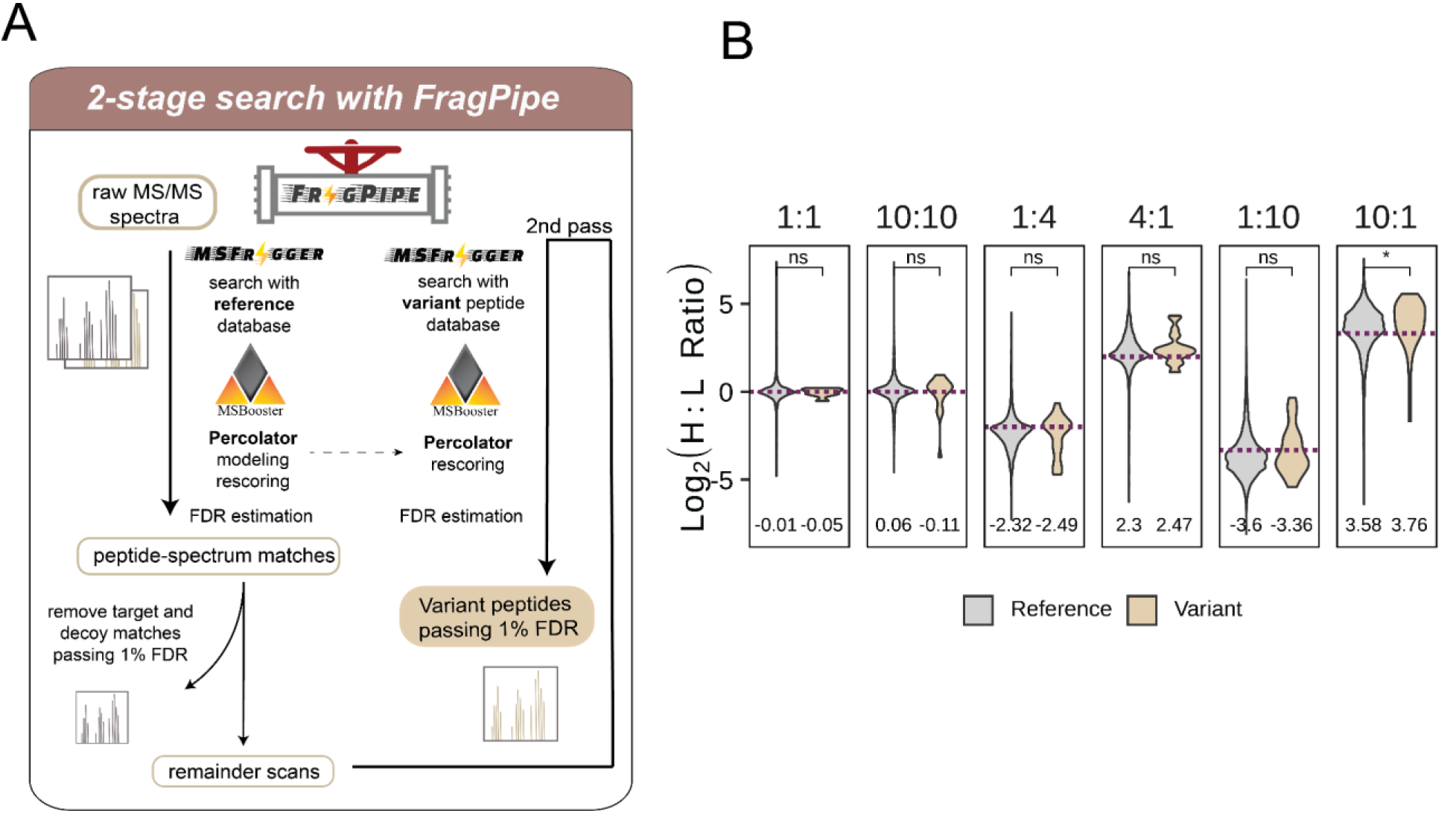
2-stage search implemented into FragPipe GUI with Percolator rescoring. A) 2-stage search incorporation into FragPipe GUI workflow. B) Heavy to light ratios (H:L) from triplicate datasets comparing identifications from reference and variant searches; mean ratio value indicated, *dashed lines* indicate ground-truth log2 ratio, statistical significance was calculated using Mann-Whitney U test, * p < 0.05, ** p < 0.01, ns p > 0.05. Data provided in **Table S8**.

In the first stage pass, with the “write sub mzML” option enabled, FragPipe utilizes MSFragger^85,129^ for mass calibration, search parameter optimization, and database searching. Following this, FragPipe applies MSBooster^130^ to compute the deep-learning scores^130^, Percolator^131^ for PSM rescoring, ProteinProphet^132^ for protein inference, and Philosopher for FDR filtering. Subsequently, FragPipe generates new mzML files, which include the scans that did not pass the FDR filtering (default is 1%) and those with a probability higher than a predefined threshold (default is 0).

In the second search, as the mass spectral files have already been calibrated and only scans that remained unidentified in the first search have been retained, the mass calibration should be disabled. Moreover, Percolator modeling might fail in the second pass due to a lack of sufficient number of high-scoring PSMs. Therefore, FragPipe lets Percolator reuse the model from the initial pass. FragPipe then generates a new workflow file containing optimized parameters, and a new manifest file with the new (subset) mzML files specified for the second-pass search. The user is merely required to load these two files without needing any further adjustments.

Using the new GUI features, we observe comparable coverage for both the command-line and automated GUI implementations of the 2-stage search with a slight increase in numbers of identifications observed for datasets processed with MSBooster and Percolator (**Figure S41,Table S8**). The ratio differences between variant and reference Cys peptide are comparable (**Figure 8B**).

## DISCUSSION

SAAvs are a ubiquitous feature of human proteins, which remain under sampled in established proteomics pipelines. Here, we merged genomics with mass spectrometry-based chemoproteomics to establish chemoproteogenomics as an integrated platform tailored to capture and functionally assess the missense variant cysteinome. Our chemoproteogenomics study is distinguished by a number of features including: (1) genomic stratification of the predicted pathogenicity of acquired cysteine residues, (2) cell-line paired custom combinatorial search databases, (3) FragPipe enabled 2-stage database search platform ensuring class-specific FDR estimation, and (4) capacity to pinpoint both redox-sensitive and ligandable genetic variants proteome-wide. To facilitate widespread adoption of our approach, including for applications beyond the study of the variant cysteinome, the user-friendly GUI-based FragPipe platform now features a robust semi-automated version of our 2-stage search (**Figure 8**).

To build chemoproteogenomics, we started by analyzing publically available datasets in Clinvar, COSMIC, and dbSNP, which revealed that cysteine acquisition is a ubiquitous feature of human genetic variation, which predominates in the context of DNA damage repair responses. The instability of CpG motifs is a key driver of bulk cysteine acquisition, which occurs largely hand-in-hand with bulk arginine depletion, across both cancer genomes and healthy genomes and rare and common variants. Many colon cancer cell lines and other MSI high cell lines are particularly enriched for cysteine acquisition—however, nearly all of the acquired residues in these lines are not driver mutations, which complicates their use as models for assessing the potentially druggability of variants with established clinical connections and highlights the value of future efforts to analyze additional missense variant rich cell lines and perform CRISPR-Cas9 base editing to engineer variants of interest into endogenous loci^35,133–136^.

Armed with a set of variant rich cell lines, we next generated combinatorial SAAV-peptide databases for cell-line specific SAAVs as identified in cell-line matched whole exome and transcriptome datasets. In total, across 11 cell lines sequenced, we identified 1,453 missense variants, of which 116 led to gain-of-cysteine. Looking towards future iterations of chemoproteogenomics, we expect that the use of tumor-normal paired variant calling with tools such as MuTect2^137^ will further decrease the likelihood of false discovery introduced by factors such as cell heterogeneity and low read quality—for cell lines that lack matched normal controls, we expect that the pairing of publically available datasets (e.g. DepMap, https://depmap.org/) with custom sequencing data, will prove another useful strategy to further bolster the quality and accessibility of variant-containing databases. Such multi-pronged approaches will likely prove most useful when paired with combinatorial custom databases, such as the peptide-based databases reported here, which were designed to minimize increased search space complexity while also more fully accounting for cell heterogeneity.

By building upon prior reports describing 2-stage database searches for class-specific FDR control^53–55^ as a rigorous search strategy that reduces the likelihood that a false positive variant peptide detection, here we deployed a 2-stage search approach in FragPipe, first as a custom command-line workflow and subsequently as a user-friendly semi-automated workflow in the FragPipe GUI. Enabled by our previously reported isotopically enriched heavy– and light-biotin-azide capture reagents^87^, we provide compelling evidence to support the low rates of false discovery of variant peptides using the 2-stage search—spurious false discovery of variant peptides would easily be detected from MS1 precursor ion ratios that deviate from the expected spike-in values **(Figure 3,8**). Our isotopic labeling strategy also enabled the assessment of the ligandability and redox sensitivity of variant peptides. Our discovery of a cysteine in PMPCA that exhibits variant-dependent changes in oxidation provides an intriguing anecdotal example that supports the future utility of chemoproteogenomics in more broadly characterizing the missense variant redox proteome. Given the critical role that disulfides play in protein structure and folding and the causal roles for cysteine mutations in human disease, for example the NOTCH mutations that cause the neurodegenerative disorder CADASIL^138^, we expect a subset of these lost cysteines could be implicated in altered protein abundance or activity. Through cysteine chemoproteomic capture, we identified ligandable variant-proximal cysteines in Census genes such as RAD17, including one gain-of-cysteine of uncertain significance in LMNA (R298C). Other liganded cysteines proximal to variants of uncertain significance include TJP2 (A906R) and SRRT (R415Q). Demonstrating the utility of our approach, we identified a Cys91 (Cys67) as labeled by IAA both by proteomics and gel-based ABPP. As this cysteine is shared with the pathogenic HLA-B27, it is exciting to speculate about the impact of covalent modification on HLA peptide presentation. Our application of chemoproteogenomics to screening of a focused library of electrophilic compounds, identified 32 ligandable variant-proximal Cys which demonstrates that cysteine ligandability can be assessed proteome-wide in a proteoform-specific manner.

Looking beyond our current study, we anticipate multiple high value applications for chemoproteogenomics. Application to immunopeptidomics should uncover additional covalent neoantigen sites, analogous to the recent reports for Gly12Cys KRAS^124,139^. Pairing of chemoproteogenomics with ultra-deep offline fractionation should further increase coverage and allow delineation of variants that alter protein stability, including the numerous high CADD score acquired cysteines, which we find were underrepresented in our proteomics analysis when compared to genomic identification. Inclusion of genetic variants beyond SAAVs will allow for capture of additional therapeutically relevant targets that result from indels, alternative splicing^39,140^, translocations, transversions, or even undiscovered open reading frames such as microproteins^141,142^. Thus chemoproteogenomics is poised to guide discovery of proteoform-directed therapeutics.

## Acknowledgments

We thank all members of the Backus lab for helpful suggestions. We thank the UCLA Technology Center for Genomics and Bioinformatics (TCGB). Additionally, we thank Jigar Desai for guidance on NGS data processing, Angela Wei for guidance on Kallisto data processing, and Ian Ford for providing a CuAAC-compatible IP protocol. The results here are in part based upon data generated by the COSMIC-CLP: https://cancer.sanger.ac.uk/cell_lines and TCGA Research Network: https://www.cancer.gov/tcga. This study was supported by a Beckman Young Investigator Award (K. M. B.), V Scholar Award V2019-017 (K. M. B.), UCLA Jonsson Comprehensive Cancer Center Seed Grant (K. M. B.), and the National Institutes of Health grants R01-GM094231 and U24-CA271037 (A. I. N.). The content is solely the responsibility of the authors and does not necessarily represent the official views of the National Institutes of Health.

## Author Contributions

H. S. D., K. M. B., and A.I.N. conceptualization; H. S. D. formal analysis; H. S. D. visualization; H. S. D. validation; H. S. D., L.M.B., F. Y., K.M.B data curation; H. S. D., S. O., and M. V. investigation; H. S. D., F. Y., and N.U. methodology; H. S. D. and K. M. B writing–original draft; H. S. D., S.O., L.M.B., F.Y., A. I. N., and K. M. B. writing–review and editing; A. I. N. and K. M. B. supervision; A. I. N. and K. M. B. funding acquisition.

## Methods

Experimental details and Tables S1-S9 can be found in the Supporting Information.

