## Supplementary Information for "Multi-omic stratification of the missense variant cysteinome"

### Contents

(A) Supplementary Figures 3

(B) Methods 43

(C) Running 2-stage searches with FragPipe  62

(D) References 64

### (A) Supplementary Figures

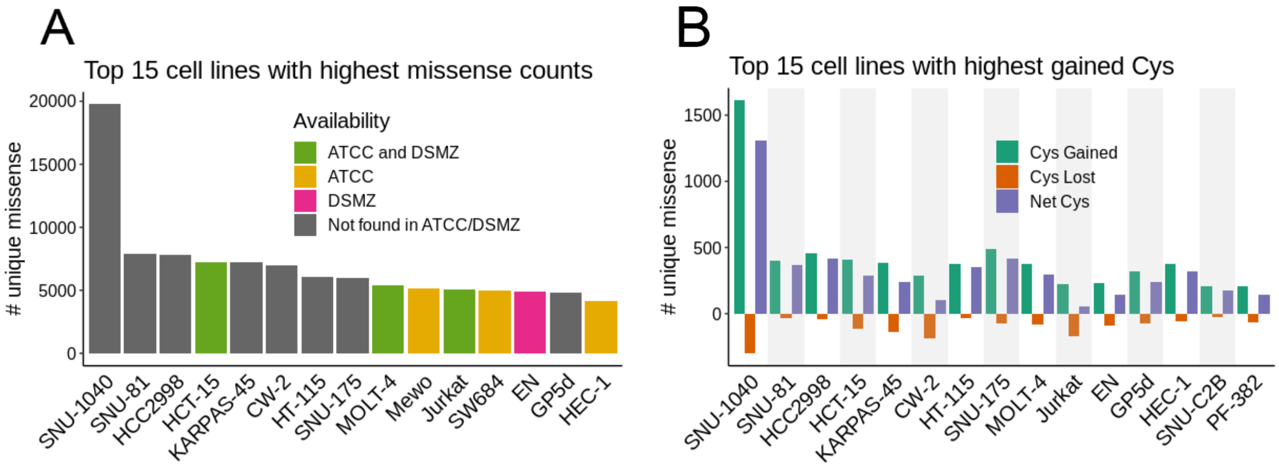


**Figure S1**. Missense counts in subsets of COSMIC Cell Lines A) Top 15 cell lines with highest missense burden in COSMIC-CLP (COSMIC Cell Lines Project release v96) color indicates availability of high mutational burden cell lines in American Type Culture Collection (ATTC) and German Collection of Microorganisms and Cell Cultures (Deutsche Sammlung von Mikroorganismen und Zellkulturen, DSMZ). B) Top 15 cell lines with highest gained cysteines showing net gain, total gained and total lost.

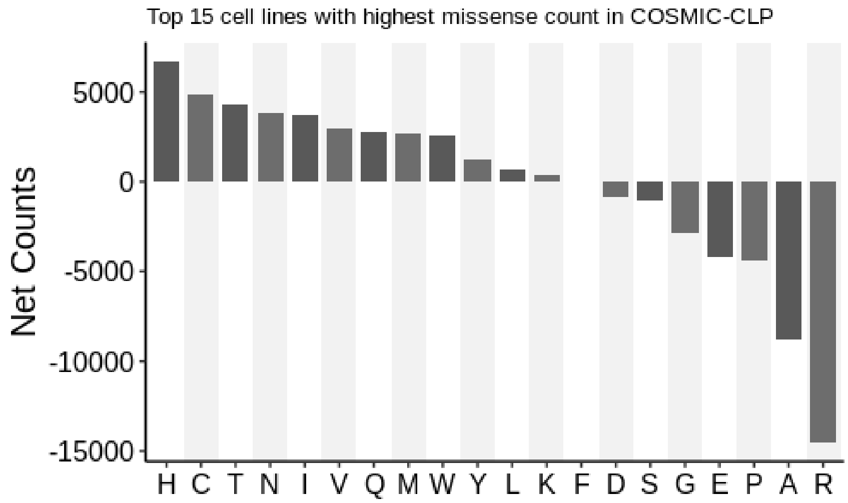


**Figure S2**. Aggregated net amino acid changes (gained counts - lost counts) in combined top 15 cell lines with highest missense burden in COSMIC-CLP (COSMIC Cell Lines Project release v96).

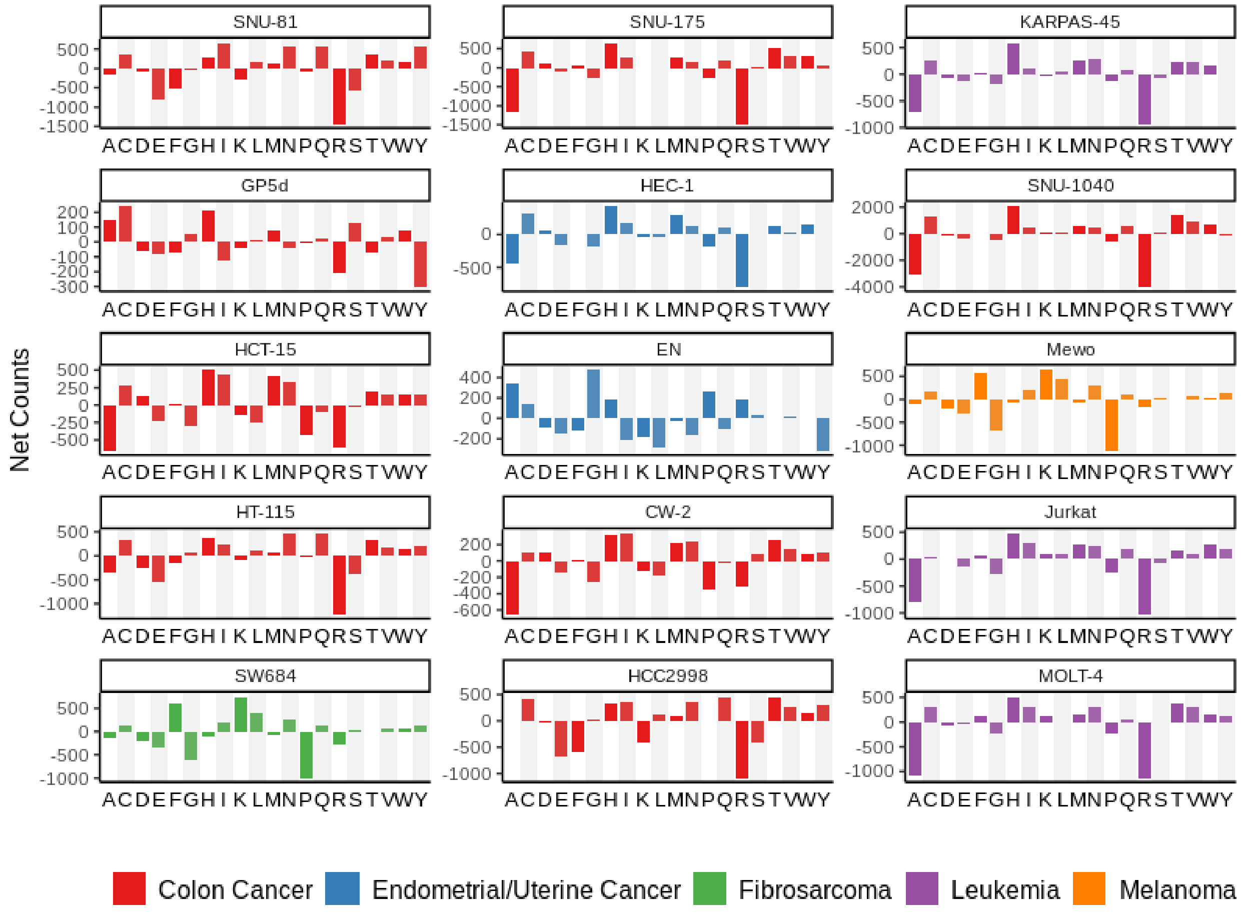


**Figure S3.** Net single amino acid mutation counts (gained counts - lost counts) in top 15 cell lines with highest missense burden (COSMIC Cell Lines Project release v96). Several cell lines show marked depletion of alanine, for example SNU-175, KARPAS-45, Jurkat (A3 subclone) and MOLT-4 cell lines, which represent two colon cancer cell lines, and two T lymphoblast cell lines, respectively. In contrast, several cell lines showed marked net gain of alanine, including EN, Gp5d, and HCC2998 cells, which are an endometrial/uterine cancer lines and two colon cancer cell lines, respectively.

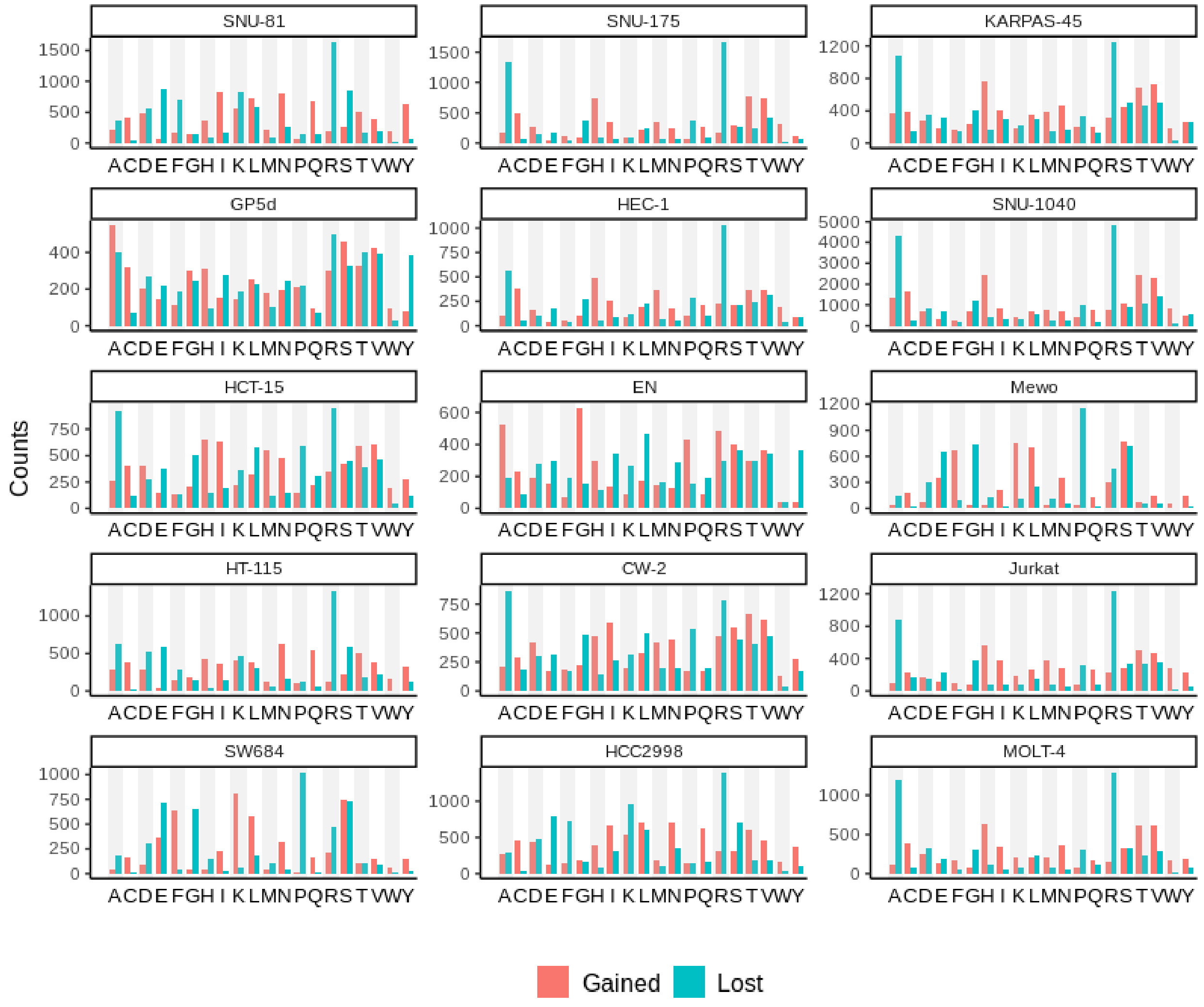


**Figure S4**. Total gain and loss missense mutation counts in the top 15 cell lines with highest missense burden (COSMIC CLP v96).

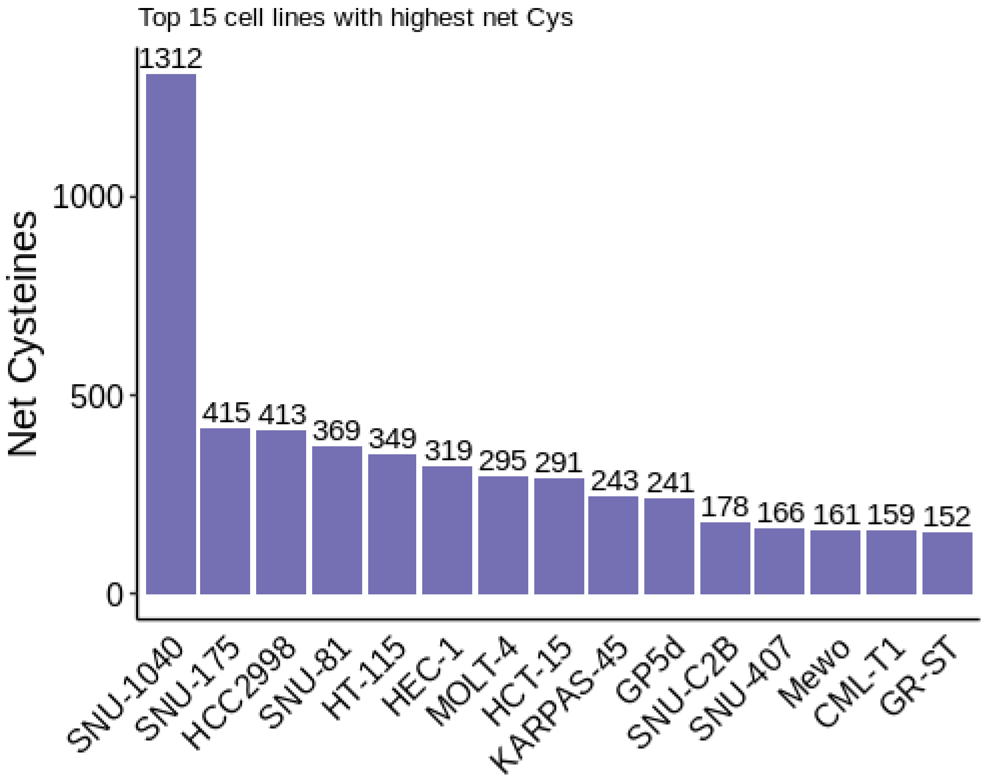


**Figure S5**. Cell lines with highest net gained cysteines (COSMIC CLP v96).

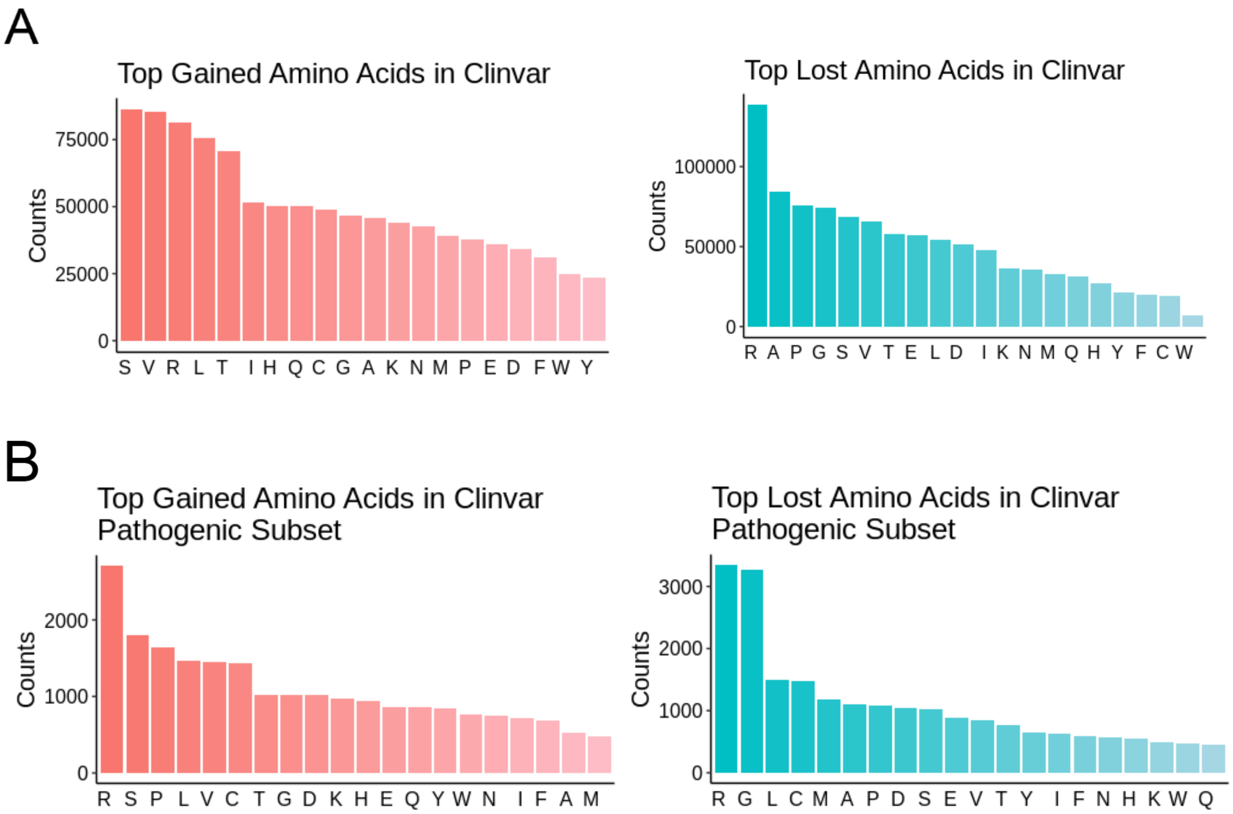


**Figure S6.** ClinVar amino acid changes A) Total gained and lost amino acids reported in all ClinVar data for unique gene name, protein position and amino acid change B) Total gained and lost amino acids for pathogenic missense variants in ClinVar. Red indicates gain amino acids and blue indicates lost amino acids

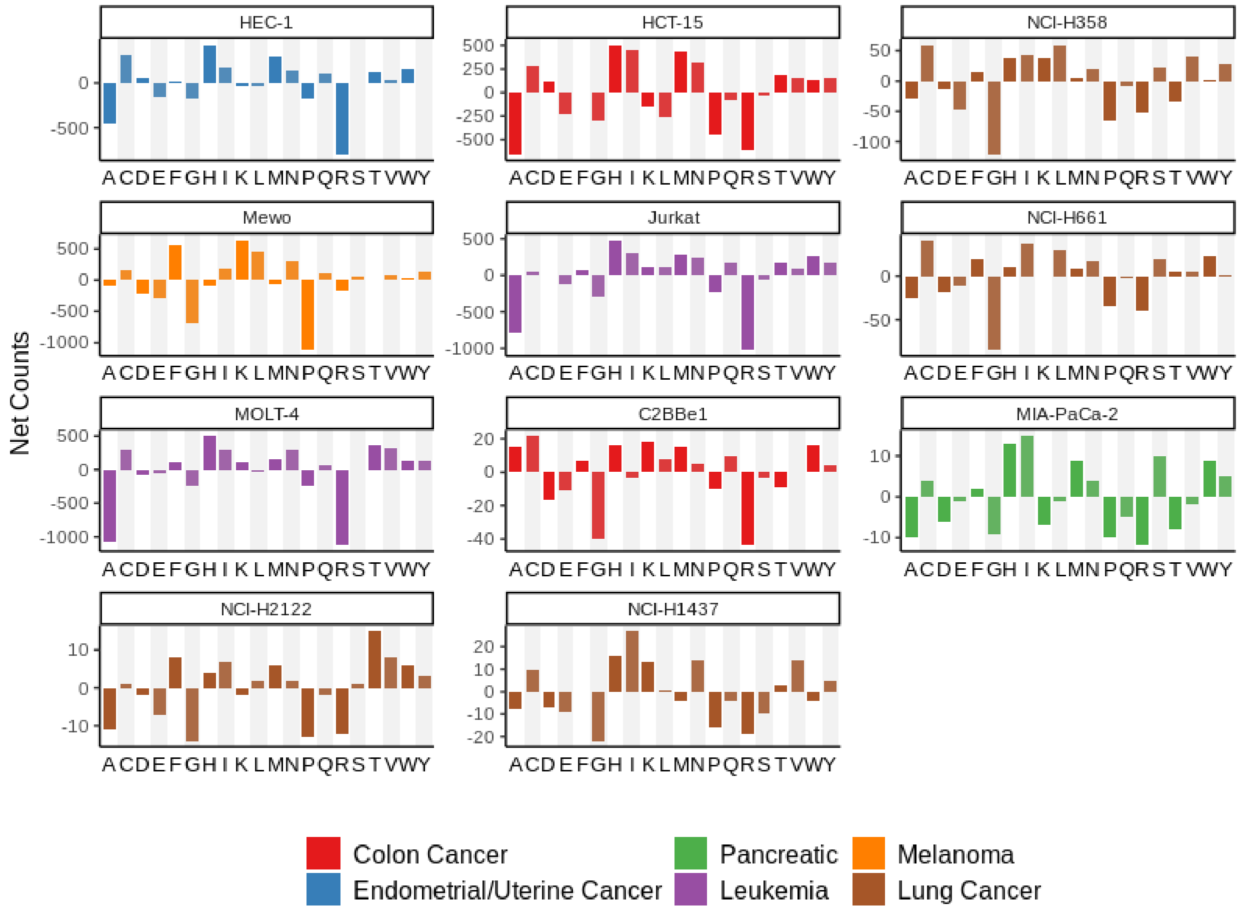


**Figure S7**. Net single amino acid mutation counts (gained counts - lost counts) in the panel of cell lines in this study (COSMIC CLP v96). C2BBe1 is a CaCo-2 cell line clone in COSMIC-CLP.

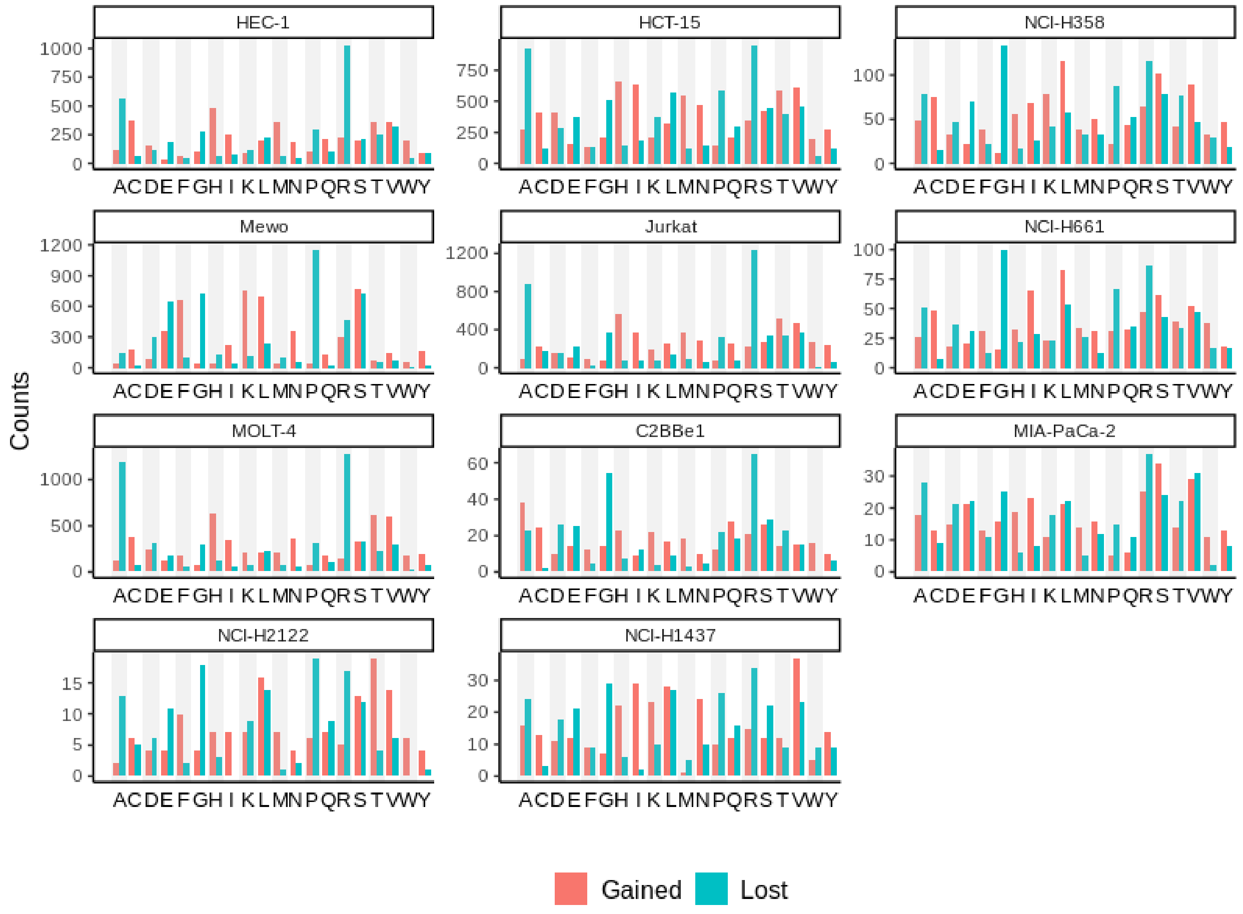


**Figure S8**. Total gain and loss missense mutation counts in a panel of cell lines in this study (COSMIC CLP v96). C2BBe1 is a CaCo-2 cell line clone in COSMIC-CLP.

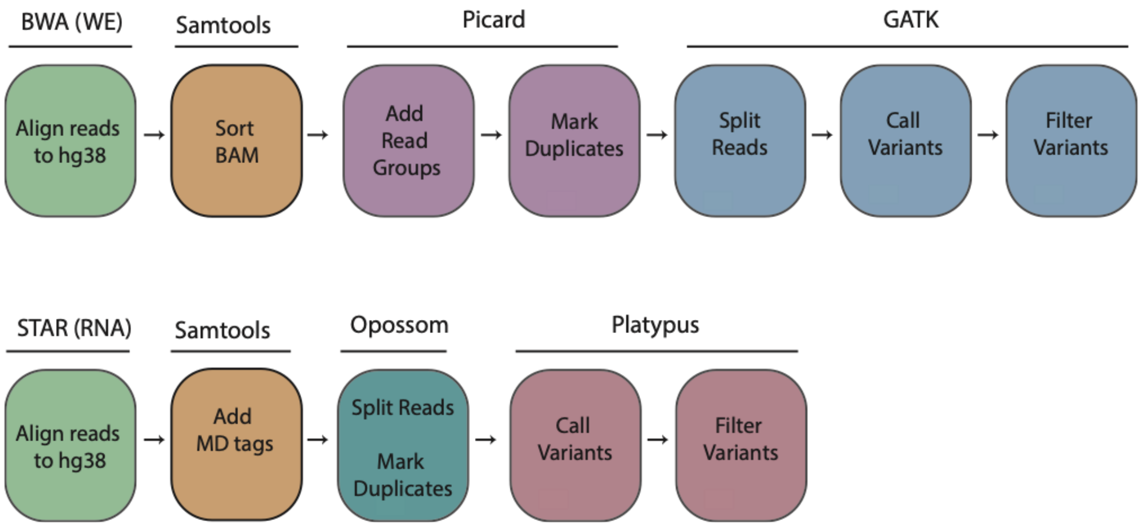


**Figure S9**. Variant calling pipelines for RNA and whole-exome (WE) datasets. Details in methods. Raw reads submitted to Sequence Read Archive (SRA) as BioProject PRJNA997729. Color indicates commands under toolkit listed.

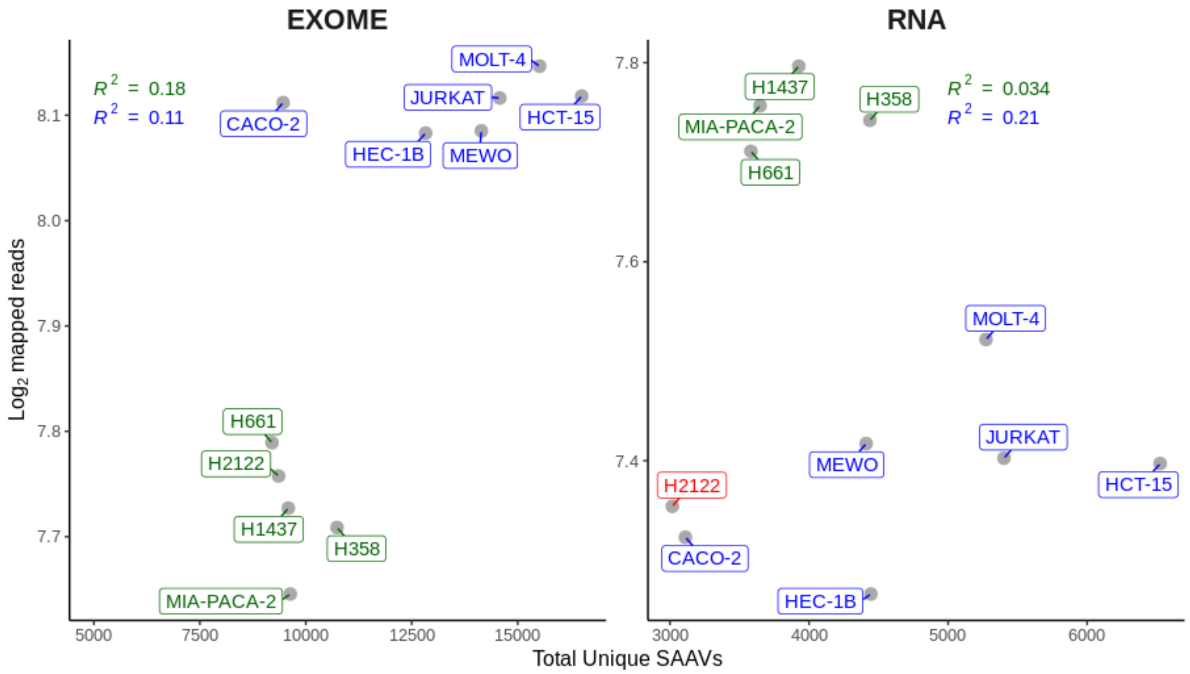


**Figure S10.** Number of mapped reads per cell line after BWA or STAR mapping and total unique SAAVs (single amino acid variants) identified per cell line. Color indicates sequencing batch

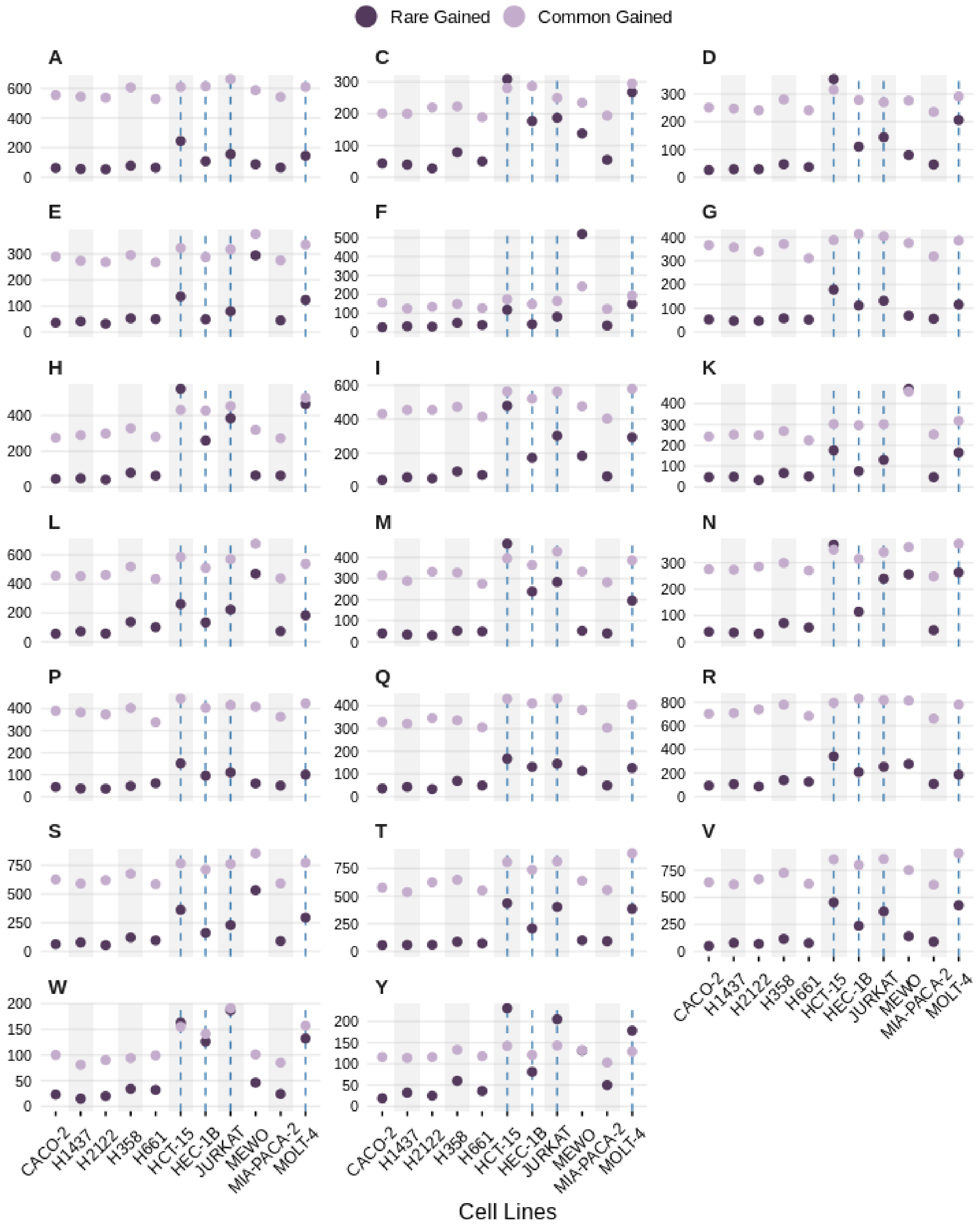


**Figure S11.** Gained amino acid counts identified in individual cell lines separated by common and rare. Dotted lines indicate dMMR cell lines

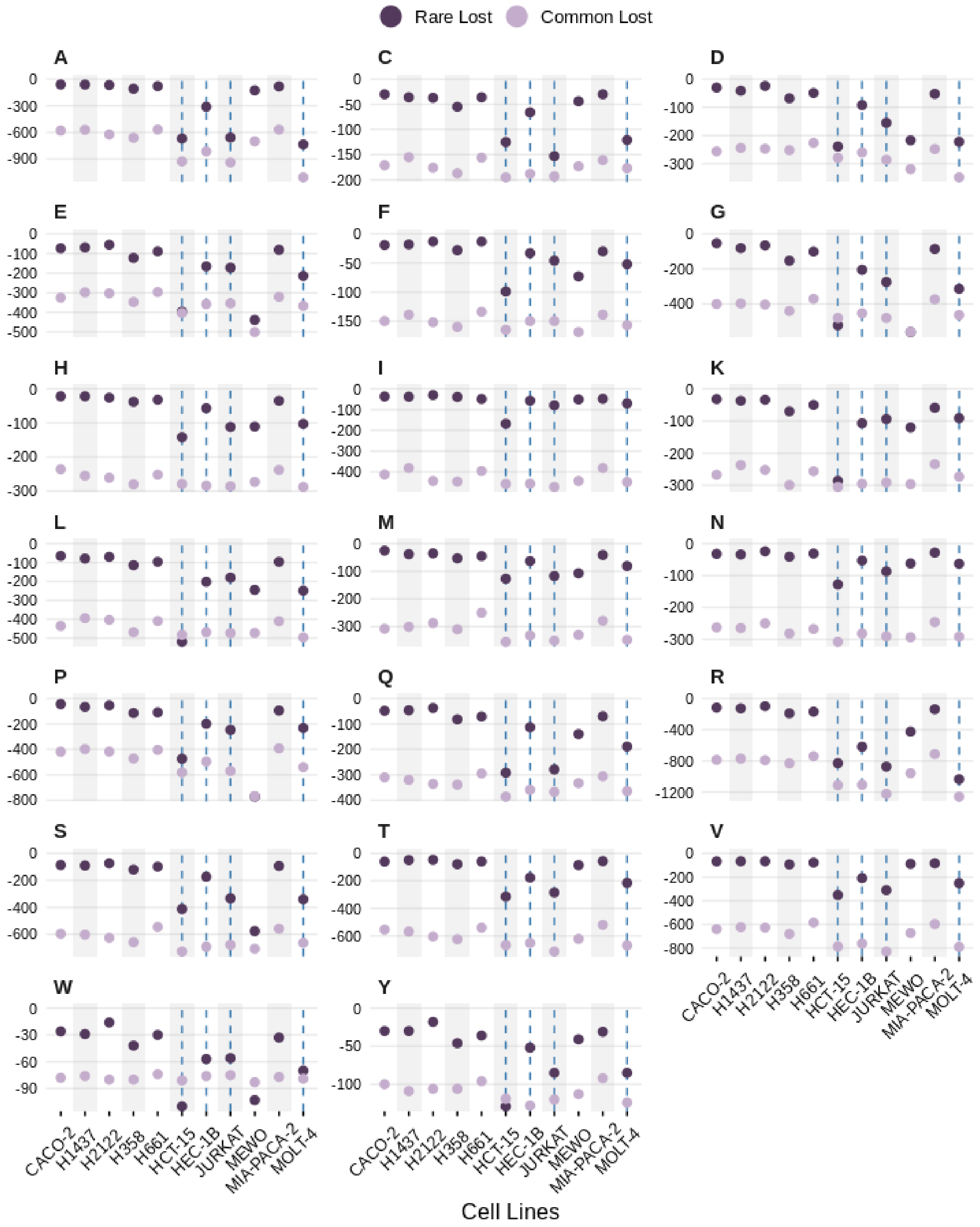


**Figure S12.** Lost amino acid counts identified in individual cell lines separated by common and rare. Dotted lines indicate dMMR cell lines.

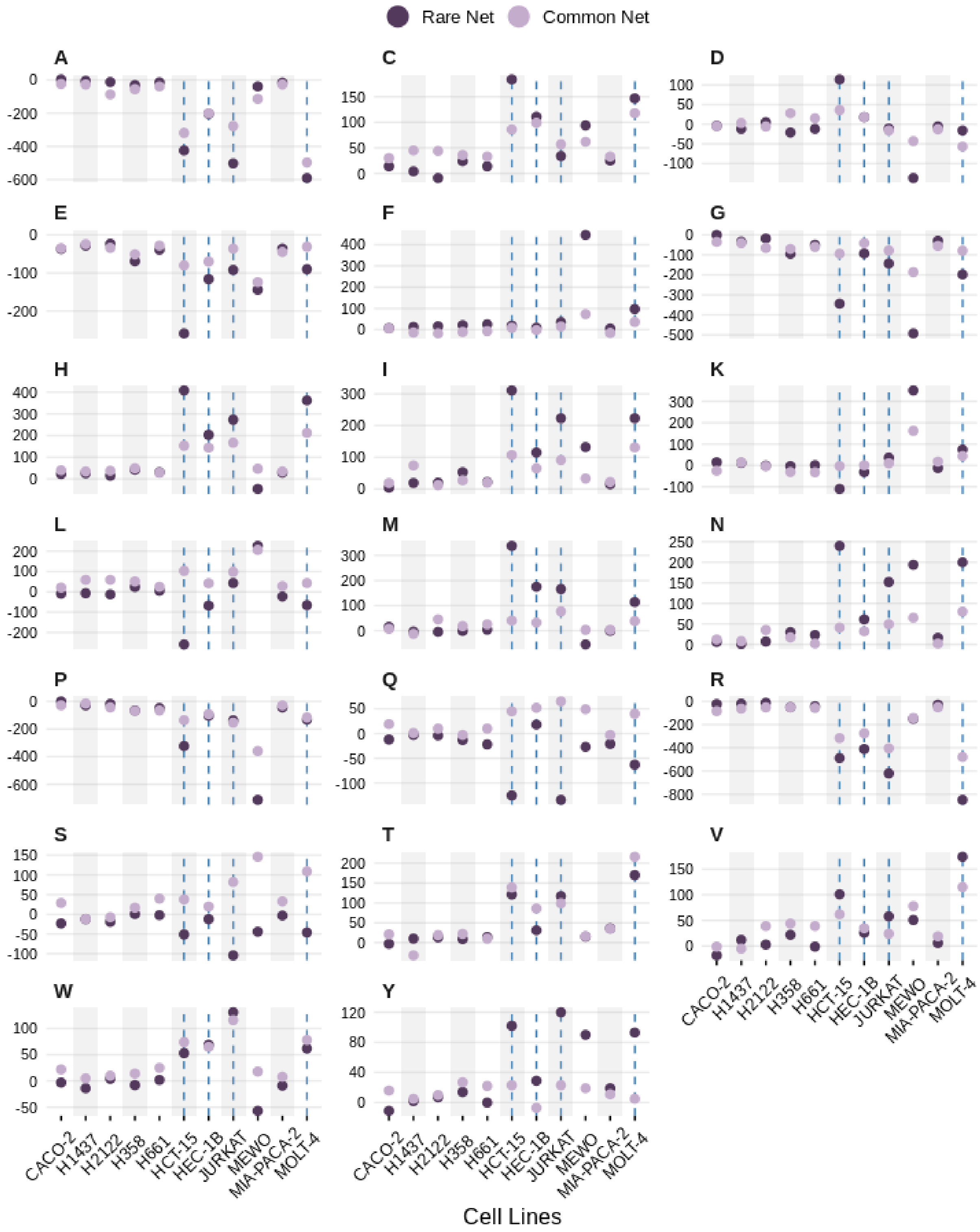


**Figure S13.** Net amino acid counts in identified individual cell lines separated by common and rare. Dotted lines indicate dMMR cell lines.

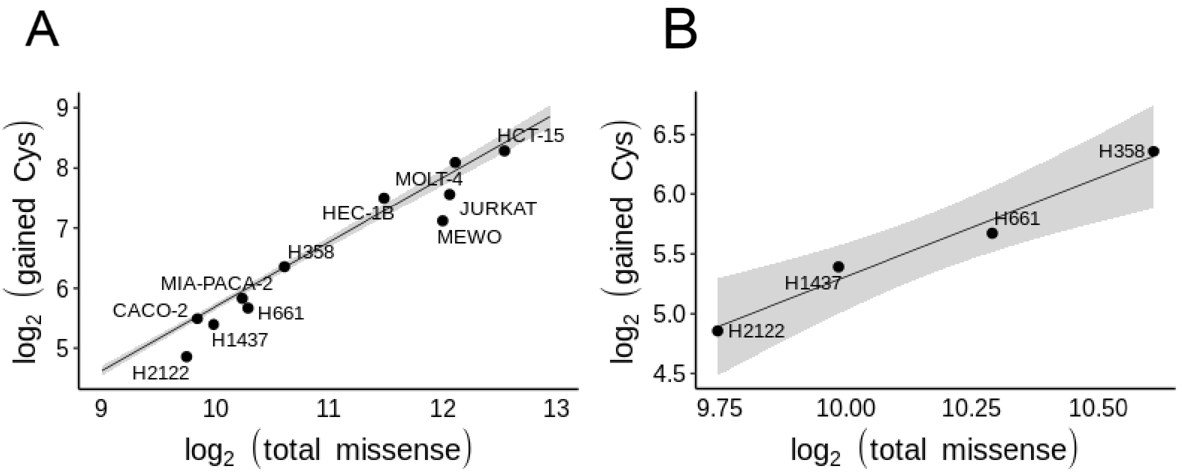


**Figure S14.** A) Gained cysteine missense mutations relative to total missense mutations from sequencing data fit to all COSMIC-CLP linear model from Figure 1B and 1C. B) Lung cancer lines subset with linear regression and 95% confidence interval shaded in gray–suggests the comparatively modest impact of smoking on cysteine acquisition for the cell lines evaluated.

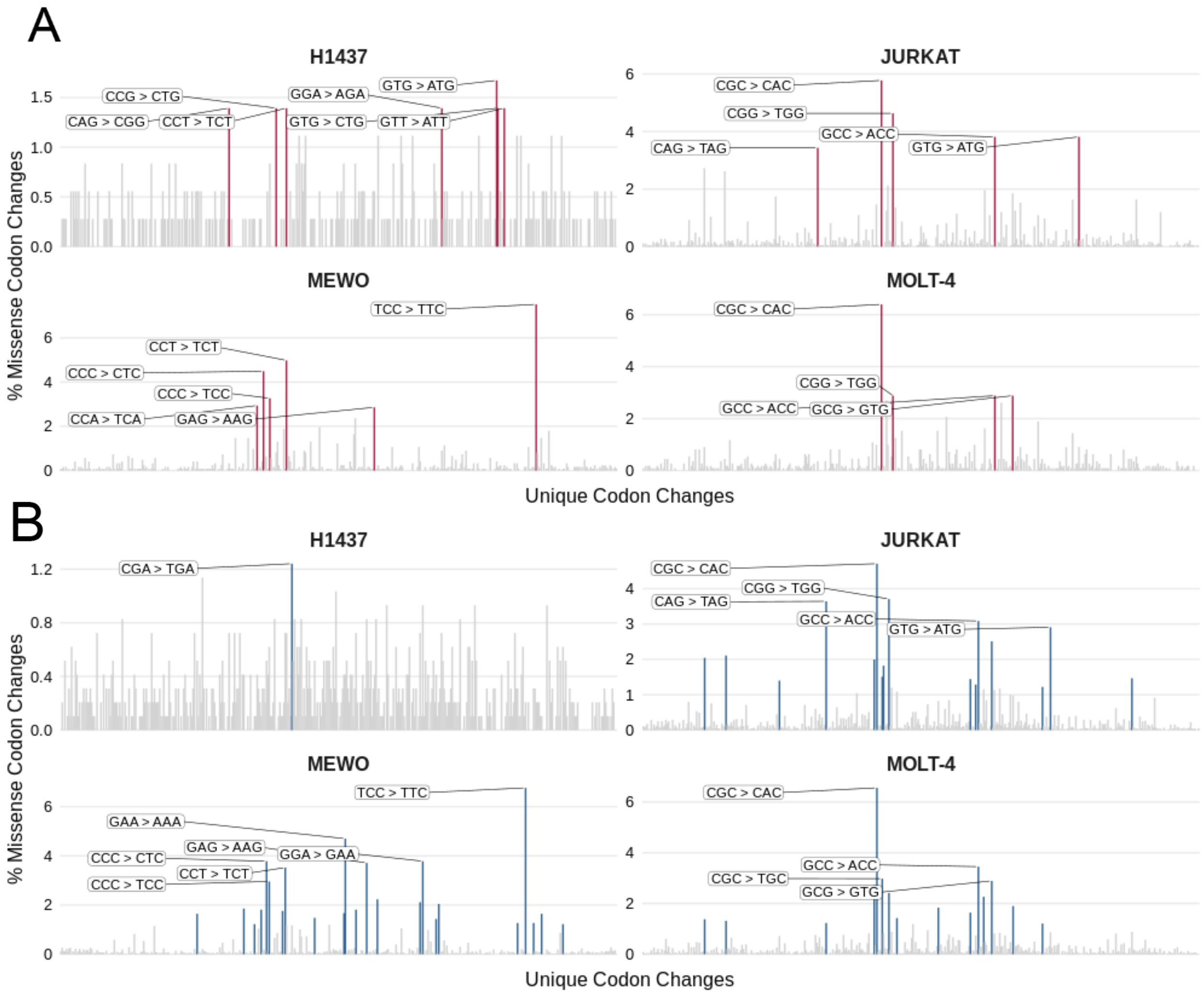


**Figure S15**. Unique codon changes in Jurkat, NCI-H1437, MOLT-4 and MeWo cell lines; RNA changes in red and exome changes in blue; analysis of rare variants. Tobacco-smoke carcinogens, such as benzo[*a*]pyrene (B[*a*]P) diol epoxides, uniquely confer G→ T (C→ A) transversions that we see enriched only in our smoking LC line. dMMR and UV mutations are predominantly C→T (G→A) transitions, the flanking nucleotides differ as UV radiation induces pyrimidine dimers which largely result in CC→TT transitions (and counterpart GG→AA) which cause gain of F/K and loss of S/E.

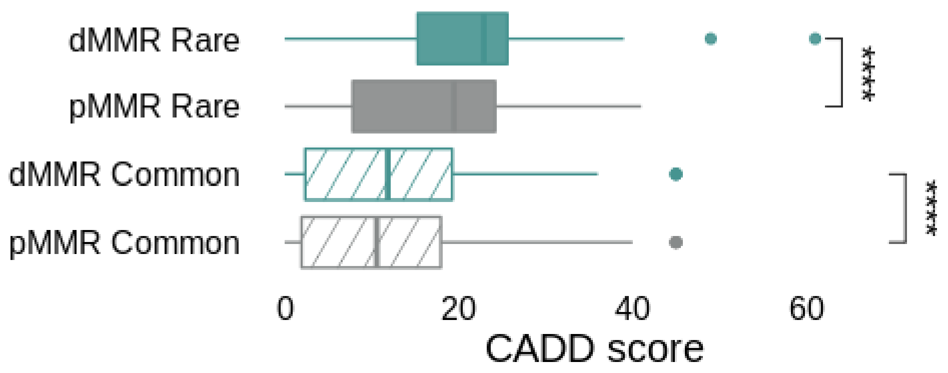


**Figure S16**. Distribution of CADD (Combined Annotation Dependent Depletion) scores for indicated variant grouping from Figure 2E data, Statistical significance was calculated using Mann-Whitney U test, **** p < 0.0001.

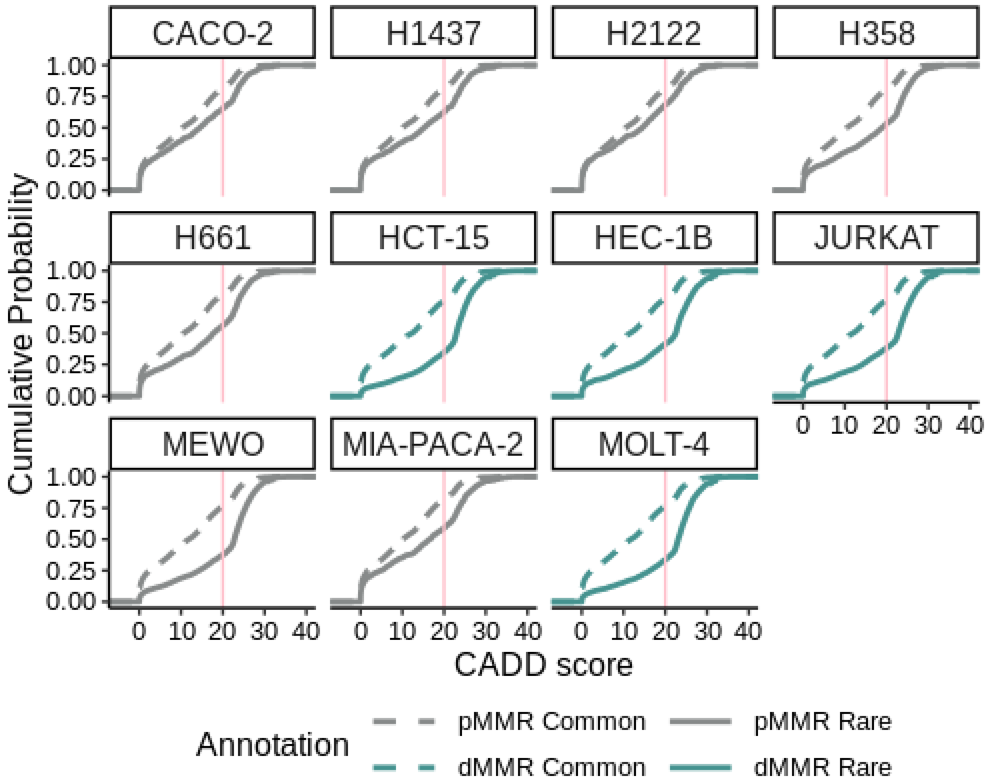


**Figure S17**. CADD-phred cumulative probabilities stratified by cell line. Pink lines indicate CADD-phred = 20

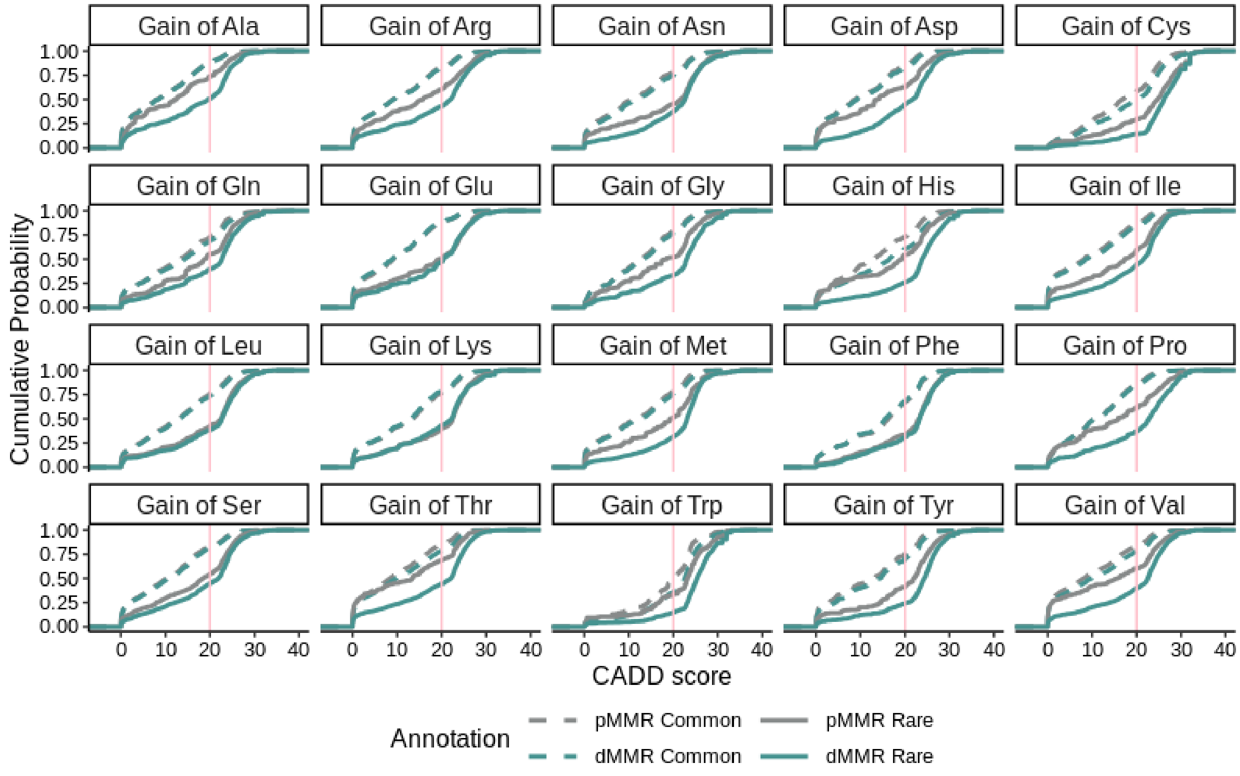


**Figure S18:** CADD-phred cumulative probabilities stratified by gained amino acid. Pink lines indicate CADD-phred = 20. One sided Mann Whitney U test p-values are in **Table S2** (tab S16).

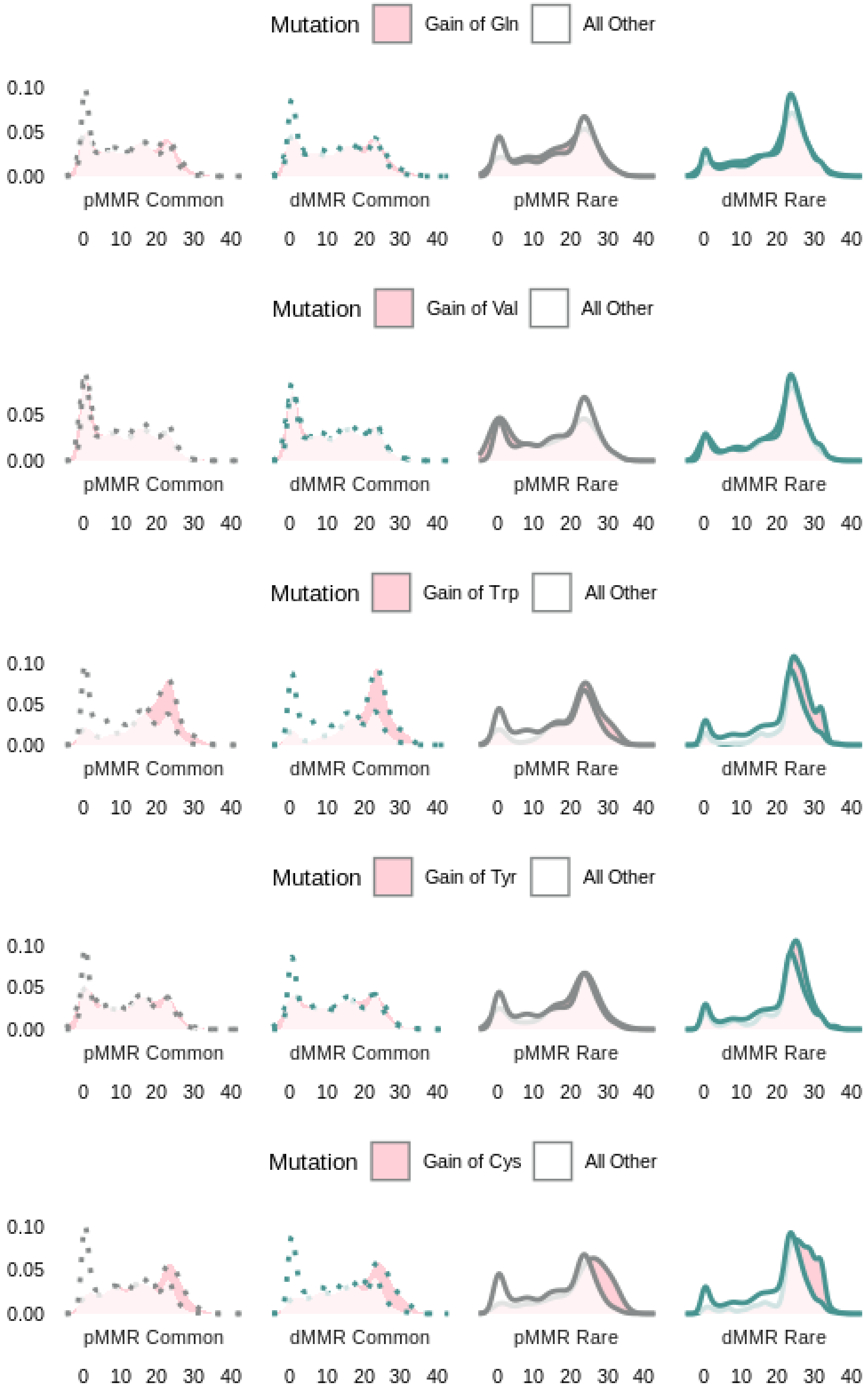
Mutation
Gain of Gln
All Other
0.10
0.05
0.00
pMMR Common
dMMR Common
pMMR Rare
dMMR Rare
0 10 20 30 40
Mutation
Gain of Val
All Other
0.05
0.00
pMMR Common
dMMR Common
pMMR Rare
dMMR Rare
0 10 20 30 40
Mutation
Gain of Trp
All Other
0.10
0.05
0.00
pMMR Common
dMMR Common
pMMR Rare
dMMR Rare
0 10 20 30 40
Mutation
Gain of Tyr
All Other
0.10
0.05
0.00
pMMR Common
dMMR Common
pMMR Rare
dMMR Rare
0 10 20 30 40
Mutation
Gain of Cys
All Other
0.10
0.05
0.00
pMMR Common
dMMR Common
pMMR Rare
dMMR Rare
0 10 20 30 40

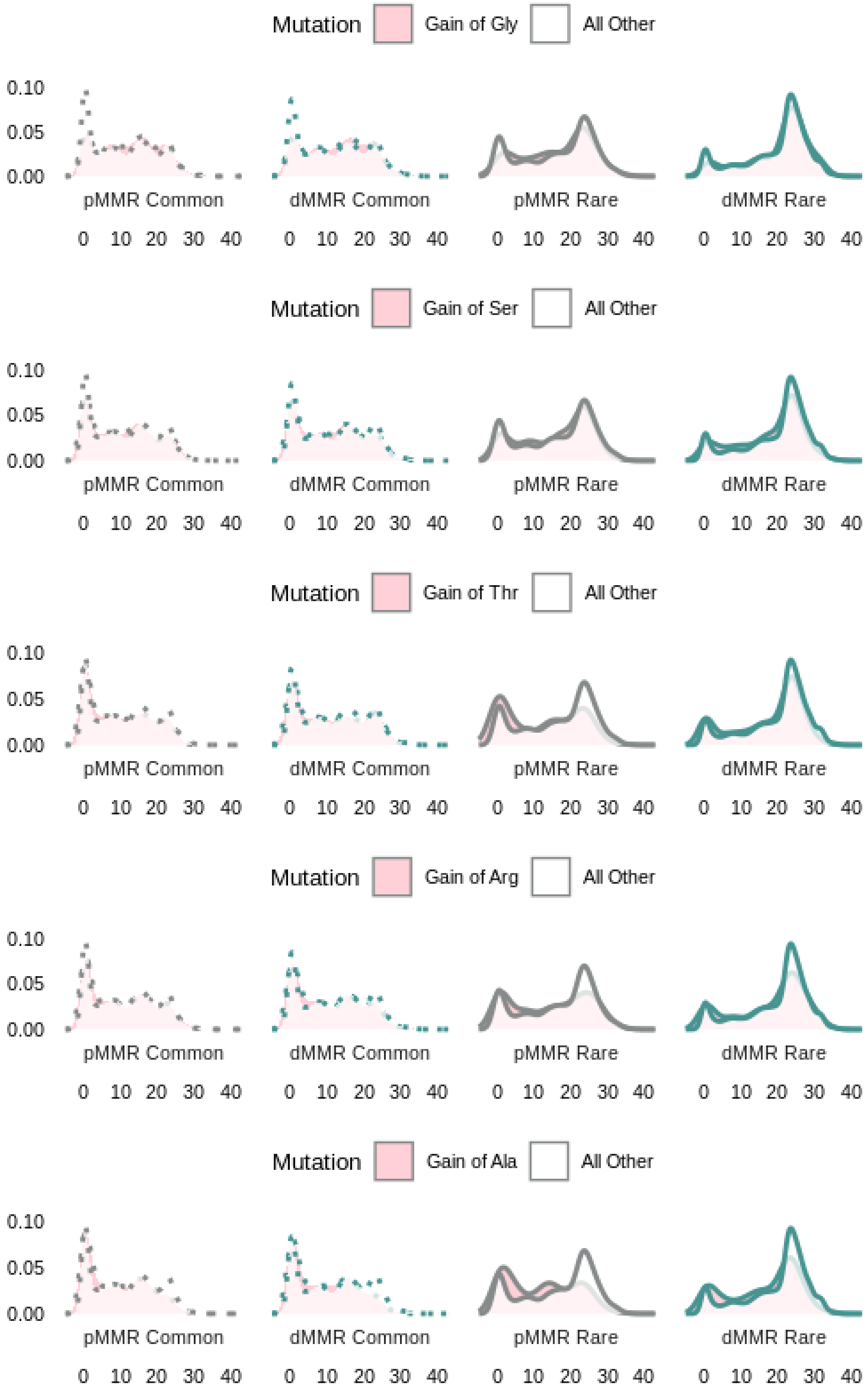
Mutation
Gain of Gly
All Other
0.10
0.05
0.00
pMMR Common
dMMR Common
pMMR Rare
dMMR Rare
0 10 20 30 40
Mutation
Gain of Ser
All Other
Mutation
Gain of Thr
All Other
Mutation
Gain of Arg
All Other
Mutation
Gain of Ala
All Other

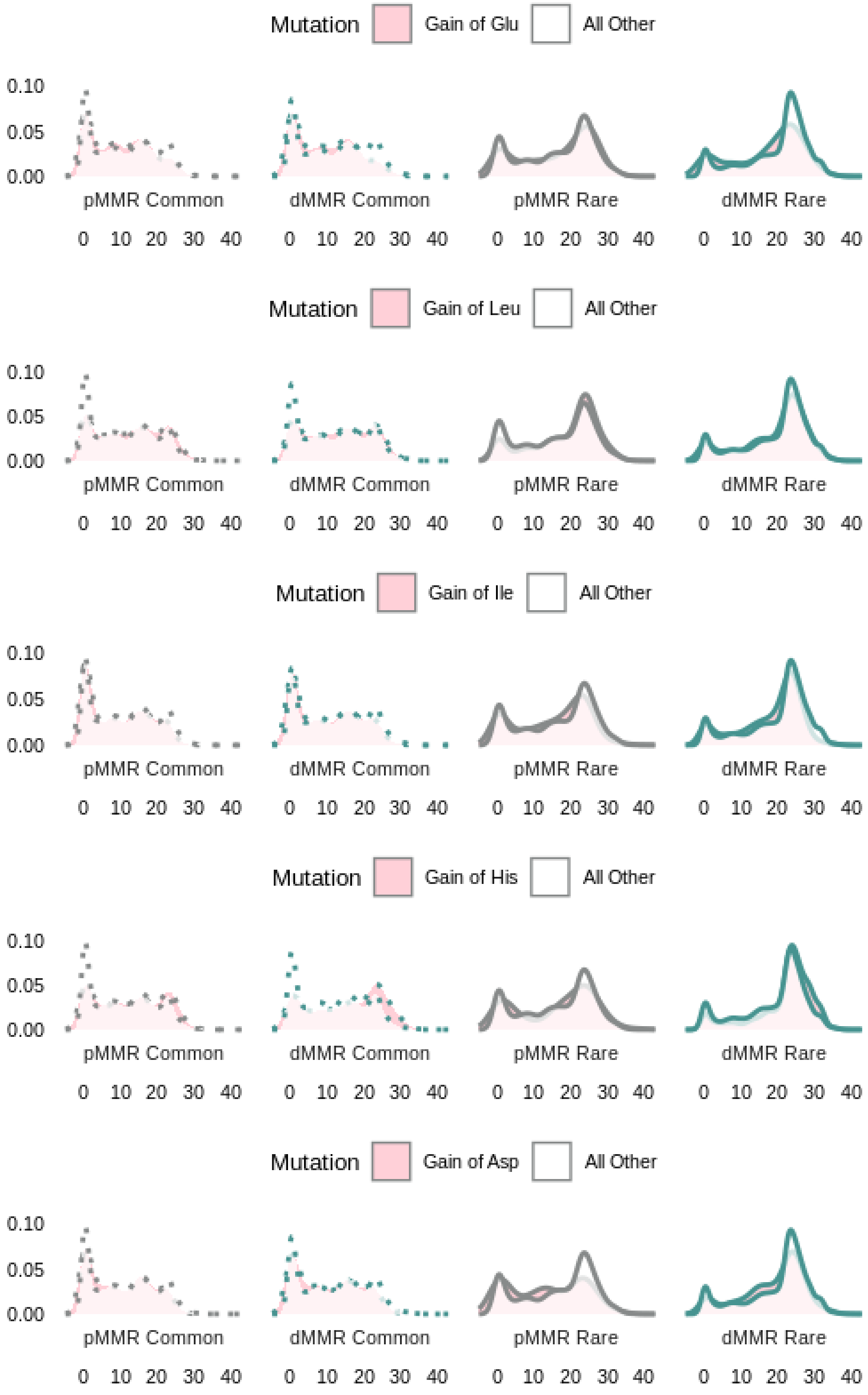
Mutation
Gain of Glu
All Other
0.10
0.05
0.00
pMMR Common
dMMR Common
pMMR Rare
dMMR Rare
0 10 20 30 40
Mutation
Gain of Leu
All Other
Mutation
Gain of Ile
All Other
Mutation
Gain of His
All Other
Mutation
Gain of Asp
All Other

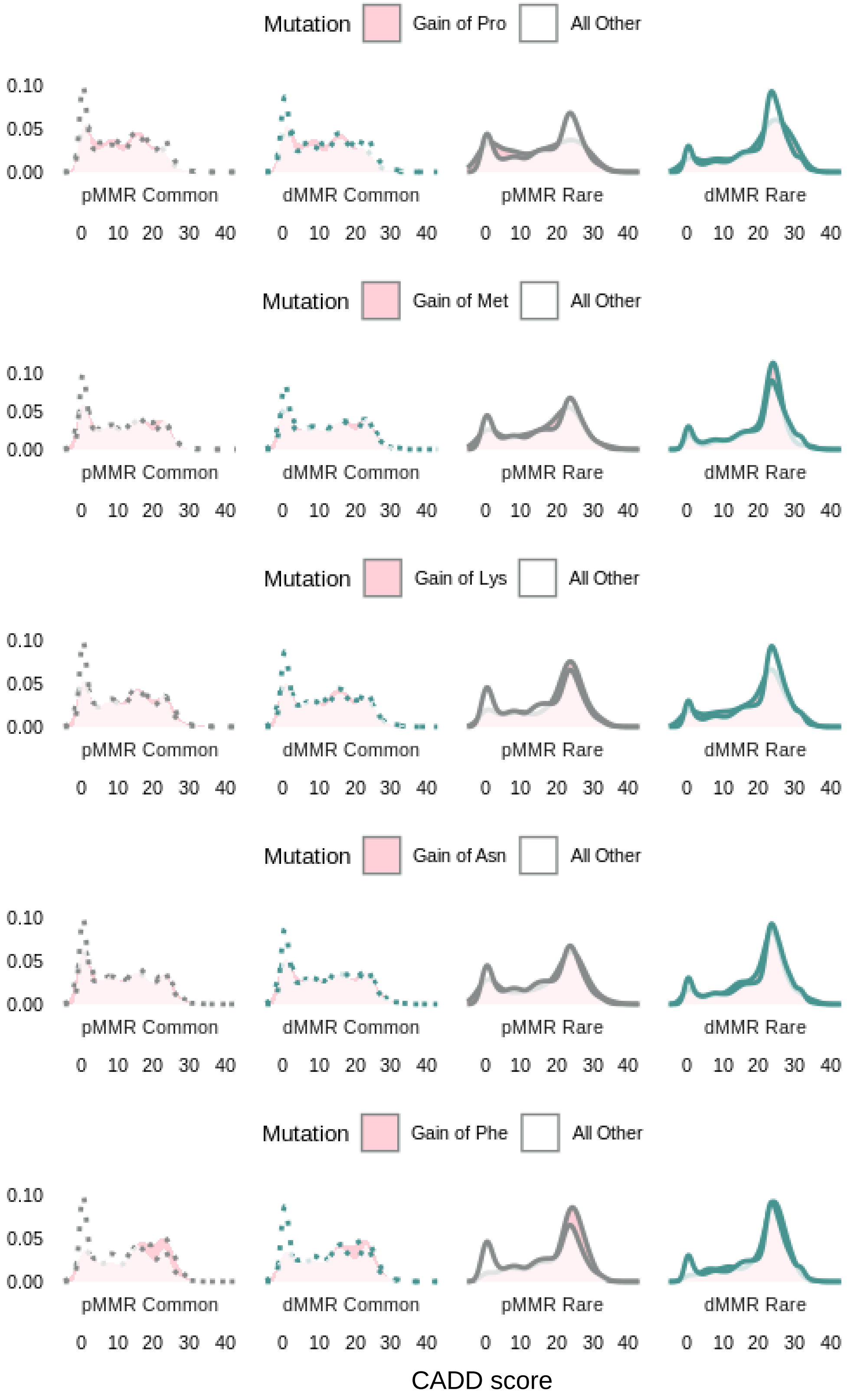


**Figure S19**. Distribution of CADD-phred scores for indicated amino-acid gained grouping. One sided Mann Whitney U test p-values are in **Table S2** (tab S16).

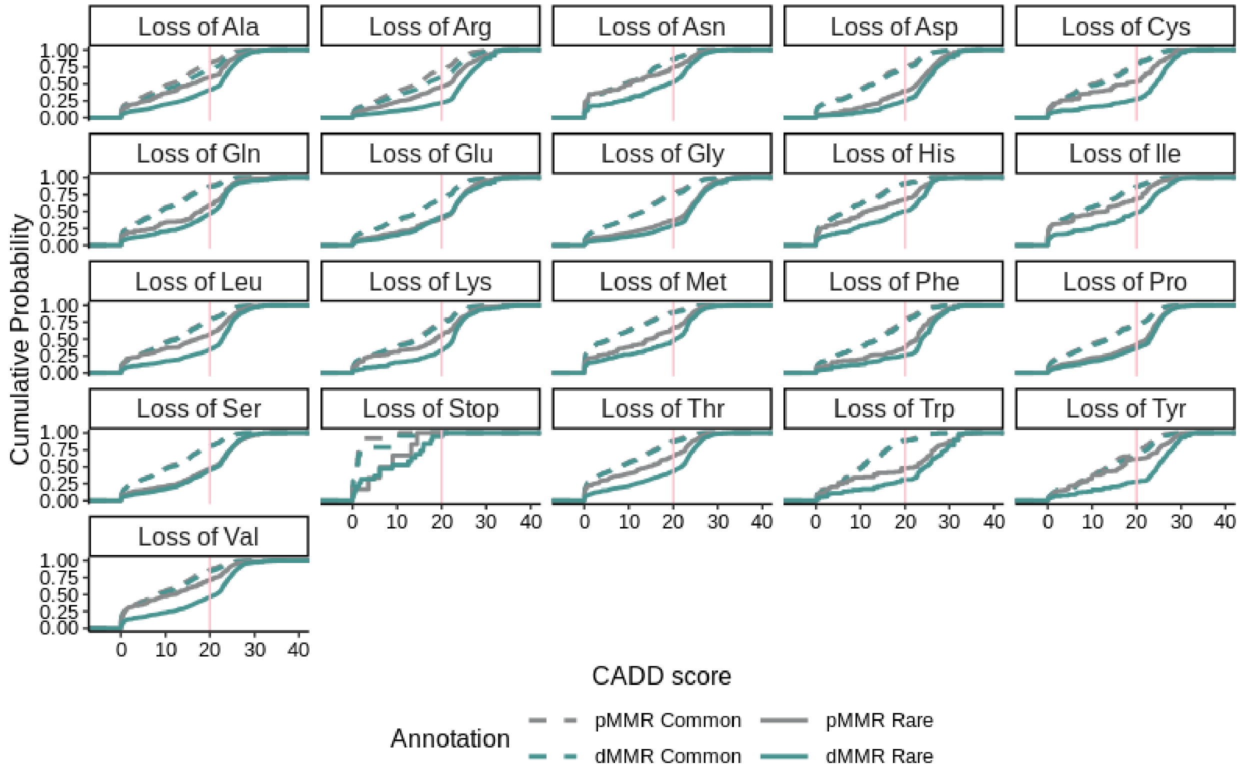


**Figure S20**. CADD-phred cumulative probabilities stratified by lost amino acid. Pink lines indicate CADD-phred = 20. One sided Mann Whitney U test p-values are in **Table S2** (tab S16).

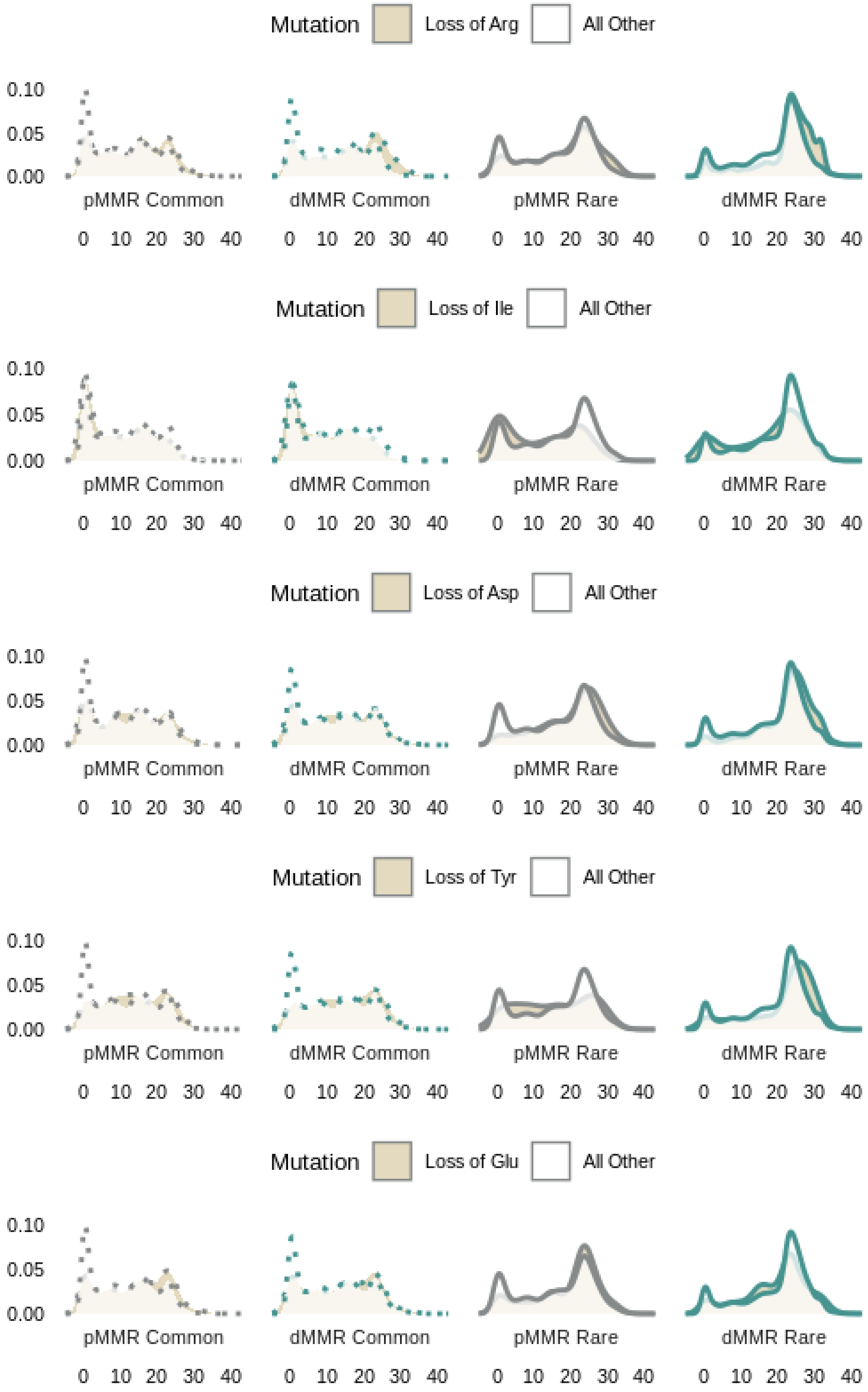
Mutation
Loss of Arg
All Other
Mutation
Loss of Ile
All Other
Mutation
Loss of Asp
All Other
Mutation
Loss of Tyr
All Other
Mutation
Loss of Glu
All Other
0.10
0.05
0.00
pMMR Common
dMMR Common
pMMR Rare
dMMR Rare
0 10 20 30 40

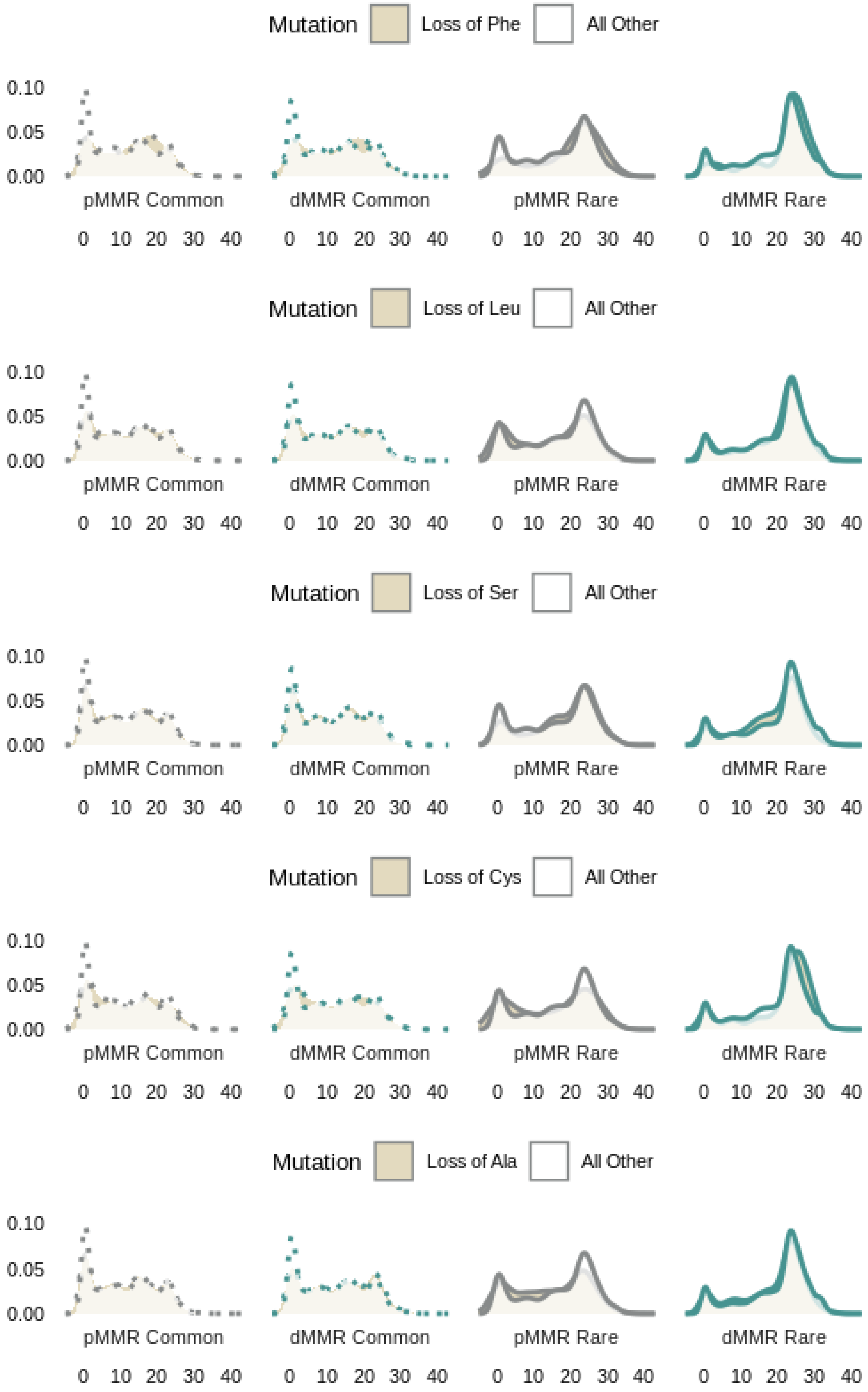
Mutation Loss of Phe All Other
0.10
0.05
0.00
pMMR Common
dMMR Common
pMMR Rare
dMMR Rare
0 10 20 30 40
Mutation Loss of Leu All Other
Mutation Loss of Ser All Other
Mutation Loss of Cys All Other
Mutation Loss of Ala All Other

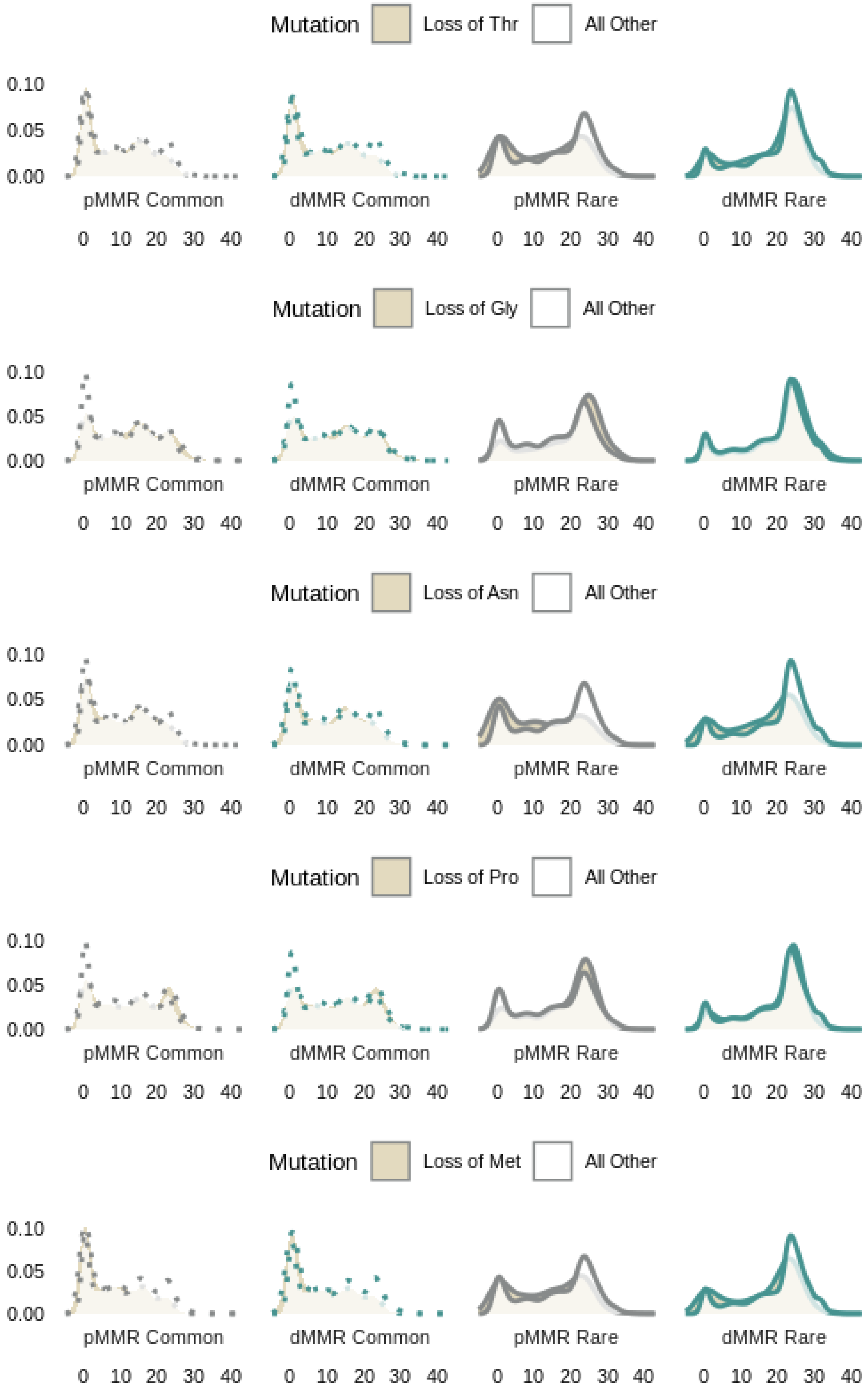
Mutation
Loss of Thr
All Other
Loss of Gly
Loss of Asn
Loss of Pro
Loss of Met
0.10
0.05
0.00
pMMR Common
dMMR Common
pMMR Rare
dMMR Rare
0 10 20 30 40

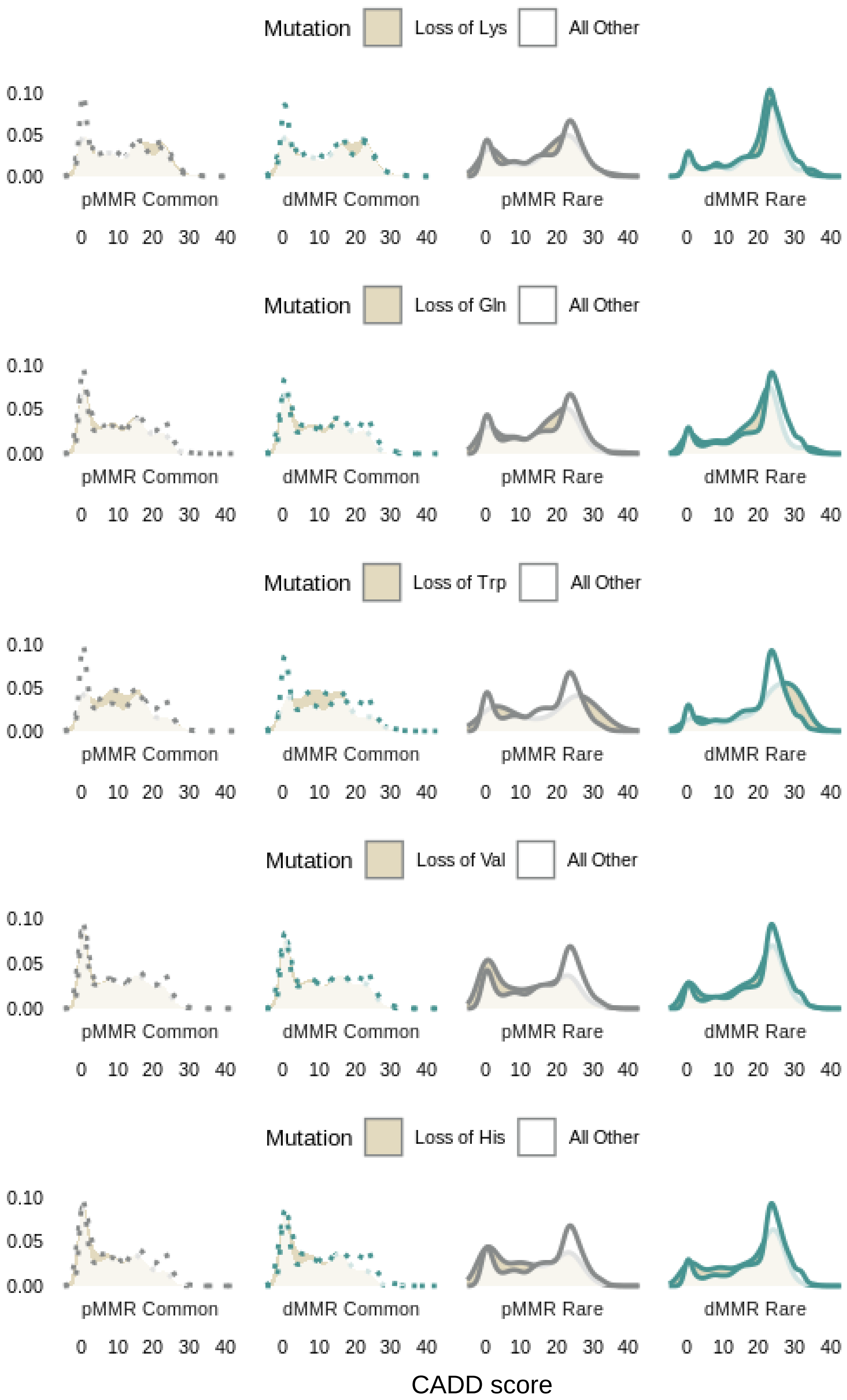


**Figure S21**. Distribution of CADD-phred scores for indicated amino-acid loss grouping. One sided Mann Whitney U test p-values are in **Table S2** (tab S16).

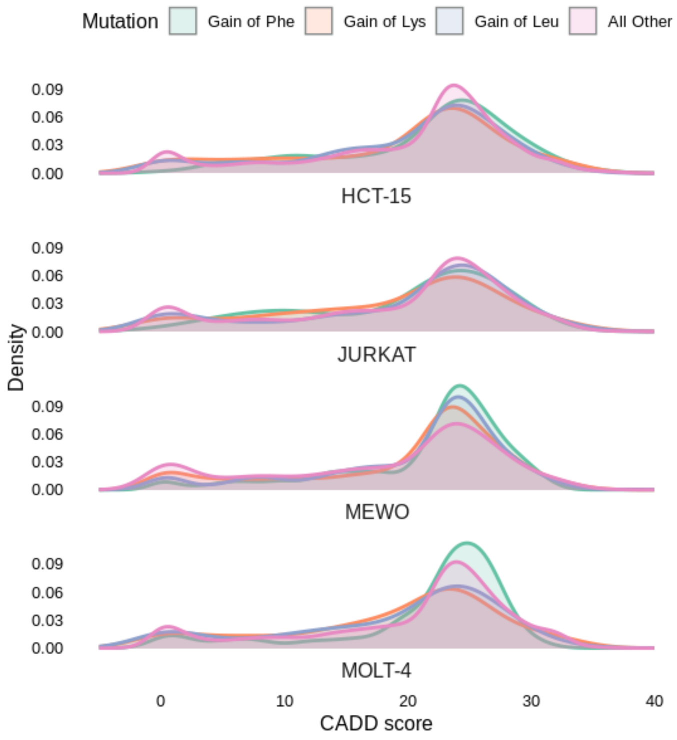


**Figure S22**. Distribution of CADD-phred scores for selected amino acid rare variants in MeWo, HCT-15, MOLT-4, and Jurkat cell lines

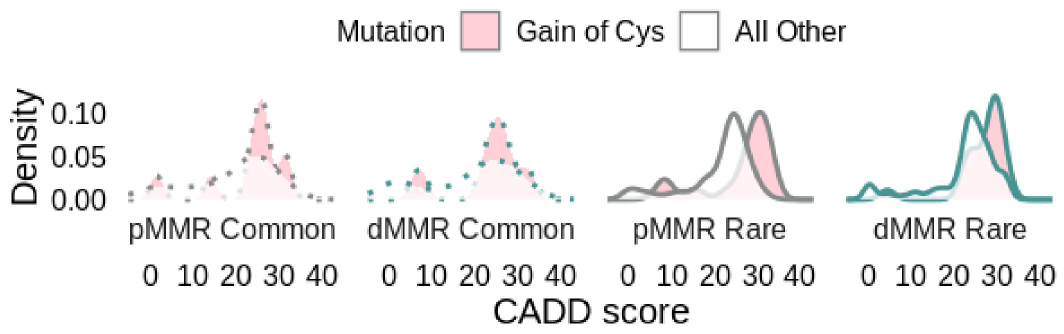


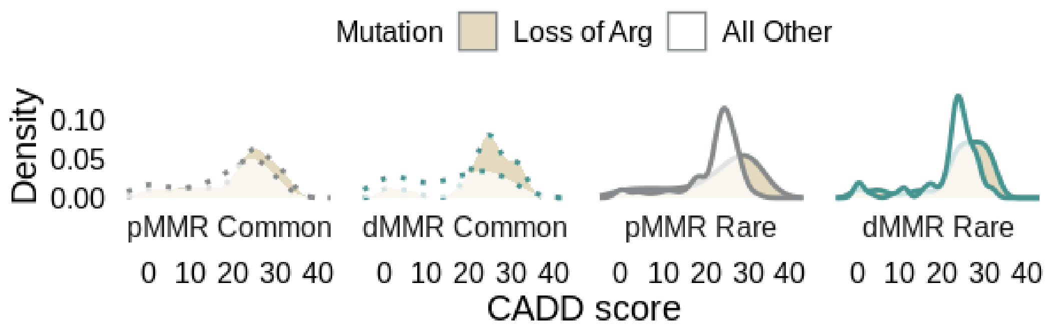


**Figure S23.** The Cancer Genome Atlas (TCGA) hotspot mutation distribution of CADD scores for indicated grouping. Highly deleterious gain of cysteines that are also TCGA hotspot mutations include FBXW7 R505C in Jurkat and ADAMTS1 R604C in MOLT-4, and SCYL3 R61C in MeWo. n=10, 6,16,21 for Cys ; n=25, 34, 31, 90 for Arg across from left.

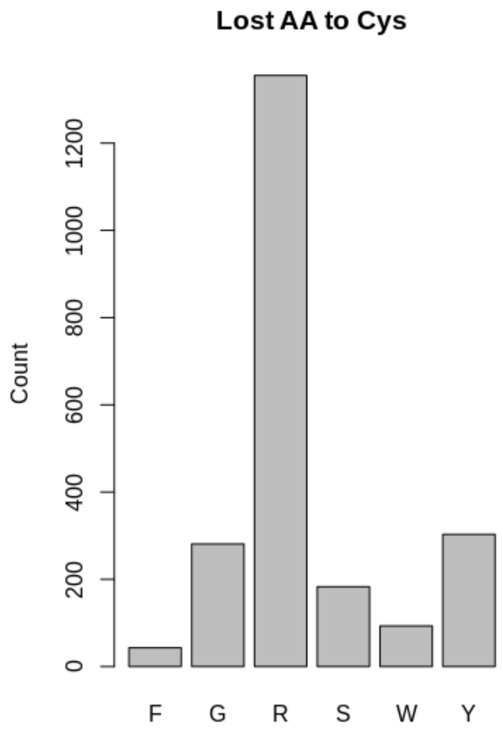


**Figure S24.** Distribution of reference/lost amino acids of gained cysteines in **Figure 2** data (60% arginine).

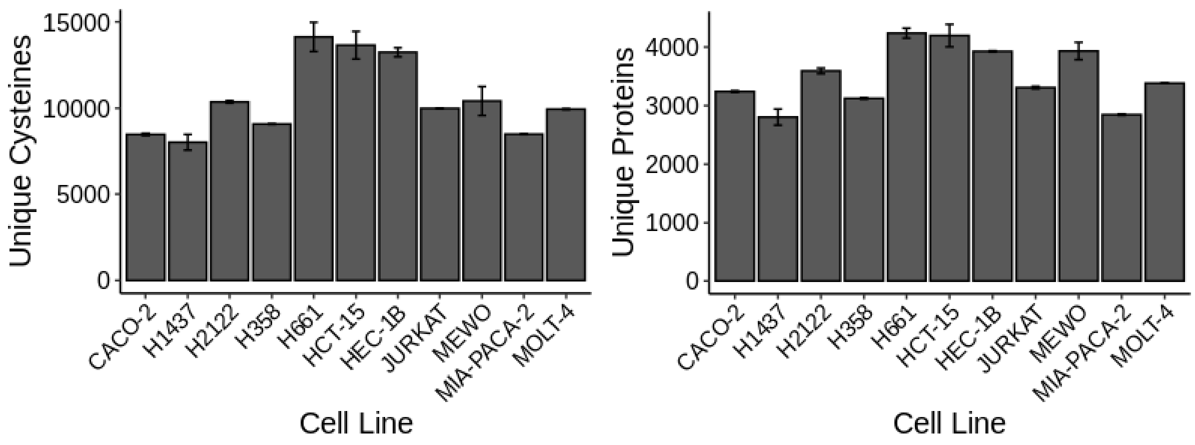


**Figure S25**. Reference cysteines and proteins identified per cell line in **Figure 4** for duplicate datasets.

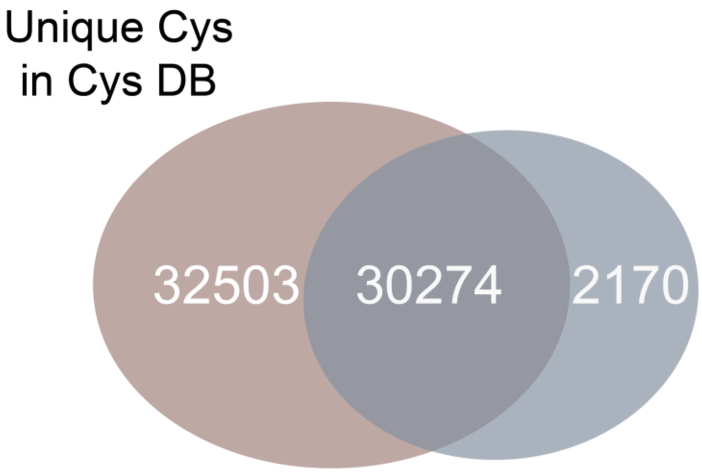


**Figure S26**. **Figure 4** reference cysteines overlap with CysDB cysteines. Red indicates Cys DB dataset, blue indicates **Figure 4** dataset.

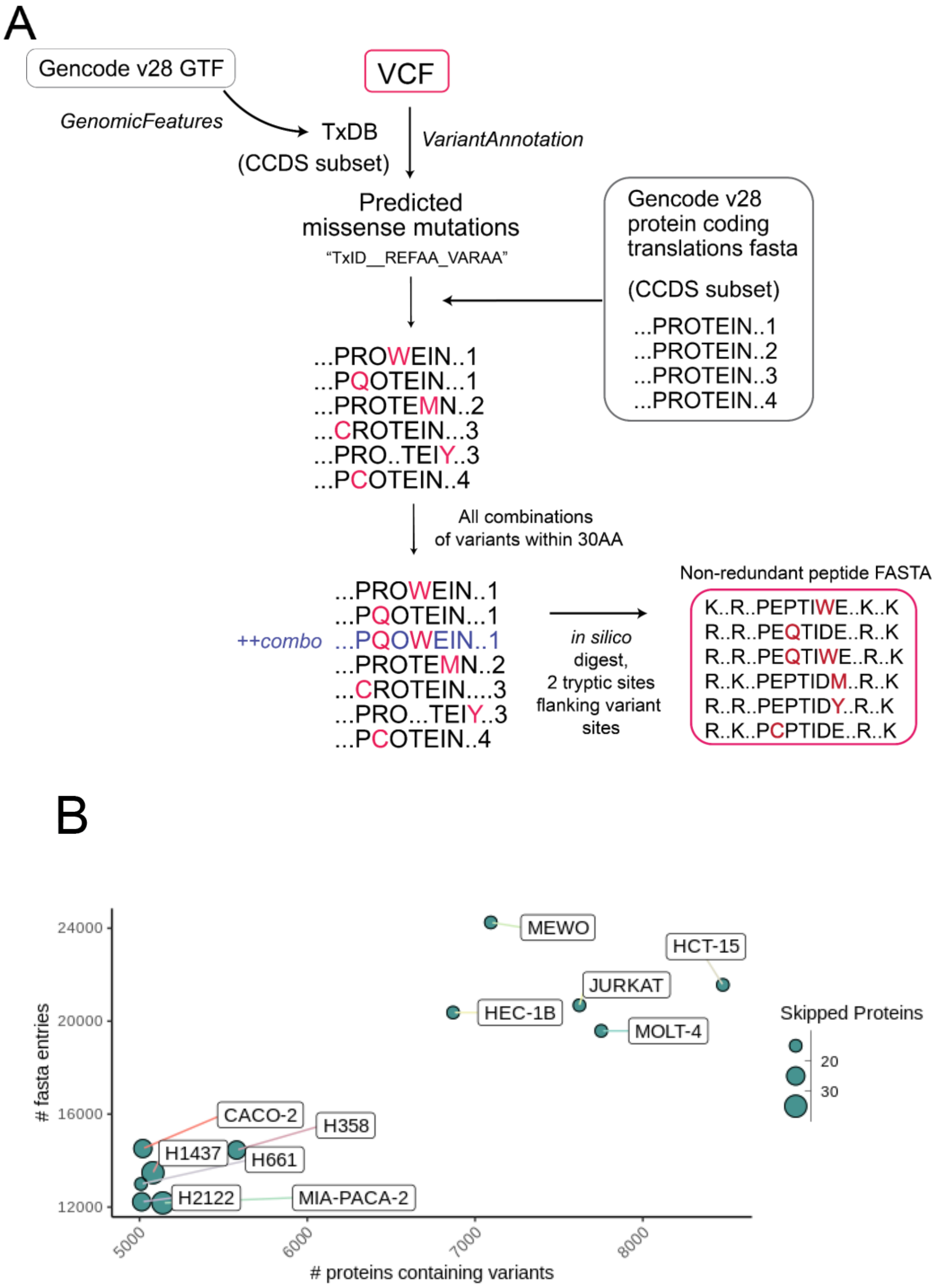


**Figure S27:** Details of database generation. A) Pipeline of packages and inputs to obtain variant peptide databases–*VariantAnnotation*[1] package is used to obtain predicted changes that replace reference sequence residues from Gencode by matching internal transcript IDs (TxID). Details in methods. Combinations are omitted from proteins with > 25 or 15 variants. B) Variant database sizes calculated as the number of FASTA entries. The skipped proteins indicate the number of proteins that were omitted from combinations in **Table S4** (tab S24).

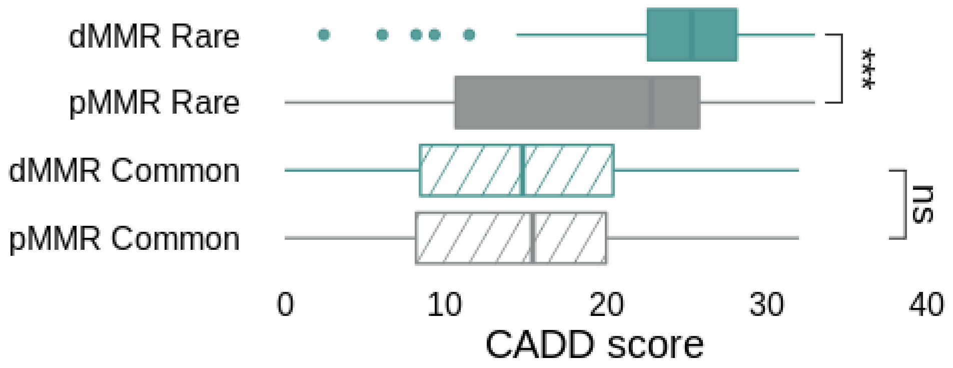


**Figure S28:** Distribution of CADD-phred scores for indicated variant grouping from **Figure 4** data. Statistical significance was calculated using Mann-Whitney U test, *** p < 0.001.

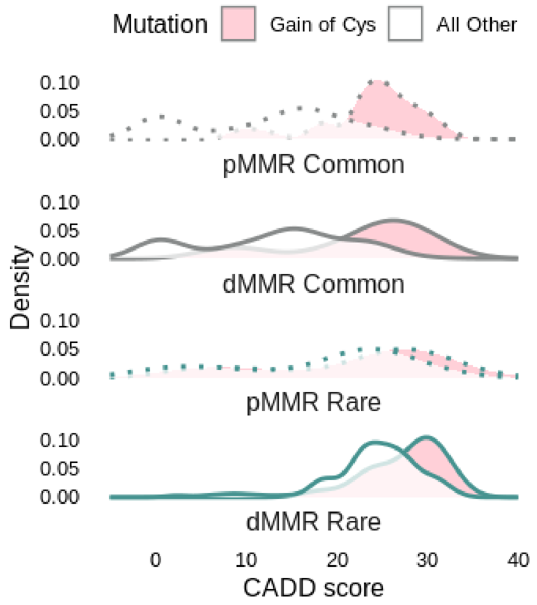


**Figure S29:** Distribution of CADD-phred scores for indicated gained cysteines vs all other from **Figure 4** data.

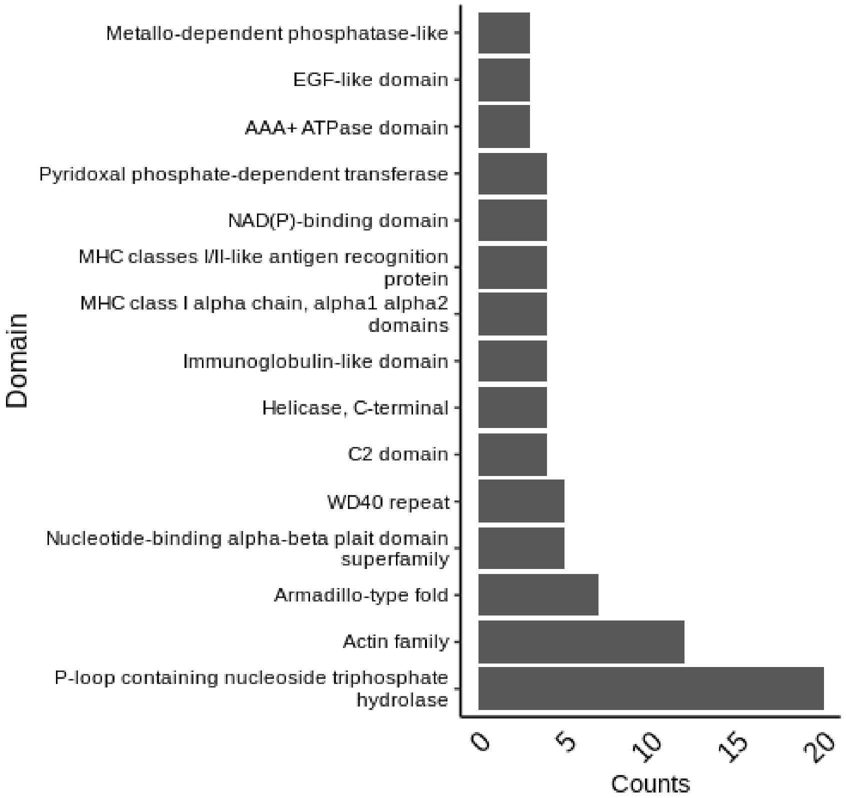


**Figure S30:** Identified variants' domain residence counts from Figure 4 data.

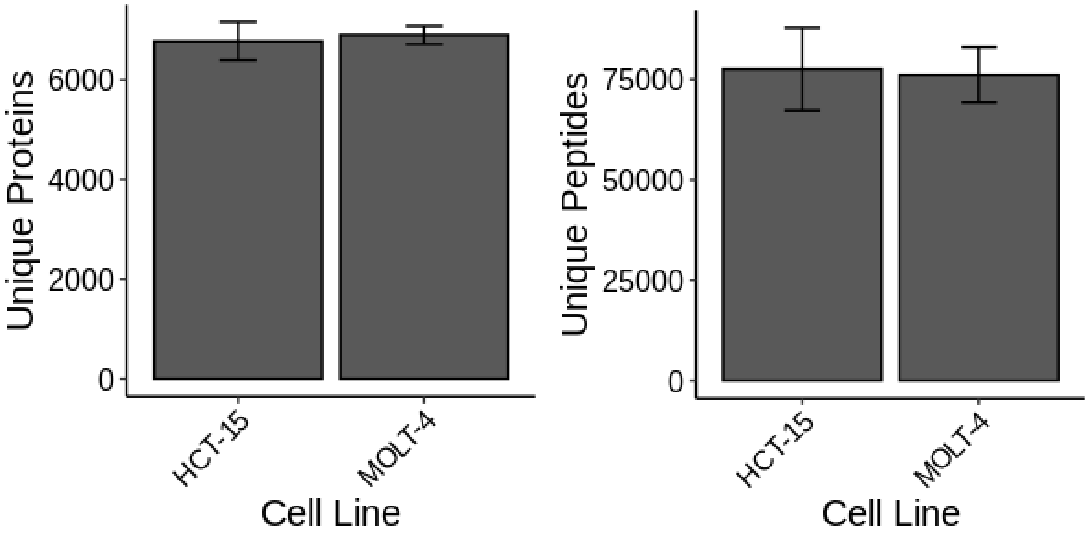


**Figure S31**. Reference peptides and proteins identified per cell line in Figure 5 for triplicate sets of high pH fractionated samples per cell line.

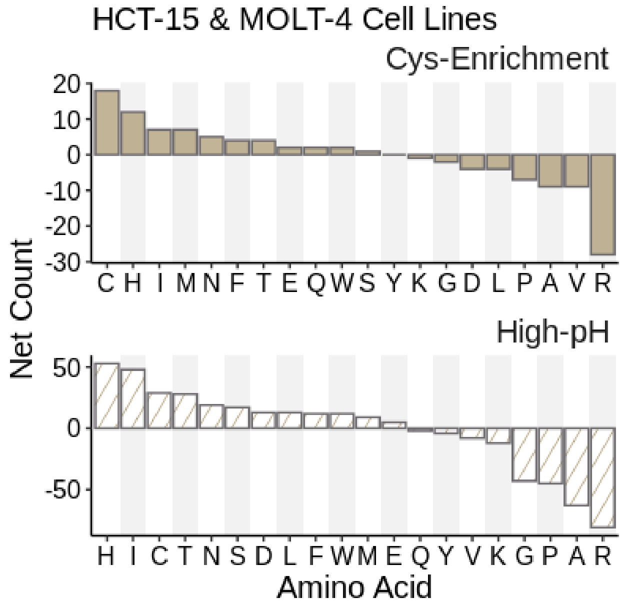


**Figure S32**. Net counts of SAAVs identified in Figure 5.

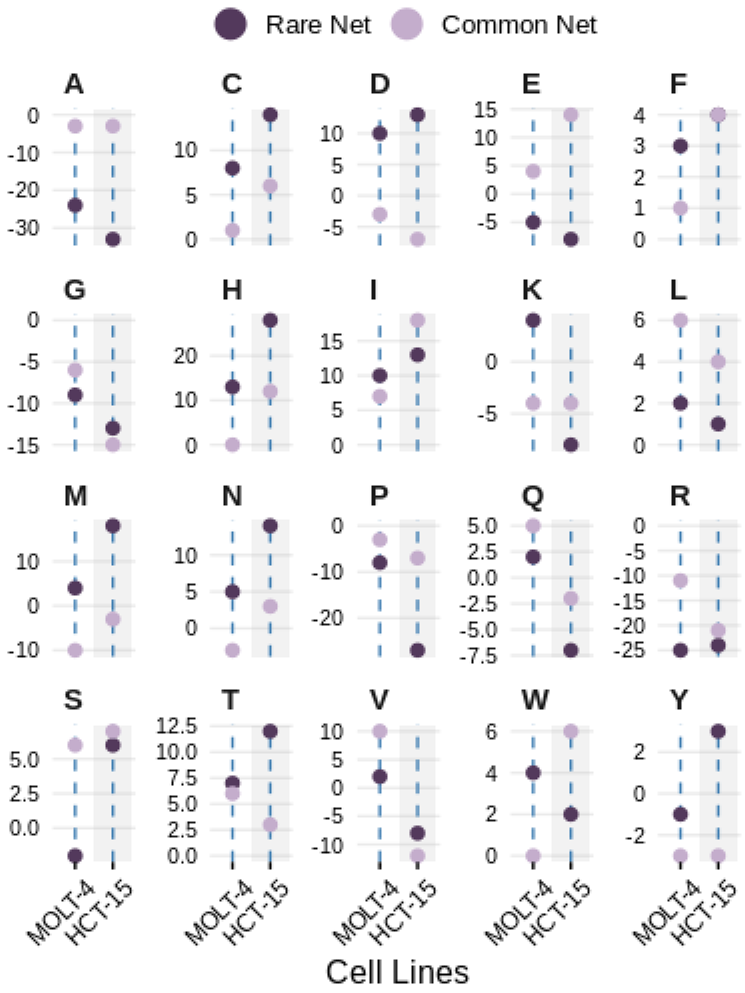


**Figure S33.** Net counts of SAAVs stratified by amino acid and common vs rare identified in Figure 5.

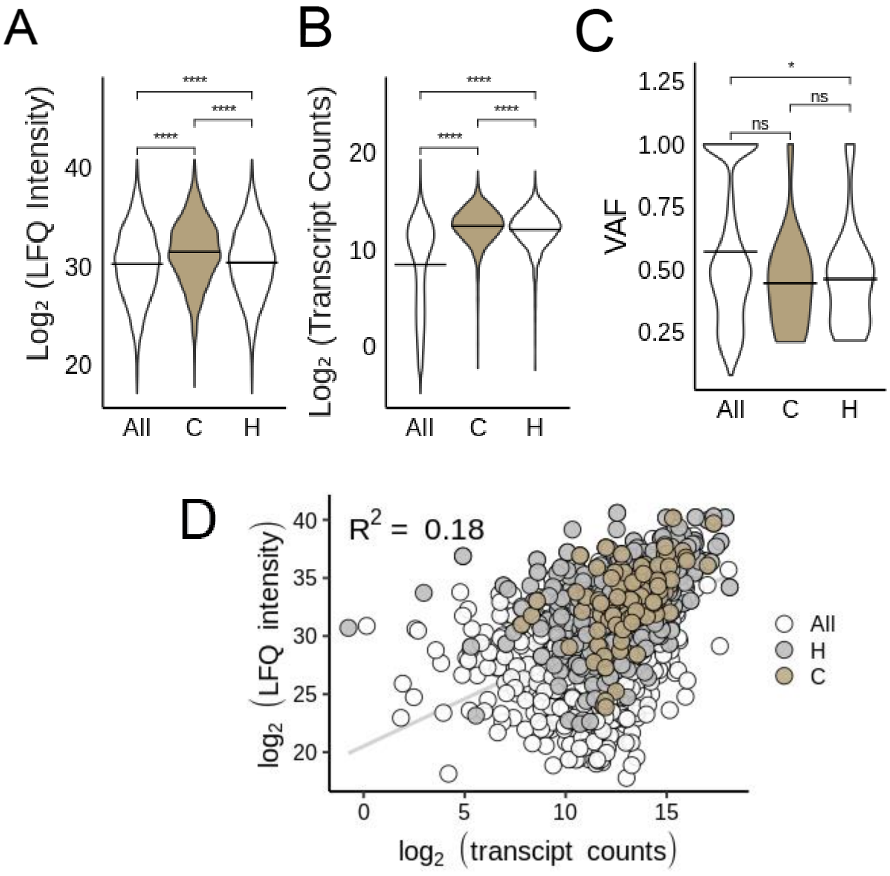


**Figure S34:** A) Label free quantitation (LFQ) intensities comparisons and B) DE-seq normalized transcript comparisons from HCT-15 cell line reference database searches. C) Variant allele frequencies for gain-of-cysteines subset in HCT-15 and Molt-4 searches. D) Matched LFQ intensities and normalized transcript count correlation of variant containing proteins/transcripts; All=all proteins in LFQ search or all transcripts, C=proteins from cys-enrichment search, H=proteins form high-pH fractionation search.

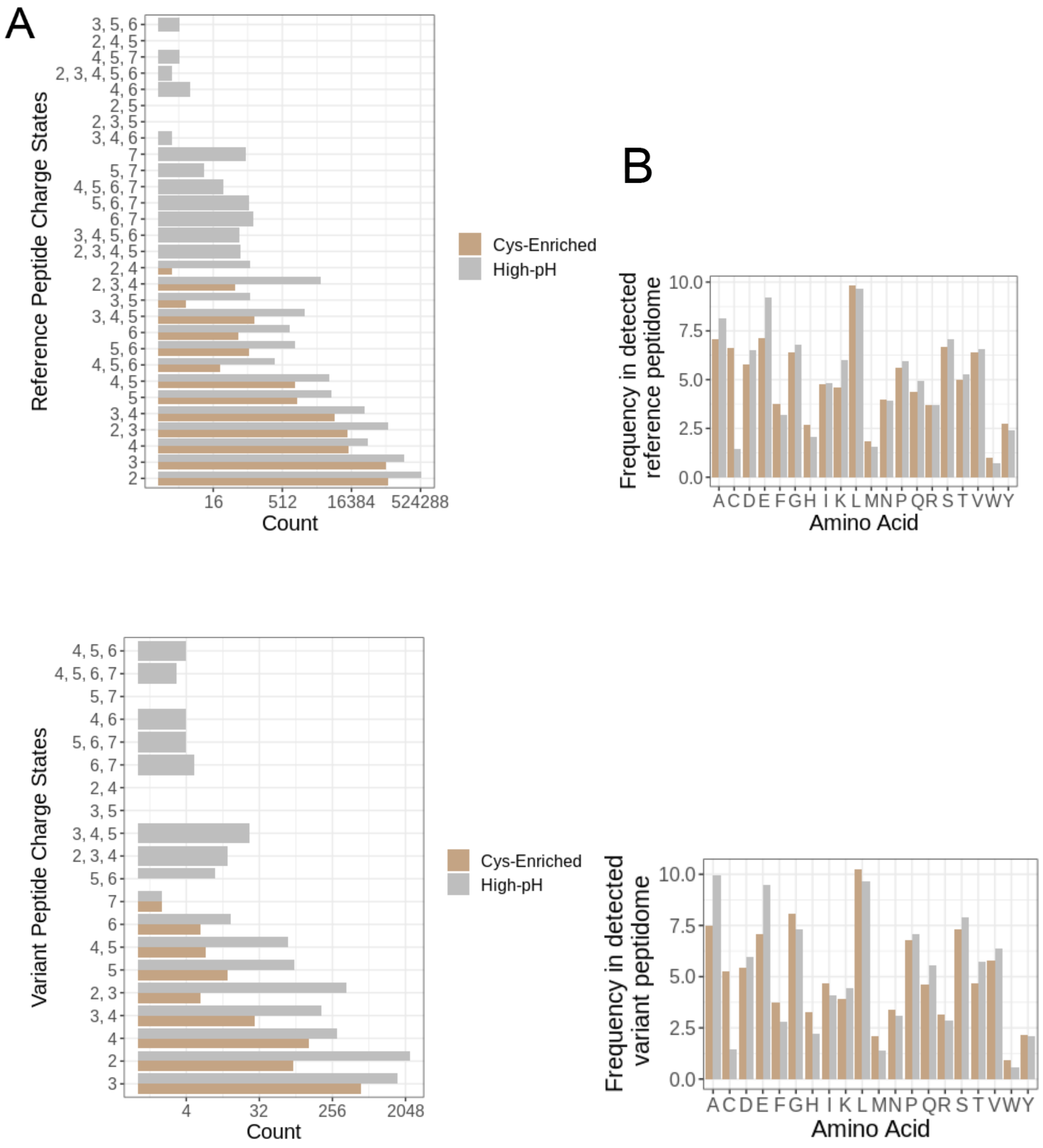


**Figure S35:** Peptide properties of detected reference peptides from cys-enrichment and high-pH fractionation A) Charge states of detected reference peptides and variant peptides. B) Abundance of amino acids in detected reference and variant peptides.

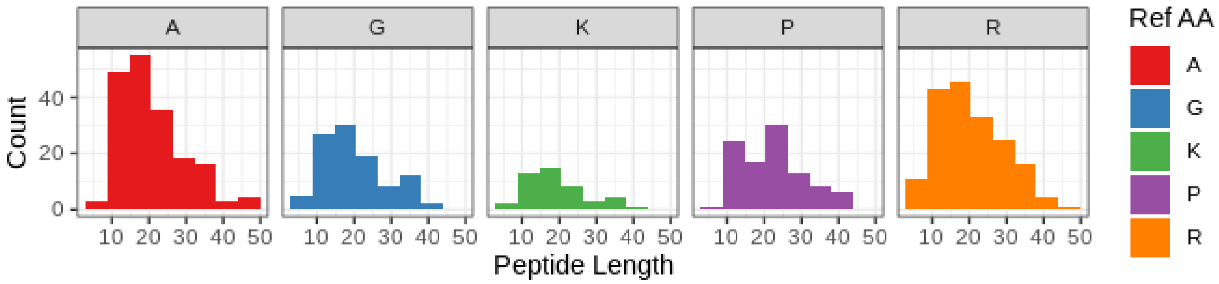


**Figure S36:** High-pH identified variant peptide length and SAAV reference amino acid for top lost amino acids.

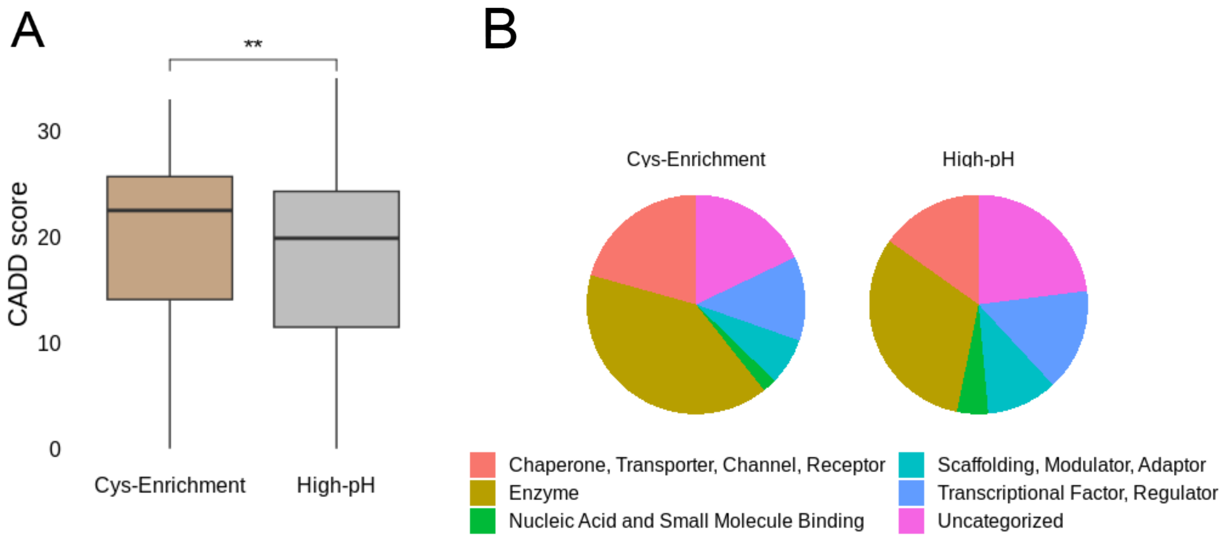


**Figure S37:** A) CADD score comparison between cys-enriched and high-pH detected variants B) Protein classes of cys-enriched and high-pH detected variants.

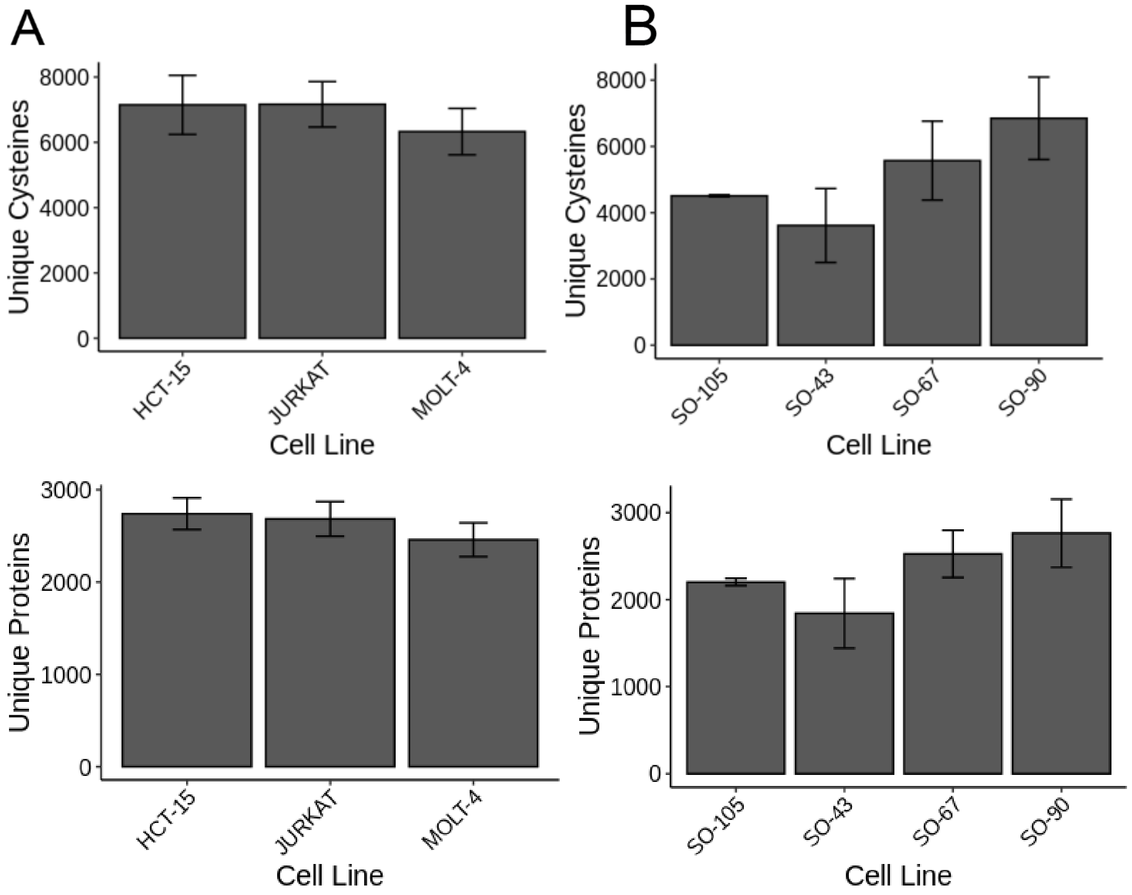


**Figure S38.** Reference identifications in Figure 6 for for A) KB02 datasets or B) HCT-15 datasets.

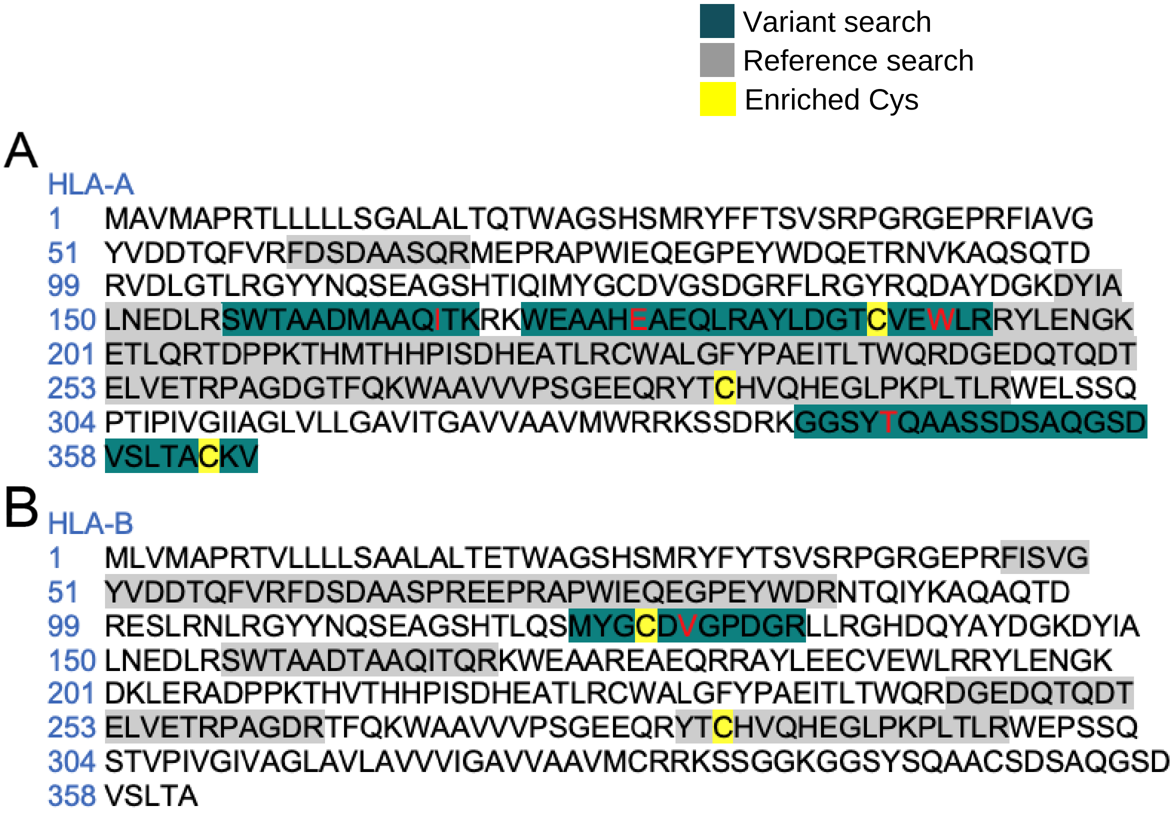


**Figure S39.** Sequence coverage of Uniprot references A) HLA-A*3:01 and B) HLA-B*7:02 peptides from HCT-15 reference and variant searches in Figure 4-6 datasets; yellow indicates enriched cysteines and red are variant sites.

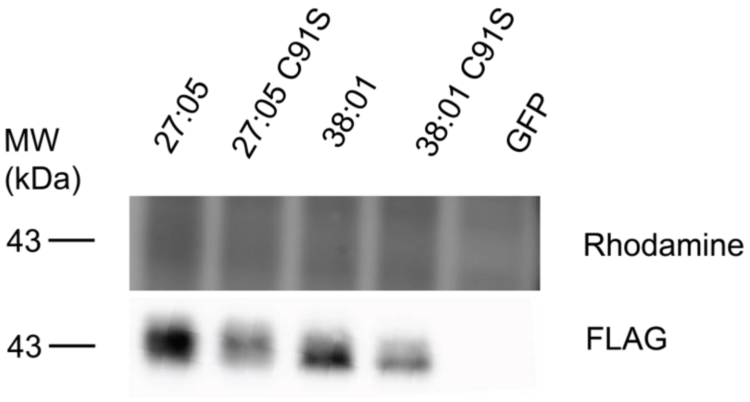


**Figure S40.** Cysteine lysate labeling of overexpressed HLA-B alleles with iodoacetamide alkyne (IAA) (representative of 2 two biological replicates) conjugation by click chemistry to a rhodamine-azide tag described in methods.

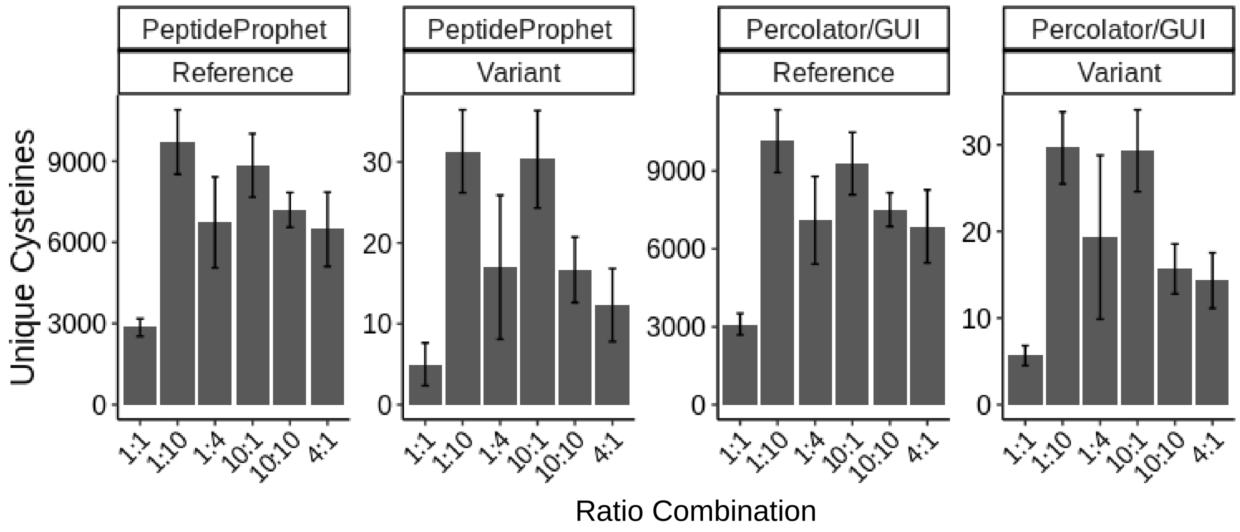


**Figure S41.** Coverage of FragPipe GUI implementing MSBooster with Percolator rescoring in comparison to PeptideProphet command line quantified cysteine identifications from validation datasets.

### (B) Methods

**Biology:**

**Cell culture and preparation of cell lysates.** Cell culture reagents including Dulbecco's phosphate-buffered saline (DPBS), Dulbecco's modified Eagle's medium (DMEM)/high glucose media, Eagle's Minimum Essential Medium (EMEM), Roswell Park Memorial Institute (RPMI) media, trypsin-EDTA and penicillin/streptomycin (Pen/Strep), and Horse Serum, heat inactivated (26-050-070) was purchased from Fisher Scientific. Fetal Bovine Serum (FBS) was purchased from Avantor Seradigm (lot # 214B17). All cell lines were obtained from ATCC and were maintained at a low passage number (< 20 passages).  HEK293T (ATCC: CRL-3216) cells were cultured in DMEM supplemented with 10% FBS and 1% antibiotics (Penn/Strep, 100 U/mL). MIA-PaCa-2 (ATCC: CRL-1420) cells were cultured in DMEM supplemented with 10% FBS, 1% antibiotics (Penn/Strep, 100 U/mL), and 2.5% horse serum. H661 (ATCC: HTB-183), H1437 (ATCC: CRL-5872), H358 (ATCC: CRL-5807), HCT-15 (ATCC: CCL-225), Jurkat (ATCC: TIB-152), MOLT-4 (ATCC: CRL-1582) and H2122 (ATCC: CRL-5985) cells were cultured in RPMI-1640 supplemented with 10% FBS and 1% antibiotics (Penn/Strep, 100 U/mL). HEC-1-B (ATCC: HTB-113), MeWo (ATCC: HTB-65), CaCo-2 (ATCC: HTB-37) cells were cultured in EMEM supplemented with 10% FBS and 1% antibiotics (Penn/Strep, 100 U/mL). Cells were maintained in a humidified incubator at 37 °C with 5% CO2. Cells were harvested by centrifugation (4500g, 5 min, 4 °C) and washed twice with cold DPBS. Cell pellets were then lysed with sonication (amp=10, 10 x1 sec pulses). The lysates were then transferred to a new microcentrifuge tube. Protein concentrations were determined using a BioRad DC protein assay kit from Bio-Rad Life Science (5000113, 5000114) and the lysate diluted to the working concentrations indicated below.

**RNA-seq variant calling.** Total RNA was extracted from cells using the Invitrogen Purelink RNeasy Plus Mini Kit (Qiagen, 166043750) or PureLink RNA mini kit (ThermoFisher, 12183018A) or Library preparation and RNA sequencing was carried out by the UCLA Technology Center for Genomics and Bioinformatics (TCGB). Libraries were prepared using the KAPA stranded mRNA kit.  Paired-end sequencing (2x150) was performed to a depth of 50-60x with an Illumina HiSeq3000 system. RNAFastq paired-end reads for each cell line were aligned to Gencode reference genome hg38 (GRCh38.p13) using STAR-2 PASS alignment (v2.7.3a)[2]. We ran STAR with default settings for paired-reads and the following additional parameters:–outSAMtype BAM SortedByCoordinate, --outSAMunmapped Within, and --sjdbFileChrStartEnd in second alignment. Samtools (v1.7)[3,4] calmd was used to add MD tags to sorted BAM files. Opposum (v0.2, 02-23-2017)[5] was used to split reads and mark duplicates using default parameters and an additional parameter --SoftClipsExist True. Platypus (v0.8.1)[6] was used to call variants and generate VCF files using default parameters. Samtools flagstat was used to obtain BAM file mapped read counts. Raw reads submitted to Sequence Read Archive (SRA) as BioProject PRJNA997729.

**WE-seq variant calling.** Genomic DNA was extracted from cells using the Zymo Quick DNA Miniprep Plus Kit kit (Fisher Sci, 50-444-149). Library preparation and exome-sequencing was carried out by the UCLA Technology Center for Genomics and Bioinformatics (TCGB). Libraries were prepared using the Nimblegen Capturing Kit. Paired-end sequencing (2x150) was performed to a depth of 50-60x with an Illumina HiSeq3000 system. Fastq paired-end reads for each cell line were aligned to Gencode reference genome hg38 (GRCh38.p13) using BWA-MEM alignment and default parameters for paired-reads. Output SAM files were converted to BAM files using Samtools (v1.7) and Picard (v2.21.4) (https://broadinstitute.github.io/picard/) was used to generate coordinate-sorted BAM files with read groups added. Samtools was used to index the files and duplicates were marked with Picard. GATK-HaplotypeCaller (v4.1.8.1)[7] was used to split reads. Since we do not have matched normal samples, we opted to use the germline caller GATK-HaplotypeCaller for exome data. Variants were called using default parameters with the exception of the ploidy option which was set to the value outlined in **Table S2** (tab S11). GATK was used to index the VCF file and filter the variants using the following parameters: -window 35 -cluster 3 --filter-name FS --filter-expression "FS > 30.0" --filter-name QD --filter-expression "QD < 2.0". Samtools flagstat was used to obtain BAM file mapped read counts. Raw reads submitted to Sequence Read Archive (SRA) as BioProject PRJNA997729.

**HCT-15 expression analysis.** 5 biological replicates RNA extracts were sequenced as described and aligned to hg38 as described in *RNA-seq variant calling.* Kallisto (v0.46.1)[8] was used to estimate transcript counts with indexed Gencode v28 transcriptome (gencode.v28.transcripts.fa) and -b (bootstrap) set to 100. Abundance transcript files were normalized with DE-seq2 (v1.28.1)[9]. Counts table was subsetted to a curated set of nonredundant CCDS transcript ID's (24,950) in **Table S2** (tab S9) and mean counts were calculated for downstream analysis. Raw reads submitted to Sequence Read Archive (SRA) as PRJNA997729.

**Predicting amino acid changes: Table S2** (S9) (nonredundant CCDS transcript ID's) was used to remove redundant proteins. VCFs from variant calling pipelines for both RNA and WES were processed using R package 'Variant Annotation'[1]. First, a TxDB object was made using the Gencode v28 annotation GTF file. The 'predictCoding' function using genome hg38 (GRCh38.p13) was used to obtain protein level changes from the VCFs, and 'nonsynonymous' and 'nonsense' changes were extracted; the resulting table includes a set of internal transcript IDs labeled 'TXID'. A database of common SNPs from NCBI (04-23-2018 00-common_all.vcf.gz) was used to annotate SNPs from rare mutations. The output missense table (**Table S2** (tab S10)) lists reference/variant codons and amino acids. This table was filtered to contain matches to the CCDS set of 24,950 Ensembl transcript IDs only and those that resulted in single amino acid variants (SAAVs), ignoring small indels and multi-nucleotide variants (48,552 variants). Variants *passing* variant-calling filters were used in Figures 1-2 (48,301 variants) and non-PASS variants are included in the proteomics analyses following.

**Generation of sample-specific custom databases with all combinations of variants.** Several R packages were used in generating custom databases: *VariantAnnotation*[1], *GenomicFeatures*[10], *biomaRt*[11] and *BSgenome.Hsapiens.UCSC.hg38* ( 10.18129/B9.bioc.BSgenome.Hsapiens.UCSC.hg38). A curated set of CCDS transcript ID's (24,950) (**Table S2**) was used to subset the Gencode v28 protein coding translations FASTA file by Ensembl transcript IDs. These sequences consist of a non-redundant UniProtKB[12] subset of cross-referenced CCDS proteins. Using the previously generated TxDB object from '*Predicting* *amino acid changes*', and the biomaRt select function, corresponding TXID headers for the protein FASTA file were obtained by selecting 'TXID' with Ensembl transcript ID keys ('TXNAME'). Matching TXIDs from all SAAVs with new protein sequence TXIDs, positions in the corresponding wild-type protein sequences were replaced with the corresponding variant amino acid to generate a list of protein sequences containing only one variant per sequence. Protein sequences containing variants shared between RNA and exome-derived variants were grouped as one sequence. For proteins containing multiple variants, all possible combinations were generated for variants within 30 amino acids windows for proteins with 25 (or 15, see SI tables) or fewer total variants. Output sequences were written to a FASTA file with headers containing corresponding Uniprot-ID, Gene ID, Ensembl transcript ID, and missense changes as well as cell-line, and sequencing origin (RNA or WE). To limit the increased search space, the database variant protein sequences were *in-silico* digested. A custom python script was used to *in-silico* digest the FASTA to generate tryptic peptides containing 2 misscleavages (two tryptic sites flanking amino acids surrounding the individual variant). Any duplicated peptide sequences were removed to leave unique sequences. For compatibility with MSFragger-based searches, simplified FASTA headers were used containing only the Uniprot ID. Result peptides are mapped back to detailed FASTA files for variant information. Scripts are available at https://github.com/hdesai17/chemoproteogenomics.git.

**Proteomic sample preparation for unenriched sample analysis.** HCT-15 and MOLT-4 lysates were incubated in 2 M urea/PBS at RT (final concentration = 2 mg/mL). DTT (10 µL of 200 mM stock in water, final concentration = 10 mM) was added into each sample and the sample was incubated at 65 °C for 15 min. To this, iodoacetamide (10 µL of 400 mM stock in water, final concentration = 20 mM) was added and the solutions were incubated for 30 min at 37 °C. Following addition of 3 µL trypsin solution (Worthington Biochemical, LS003740, 1 mg/mL in 666 µL of 50 mM acetic acid and 334 µL of 100 mM $CaCl_2$, final weight = 2 ng), digest was allowed to proceed overnight at 37 ºC with shaking. The next day, 90 µL from each digest was combined with 210 µL water and 0.3 µL TFA (final concentration ~0.1% TFA and ~180 µg peptides). Samples were fractionated into low-bind eppendorf tubes using a high-pH reversed phase fractionation kit (Pierce, 84868). Fractions were dried (Speed Vac) then reconstituted with 15 µL 5% acetonitrile and 1% FA in MB water and analyzed by LC-MS/MS. Samples were fractionated in triplicates for a total of 48 samples.

**Proteomic sample preparation for cysteine-enrichment sample analysis.** Proteome samples (200 µL of 1 mg/mL, prepared as described in preparation of cell lysates) Samples were then labeled with 2 mM IAA (2 µL of 200 mM stock solution in DMSO, final concentration = 2 mM) for 1h at RT (700rpm). CuAAC was performed with biotin azide (2) (4 µL of 200 mM stock in DMSO, final concentration = 4 mM), TCEP (4 µL of fresh 50 mM stock in water, final concentration = 1 mM), TBTA (12 µL of 1.7 mM stock in DMSO/tbutanol 1:4, final concentration = 100 µM), and $CuSO_4$ (4 µL of 50 mM stock in water, final concentration = 1 mM) for 1h at ambient temperature. After CuAAC, 10 µL of 20% SDS was added to each sample. Samples were incubated with 0.5 µL benzonase (Fisher Scientific, 70-664-3) for 30 min at 37°C. The samples were then subjected to SP3 sample loading, SP3 digest and elution, NeutrAvidin enrichment and LC MS/MS analysis, as described below. Experiments were conducted in duplicate for each cell line.

**Proteomic sample preparation for ligandability screening.** HCT-15, MOLT-4, and MeWo proteome samples (200 µL of 2 mg/mL, prepared as described in preparation of cell lysates). Compound (500 µM) or DMSO vehicle was added to lysates for 1 hr (2 µL 50 mM stocks or 2 µL DMSO). Samples were chased with 2 mM IAA (2 µL of 200 mM stock solution in DMSO, final concentration = 2 mM) for 1hr. CuAAC was performed with heavy biotin azide (DMSO samples) or light biotin azide (compound labeled samples) (4 µL of 200 mM stock in DMSO, final concentration = 4 mM), TCEP (4 µL of fresh 50 mM stock in water, final concentration = 1 mM), TBTA (12 µL of 1.7 mM stock in DMSO/tbutanol 1:4, final concentration = 100 µM), and CuSO4 (4 µL of 50 mM stock in water, final concentration = 1 mM) for 1h at ambient temperature. After CuAAC, 10 µL of 20% SDS was added to each sample. Samples were incubated with 0.5 µL benzonase (Fisher Scientific, 70-664-3) for 30 min at 37 °C. The samples were then subjected to *SP3 sample loading* using 80 µL total bead volumes, *SP3 digest and elution*, *NeutrAvidin enrichment* and *LC MS/MS analysis*, as described below. Experiments were conducted in triplicate for each compound per cell line.

**SP3 sample loading.** SP3 sample cleanup was performed generally at a bead/protein ratio of 10:1 (wt/wt) (38). For each 200 µL sample, 20 µL (or 40 µL) Sera-Mag SpeedBeads Carboxyl Magnetic Beads, hydrophobic (GE Healthcare, 65152105050250, 50 µg/µL, total 1 mg) and 20 µL (or 40 µL) Sera-Mag SpeedBeads Carboxyl Magnetic Beads, hydrophilic (GE Healthcare, 45152105050250, 50 µg/µL, total 1 mg) were aliquoted into a single microcentrifuge tube and gently mixed. Tubes were then placed on a magnetic rack until the beads settled to the tube wall, and the supernatants were removed. The beads were removed from the magnetic rack, reconstituted in 1 mL of MB water, and gently mixed. Tubes were then returned to the magnetic rack, beads allowed to settle, and the supernatants removed. Washes 20 were repeated for two more cycles, and then the beads were reconstituted in 40 µL MB water. The bead slurries were then transferred to the proteome samples, incubated for 10 min at RT with shaking (1000 rpm).

**SP3 digest and elution.** Absolute ethanol (400 µL) was added to each sample, and the samples were incubated for 5 min at RT with shaking (1000 rpm). Beads were washed twice with 80% ethanol as described above. Beads were then resuspended in 200 µL 0.5% SDS in PBS containing 2 M urea. DTT (10 µL of 200 mM stock in water, final concentration = 10 mM) was added into each sample and the sample was incubated at 65 °C for 15 min. To this iodoacetamide (10 µL of 400 mM stock in water, final concentration = 20 mM) was added and the solution was incubated for 30 min at 37 °C with shaking. After that, absolute ethanol (400 µL) was added to each sample, and the samples were incubated for 5 min at RT with shaking (1000 rpm). Beads were then again washed three times with 80% ethanol in water (400 µL). Next, beads were resuspended in 150 µL PBS containing 2 M urea followed by addition of 3 µL trypsin solution (Worthington Biochemical, LS003740, 1 mg/mL in 666 µL of 50 mM acetic acid and 334 µL of 100 mM $CaCl_2$, final weight = 2 ng). Digest was allowed to proceed overnight at 37 ºC with shaking. After digestion, ~ 4 mL acetonitrile (> 95% of the final volume) was added to each sample and the mixtures were incubated for 10 min at RT with shaking (1000 rpm). Supernatants were then removed and discarded using the magnetic rack, and the beads were washed (3 × 1 mL acetonitrile). Peptides were then eluted from SP3 beads with 100 µL of 2% DMSO in MB water for 1 hour at 37 °C with shaking (1000 rpm). The elution was repeated again with 100 µL of 2% DMSO in MB water. Peptide concentration assay (Pierce, 23275) was performed to test the concentration of the peptide. The elution can be used for NeutrAvidin enrichment or analyzed by LC-MS/MS.

**NeutrAvidin enrichment of labeled peptides.** For each sample, 50 µL of NeutrAvidin® Agarose resin slurry (Pierce, 29200) was washed three times in 10 mL IAP (immunoaffinity purification) buffer (50 mM MOPS–NaOH (pH 7.2), 10 mM $Na_2HPO_4$, 50 mM NaCl) and then resuspended in 500 µL IAP buffer. Peptide solutions eluted from SP3 beads were then transferred to the NeutrAvidin® Agarose resin suspension, and the samples were then rotated for 2h at RT. After incubation, the beads were pelleted by centrifugation (21,000 g, 1 min) and washed by centrifugation (3 × 1 mL PBS, 6 × 1 mL water). Bound peptides were eluted with 60 µL of 80% acetonitrile in MB water containing 0.1% FA (10 min at RT). The samples were then collected by centrifugation (21,000 g, 1 min) and residual beads separated from supernatants using Micro BioSpin columns (Bio-Rad). The remaining peptides were then eluted from pelleted beads with 60 µL of 80% acetonitrile in water containing 0.1% FA (10 min, 72 °C). Beads were then separated from the eluants using the same Bio-Spin column. Eluents were collected by centrifugation (21,000 g, 1 min) and dried (SpeedVac). The samples were then reconstituted with 5% acetonitrile and 1% FA in MB water and analyzed by LC-MS/MS.

**Liquid-chromatography tandem mass-spectrometry (LC-MS/MS) analysis**. The samples were analyzed by liquid chromatography tandem mass spectrometry using a Thermo Scientific™ Orbitrap Eclipse™ Tribrid™ mass spectrometer coupled with a High Field Asymmetric Waveform Ion Mobility Spectrometry (FAIMS) Interface. Peptides were resuspended in 5% formic acid and fractionated online using a 18cm long, 100 µM inner diameter (ID) fused silica capillary packed in-house with bulk C18 reversed phase resin (particle size, 1.9 µm; pore size, 100 Å; Dr. Maisch GmbH). The 70-minute water acetonitrile gradient was delivered using a Thermo Scientific™ EASY-nLC™ 1200 system at different flow rates (Buffer A: water with 3% DMSO and 0.1% formic acid and Buffer B: 80% acetonitrile with 3% DMSO and 0.1% formic acid). The detailed gradient includes 0 – 5 min from 3 % to 10 % at 300 nL/min, 5 – 64 min from 10 % to 50 % at 220 nL/min, and 64 – 70 min from 50 % to 95 % at 250 nL/min buffer B in buffer A. For bulk fractionation data, the detailed 80 min gradient includes 0 – 3 min from 1 % to 10 % at 300 nL/min, 3 – 63 min from 10 % to 40 % at 220 nL/min, 63 – 73 min from 40 % to 50 % at 220 nL/min, and 73 – 80 min from 50 % to 95 % at 250 nL/min buffer B in buffer A. Data was collected with charge exclusion (1, 8,>8). Data was acquired using a Data-Dependent Acquisition (DDA) method comprising a full MS1 scan (Resolution = 120,000) followed by sequential MS2 scans (Resolution = 15,000) to utilize the remainder of the 1 second cycle time. Time between master scans was set 1 s and 3s for compound labeling datasets, validation datasets, and fractionation datasets. HCD collision energy of MS2 fragmentation was 30 %. Raw file names used for figures are in **Table S9**.

**Command-line MSFragger-based variant peptide identification and quantitation.** Raw data collected by LC-MS/MS were searched using a 2-stage search scheme implemented using custom bash scripts: MSFragger (version 3.5), Philosopher (version 4.2.2) and IonQuant (version 1.8.0) enabled[13–16]. Precursor and fragment mass tolerance was set as 20 ppm. Missed cleavages were allowed up to 2. Peptide length was set 7 - 50 and peptide mass range was set 500 - 5000. Cysteine residues were searched with variable modifications at cysteine residues for carboxyamidomethylation (+57.02146), biotin-azide (+463.2366), and heavy biotin-azide (+469.2742) added for quant searches in Figure 5 datasets. Labeling was set allowing for 3 max occurrences and 'all mods used in first search' checked. Peptide and protein level FDR were set to 1%. For ligandability screening, permissive IonQuant parameters allowed minimum scan/isotope numbers set to 1. First, raw spectra are searched with normal reference protein sequences (CCDS set) and peptide to spectrum matches (PSM) are filtered to 1% FDR. Custom bash scripts were used to extract 1% FDR filtered PSM scan numbers from this first search. Prior to a second search using the same parameters and custom database, a text file of these scan numbers is generated with leading zeros removed and included as option 'excluded_scan_list_file', allowing remaining scans to be searched with a a cell line-specific custom database containing Uniprot identifiers and tryptic peptide sequences as described in *Generation of sample-specific custom databases*. PeptideProphet[17] was used for rescoring for both searches. Bash scripts are available at https://github.com/hdesai17/chemoproteogenomics.git.

**FragPipe label-free quantitation of HCT-15 fractionation data.** Raw data collected by LC-MS/MS were searched with the default LFQ-MBR workflow provided by FragPipe. With each experimental group corresponding to fractionation set for a total of three intensity values per protein in combined.protein.tsv output. Mean LFQ intensities were calculated and the non-redundant set of Uniprot IDs (**Table S2** (tab S9)) was used in downstream analyses.

The MS search results and fasta files have been deposited to the ProteomeXchange Consortium (http://proteomecentral.proteomexchange.org) via the PRIDE partner repository[18] with the dataset identifiers PXD043879 for newly generated data, and PXD023059 and PXD029500 for re-analyzed data.

**FragPipe GUI with improved 2-stage search.** FragPipe generates a "fragpipe-second-pass.workflow" and a "fragpipe-files-second-pass.fp-manifest" after the first search. The manifest file points to the calibrated mzML files generated from the first pass. The workflow file has mass calibration and optimization turned off. Thus, using those two files, the second-pass search skips the calibration and searches the calibrated data. See document on 2-stage searches for more details.

**Transient expression of HLA-B alleles.** Expression plasmids (pTwist CMV) containing HLA-B*38:01, and HLA-B*27:05, HLA-B*38:01 C91S ,and HLA-B*27:05 C91S inserts with C-terminal FLAG-tags were obtained from Twist Bioscience. pDONR223_B2M_WT was a gift from Jesse Boehm & William Hahn & David Root (Addgene plasmid # 81810; http://n2t.net/addgene:81810; RRID:Addgene_81810) and subcloned using GateWay cloning into C-terminal FLAG destination vector generated from a pRK5 backbone vector, which was a kind gift from T Wucherpfennig**.** Plasmids were co-transfected into 60% confluent 6cm plated 293T cells using 14µL PEI, 140 µL serum-free DMEM, and 1 µg co-transfections or 2 µg eGFP expression plasmid. Cells were harvested after 24 hour transfections

**FLAG-IP and Gel based-ABPP of HLA-B alleles: cell surface and lysate labeling.** Cells were washed once with PBS and resuspended in 100µL serum-free DMEM. One-half of cells were rotated at RT in 200 µM IAA (1 µL of 10 mM IAA stock) for 1 hr for cell-surface labeling. After spinning down at 1800 xg, the supernatant was removed. Cells were lysed in 30 µL 2% CHAPS/PBS for 30 min on ice. Remainder cells were lysed in 30 µL 2% CHAPS/PBS. Dilute all samples to 300 µL with PBS and spin 1800 xg for 1 min. Samples were adjusted to 2 mg/mL using a Bio-Rad DC protein assay kit from Bio-Rad Life Science (Hercules, CA). 200 µL of unlabeled lysates were incubated with 200 µM IAA (2 µL of 20 mM IAA stock) for 1 hr RT. 50 µL EZred FLAG bead suspension per sample (Sigma, F2426) were washed with tris-buffered saline (TBS) buffer according to manufacturer recommendations. 50 µL washed beads were added to each sample and rotated for 2 hours at 4C. Beads were washed 3x with 500 µL TBS pelleted at 8200 xg and resuspended in 50µL PBS with 250 µg/mL 3x FLAG peptide (Sigma, F4799) and 0.2% NP-40 alternative (Millipore Sigma, 492016) and rotated for 30 min at 4C. Beads were pelleted at 8200 xg to capture eluted proteins. Eluant was clicked on to rhodamine-azide (Click Chemistry Tools, AZ109-5) (25 µM rhodamine-azide (1.25 mM stock), 1 mM Tris(2carboxyethyl)phosphine (TCEP) (SigmaAldrich) (50 mM stock), 100 µM Tris[(1-benzyl-1H-1,2,3-triazol-4-yl)methyl]amine (TBTA) (Sigma-Aldrich) (1.7 mM stock), 1 mM CuSO4 (50 mM stock)) for 1 hour at RT. All samples were denatured (5 min, 95 °C) and loaded onto 4-12% Criterion™ XT Bis-Tris gels with XT MOPS running buffer from Bio-Rad followed by semi-dry transfer to nitrocellulose membrane. 1:2000 dilution of anti-FLAG rabbit antibody (14793, Cell Signaling) and followed by 1:5000 IRDye® 800CW Goat anti-Rabbit IgG (102673-330, VWR) as well as 1:3000 GAPDH rabbit antibody (2118S, Cell Signaling) followed by 1:5000 IRDye® 680RD Goat anti-Rabbit IgG was used for visualization of loading and rho signal for IAA labeling.

**Data processing for cys-enrichment and quantitative ratio peptide analysis.** Custom R scripts were implemented to compile modified_peptide_label_quant.tsv (quant) outputs from command line MSFragger pipeline or FragPipe to count unique quantified cysteines. Unique cysteines were quantified for each dataset using unique identifiers consisting of a UniProt protein ID and the amino acid number of the modified cysteine and an additional parameter specifying single or double isotopic labeling (heavy and/or light). Unique proteins were established based on UniProt protein IDs. Residue numbers were found by aligning the peptide sequence to the corresponding UniProt ID protein sequence specified by FragPipe outputs. Variant residue sites were obtained by mapping peptide sequences to respective custom FASTA files containing variant info in the headers. For enriched samples, nonspecific non-Cys containing peptides were omitted from analysis. For ratios data, methionine oxidized peptides were omitted, unpaired heavy or light-identified peptides were kept by setting ratios to $\log_2(20)$ or $\log_2(1/20)$. Outputs were generated by taking the median H:L ratio among all tryptic peptides for unique cysteines in replicate datasets (modified_peptide_label_quant.tsv); mean ratio values were calculated across replicate datasets; quantified cysteines appearing in at least two replicates with ratio SD =< 1 were kept (Figure 6 and Figure 4 data) and no SD filter was applied for Figure 3 data to interpret ratio skew. For comparisons to CysDB cystines, unique UniprotID_CysPostion identifiers were used and FragPipe assigned Protein ID was used; for ClinVar variant matching, chromosome position and nucleotide changes of associated variants were used. The MS search results and

FASTA files have been deposited to the ProteomeXchange Consortium via the PRIDE partner repository with the dataset identifiers PXD043879 for newly generated data and FASTA files. R and Python scripts are available at https://github.com/hdesai17/chemoproteogenomics.git.

**Data processing for high-pH data analysis.** Custom R scripts were implemented to compile peptide.tsv outputs from command line MSFragger pipelines. Unique proteins were established based on UniProt protein IDs. Variant residue sites were obtained by mapping peptide sequences to respective custom FASTA files containing variant info in the headers. Residue numbers were found by aligning the peptide sequence to the corresponding UniProt ID protein sequence specified by FragPipe outputs. For ClinVar variant matching, chromosome position and nucleotide changes of associated variants were used.

**Linear sequence and spatial site-analysis.** For linear sequence analysis in Figure 2-5 datasets, residues 'in or near' UniProtKB annotated active or binding sites, DisProt annotated sites[19–21], or PhosphoSite annotated sites[22] were within +/-10 amino acids in linear sequence. For Figure 6 datasets, residues 'in or near' UniProtKB annotated active or binding sites, disprot annotated sites, or phosphosite annotated sites were assessed using 3D Protein Data Bank (PDB) structures. PDB structures were parsed to find all neighboring residues within a 10 Angstrom distance of the liganded Cys (alpha carbon atom). PDB_UniProtKB identifiers were created for each cysteine and corresponding list of neighboring residues. If the UniProtKB annotated active or binding sites were resolved in an associated crystal structure and found within the 10 Angstroms net, it was classified as a cysteine proximal to a known active or binding site.

**Data Analysis and Statistics:**

**Fig 1B, C, S1-S5,**: COSMIC Cell Lines Project Complete mutation data 'CosmicCLP_MutantExport.tsv.gz' was downloaded (release v96, 31st May 2022). For each cell line, the 'Substitution-Missense' amino acid changes were totaled up; the gained counts (example Cys, X–>C) were subtracted from the lost counts (example Cys, C–>X) to obtain net counts for each amino acid. The analysis was limited to accession numbers matching a curated set of non-redundant Ensembl transcript IDs (24,950) to limit any over/under counting due to mutations in multi-transcript proteins. **Fig S6**: ClinVar (3-28-23) data filtered for unique gene name,protein position, and amino acid change to calculate total gains and losses. **Fig 1D**: common SNPs from NCBI (04-23-2018 00-common_all.vcf.gz) were filtered for missense mutations (645,395) and further filtered to include only variants in the non-redundant Ensembl transcript set. Net changes were calculated from unique 'GeneID_amino acid change identifiers'.

**Fig 2B,C,D,S24**: Missense table in **Table S2** (tab S10) was filtered for unique 'CellLine_Protein_AAchange' identifiers, removing transcripts IDs that result in the same amino acid changes per protein. **Fig 2F,G,H,S11-S23:** CADD-PHRED score distributions were obtained by filtering the missense table for only single nucleotide base substitutions and uploading them as a vcf ('#CHROM', 'POS', 'ID', 'REF', 'ALT') to the CADD query tool https://cadd.gs.washington.edu/score using GRCh38-v1.6. Distribution of CADD scores shown by indicated grouping, unique 'CellLine_Protein-AAchange-CADD-score' identifiers were used to remove variants identified in both RNA and exomes. **Fig 2I,2J:** Active site/binding site analysis:

We consider single variant residues in or within +/- 10 amino acids in primary sequence from an annotated active/binding site range as annotated by Uniprot.

**Fig 3C (and Fig 8B):** Quantitative output obtained as described in *Data processing for cys-enrichment and quantitative ratio peptide analysis* for triplicate datasets. **Fig 3D**: From combined_ion_label_quant.tsv outputs for reference and variant searches, apex light - apex heavy retention times per ion were calculated for each replicate set. Mean values were obtained for ions appearing in at least two of three replicates.

**Fig 4:** Data was processed as described in methods for *Data processing for cys-enrichment* for duplicate datasets. **Figure S26:** Interpro ascension numbers and interpro family/domain names were pulled for all identified variant genes using EnsDb.Hsapiens.v86 from the *ensembldb* package[23] **Fig 4F**: The provided reference FASTA was *in-silico* digested to check for cysteines theoretically detectable using the FragPipe peptide length limits of 7-50 amino acids. Cysteine peptides with loss or gain of Arg/Lys were manually checked against theoretically detectable cysteines. **Fig 4J**: Site level analyses from Fig2. **Fig 4K:** SP3-Rox dataset were re-searched with 2-stage search and custom database; quantitative output obtained as described in data processing for triplicate datasets.

**Fig 5:** Data was processed as described in methods for *Data processing for high-pH data analysis*. **Fig 5D:** For each cell line, protein sequences from the subsetted Gencode FASTA provided in supplemental information (24,950 proteins) and as described in *custom database generation* were used to calculate individual amino acid frequencies (# amino acid/total # amino acids) for every protein that contained a missense mutation. The total gained AA (identified in proteomic data) was used to calculate the gained frequencies (# gained AA/total missense changes identified). The fold enrichment was calculated by taking the ratio of the gained frequency to the AA frequency. **Fig 5E:** Transcript counts and LFQ data for 'high-ph' and 'all' categories were restricted to those containing at least one cysteine residue.

**Fig 6:** Quantitative output obtained as described in *Data processing for cys-enrichment and quantitative ratio peptide analysis.* **Fig 6D:** Ligandable cys residues were restricted to those appearing in more than one replicate, with Log2 ratios > 2, and with standard deviations of ratios across replicates <= 1. **Fig 6C-G:** Values are restricted to those that appear in more than one replicate and with standard deviations =<1 for both reference and variant Cys peptides.

**Chemistry:**

**General Methods.** All solution-phase reactions were performed in dried glassware under an atmosphere of dry $N_2$ except where water was used as a solvent. Silica gel P60 (SiliCycle) was used for column chromatography. Plates were visualized by fluorescence quenching under UV light or by staining with iodine. Other reagents were purchased from Sigma-Aldrich (St. Louis, MO), Alfa Aesar (Ward Hill, MA), EMD Millipore (Billerica, MA), Fisher Scientific (Hampton, NH),

Oakwood Chemical (West Columbia, SC), Combi-blocks (San Diego, CA) and Cayman Chemical (Ann Arbor, MI) and used without further purification. $^{1}H$ NMR and $^{13}C$ NMR spectra for characterization of new compounds and monitoring reactions were collected in $CDCl_3$, $CD_3OD$, or DMSO-$d_6$ (Cambridge Isotope Laboratories, Cambridge, MA) on a Bruker AV 400 MHz spectrometer in the Department of Chemistry & Biochemistry at The University of California, Los Angeles. All chemical shifts are reported in the standard notation of parts per million using the peak of residual proton signals of the deuterated solvent as an internal reference. Coupling constant units are in Hertz (Hz). Splitting patterns are indicated as follows: br, broad; s, singlet; d, doublet; t, triplet; q, quartet; m, multiplet; dd, doublet of doublets; dt, doublet of triplets. Low-resolution mass spectroscopy was performed on an Agilent Technologies InfinityLab LC/MSD single quadrupole LC/MS (ESI source).

**Scheme:**

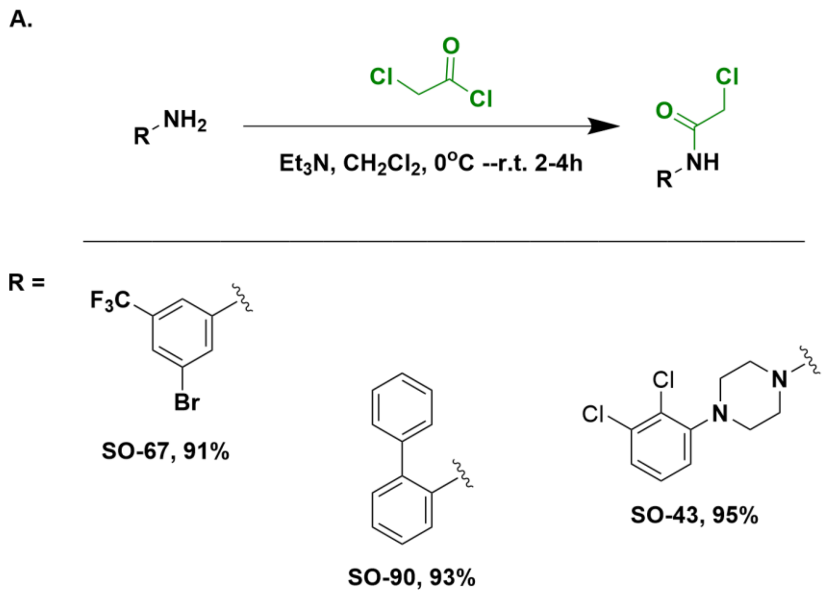

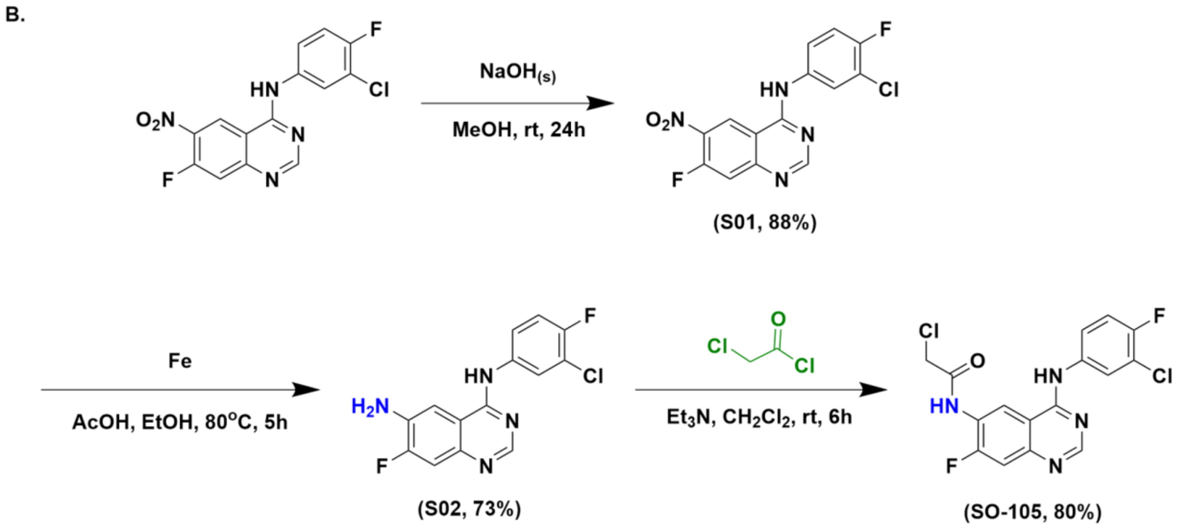


**Scheme S1.** Synthetic routes to obtain (A) electrophilic fragments SO-67, SO-90, and SO-43, and (B) prototype kinase inhibitor SO-105.

#### Synthesis

##### General Procedure 1:

To a solution of the amine (1 equiv.) and triethylamine (1.5 equiv.) in dichloromethane (5 mL), 2-chloroacetyl chloride (1.1 equiv.) was added dropwise and the reaction mixture was stirred at 0°C for 0.5 h then left to warm to room temperature (2 - 6h). The reaction progress was monitored via TLC. On reaction completion, the reaction mixture was poured slowly into distilled water (10ml). After separation of the phases, the organic layer was dried with anhydrous sodium sulfate, filtered and concentrated to obtain the crude product. This crude product was then coated on silica gel, then purified via flash chromatography (in 15-30% Ethyl acetate-hexanes) to yield the analytically pure product.

##### Synthesis of N-(3-bromo-5-(trifluoromethyl)phenyl)-2-chloroacetamide (SO-67)

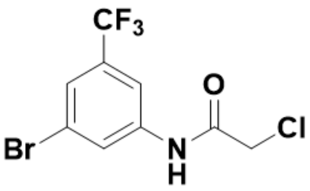

Prepared according to general procedure 1 using 3-bromo-5-(trifluoromethyl)aniline (200mg, 0.833 mmo) as the amine source. Product: white solid . **Yield**: 240 mg (91%)

**$^{1}$H NMR** (400 MHz, $CDCl_3$) δ 8.34 (s, 1H), 8.03 (s, 1H), 7.76 (s, 1H), 7.57 (s, 1H), 4.21 (s, 2H). **$^{13}$C NMR** (100 MHz, $CDCl_3$) δ 164.09, 138.38, 133.17, 132.84, 132.51, 132.20, 132.13, 125.93, 124.93, 123.23, 115.40, 42.73. **$^{19}$F NMR** (400 MHz, $CDCl_3$) δ -62.94.

HRMS (ESI-TOF) $[M+H]^+$ = $C_9H_7BrClF_3NO^+$ : calculated for 315.9305 ; Found 315.9352

**Synthesis of 2-chloro-1-(4-(2,3-dichlorophenyl)piperazin-1-yl)ethan-1-one (SO-43)**

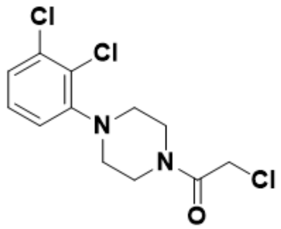

Prepared according to general procedure 1 using 1-(2,3-Dichlorophenyl)piperazine.HCl (300 mg, 1.12 mmol) as the amine source. Product: off-white solid. **Yield:** 95% (326mg).

**$^1$H NMR** (400 MHz, $CDCl_3$) δ 7.24 – 7.14 (m, 2H), 6.94 (dd, J = 7.8, 1.7 Hz, 1H), 4.12 (s, 2H), 3.77 (dt, J = 43.0, 4.9 Hz, 4H), 3.16 – 3.00 (m, 4H). **$^{13}$C NMR** (101 MHz, $CDCl_3$) δ 169.96, 165.55, 150.36, 134.26, 127.63, 125.43, 118.86, 51.47, 50.92, 46.67, 42.47, 40.76.

HRMS (ESI-TOF) $[M+H]^+$ = $C_{12}H_{14}Cl_3N_2O^+$ : calculated for 307.0172 ; Found 307.0040

**Synthesis of N-([1,1'-biphenyl]-2-yl)-2-chloroacetamide (SO-90)**

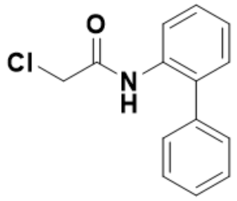

Prepared according to general procedure 1 using [1,1'-biphenyl]-2-amine (300 mg, 1.12 mmol) as the amine source. Product: off-white solid. **Yield:** 93% (410mg).

**$^1$H NMR** (400 MHz, $CDCl_3$) δ 8.46 (s, 1H), 8.36 (d, J = 8.2 Hz, 1H), 7.53 – 7.47 (m, 2H), 7.45 – 7.37 (m, 4H), 7.30 (dd, J = 7.6, 1.7 Hz, 1H), 7.24 (dd, J = 7.4, 1.2 Hz, 1H), 4.08 (s, 2H). **$^{13}$C NMR** (101 MHz, $CDCl_3$) δ 163.64 , 137.43 , 133.92 , 132.65 , 130.14 , 129.15 , 128.22 , 124.98 , 120.64 , 43.03.

HRMS (ESI-TOF) $[M+H]^+$ = $C_{14}H_{12}ClNO^+$ : calculated for 246.0607 ; Found 246.0623

**Synthesis of N-(3-chloro-4-fluorophenyl)-7-methoxy-6-nitroquinazolin-4-amine (S01)**

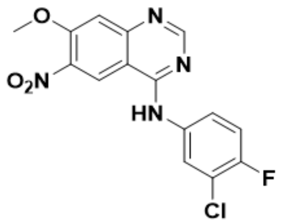

To the N-(3-chloro-4-fluorophenyl)-7-fluoro-6-nitroquinazolin-4-amine (1 g, 2.97 mmol, 1 eqv.) in methanol (23.7 mL, 551 mmol, 200 eqv.) was added sodium hydroxide pellets (1.2 g, 29.8 mmol, 10 eqv.). The reaction was refluxed overnight and monitored by TLC. Next, the reaction mixture was taken off the heat source and allowed to cool to room temperature. Then, the mixture was poured into a saturated solution of sodium bicarbonate, and then filtered under vacuum. The resultant residue was washed with water (1x), then methanol (200 mL). Then the solid residue was dried under high vacuum to yield the target compound as yellowish solid.. **Yield:** 88% (0.91g).

**$^1$H NMR** (400 MHz, DMSO-$d_6$) δ 9.09 (s, 1H), 8.51 (s, 1H), 8.05 (s, 1H), 7.71 – 7.64 (m, 1H), 7.43 – 7.28 (m, 2H), 4.02 (s, 3H). **$^{13}$C NMR** (100 MHz, DMSO-$d_6$) δ 158.12, 154.93, 154.15, 152.39, 138.63, 124.23, 123.18, 122.64, 119.22, 116.72, 109.59, 57.41.
**LC-MS (ESI) $[M+H]^+$** = $C_{15}H_{11}ClFN_4O_3^+$ : calculated for 348.04 ; Found 348.0

**Synthesis of $N^4$-(3-chloro-4-fluorophenyl)-7-methoxyquinazoline-4,6-diamine (S02)**

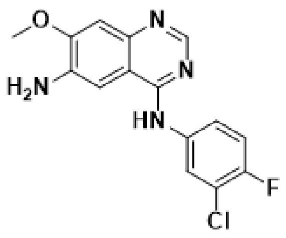


Procedure: To a stirred mixture of N-(3-chloro-4-fluorophenyl)-7-methoxy-6-nitroquinazolin-4-amine (800 mg, 0.435 mol) in ethanol (100 mL) and AcOH (5 mL) was added Fe (641 mg, 11.5 mmol) and the reaction was heated to reflux. Further, more ethanol (50 mL) and AcOH (5 mL) were added, and reflux continued for 5 h, with reaction progress monitored by LC-MS. The reaction was cooled to room temperature, when deemed complete. Next, the crude solution was filtered by passing it through celite. The filtrate was then concentrated to about one-third its original volume. The resultant precipitate was isolated and dried under high vacuum to give the target compound as a dark-brownish solid, (530mg, 73%) – confirmed by LC/MS – and this was used, as is, without further purification. **$^1$H NMR** (400 MHz, DMSO-$d_6$) δ 9.38 (s, 1H), 8.28 (d, J = 76.7 Hz, 2H), 7.81 (d, J = 8.9 Hz, 1H), 7.24 (d, J = 112.2 Hz, 3H), 5.37 (s, 2H), 3.96 (s, 3H). **$^{13}$C NMR** (101 MHz, DMSO-$d_6$) δ 153.20 , 150.73 , 137.99 , 122.92 , 106.36 , 101.30 , 56.25 .

LC-MS (ESI) $[M+H]^+$ = $C_{15}H_{13}ClFN_4O^+$ : calculated for 318.01; Found 319.01

**Synthesis of 2-chloro-N-(4-((3-chloro-4-fluorophenyl)amino)-7-methoxyquinazolin-6-yl) acetamide (SO-105)**

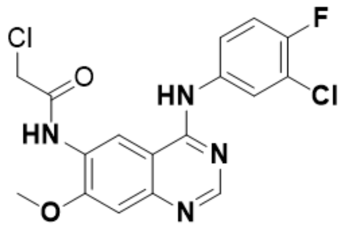


Prepared according to general procedure 1 using N$^4$-(3-chloro-4-fluorophenyl)-7-methoxyquinazoline-4,6-diamine (500 mg, 1.57 mmol) as the amine source. Product: pale-brownish solid. **Yield:** 80% (500mg).

**$^1$H NMR** (400 MHz, DMSO-$d_6$) δ 9.99 (s, 1H), 8.94 (s, 1H), 8.63 (s, 1H), 8.06 (dd, J = 6.8, 2.6 Hz, 1H), 7.75 (dd, J = 6.7, 2.3 Hz, 1H), 7.45 (s, 1H), 7.34 (s, 1H), 4.49 (s, 2H), 4.27 (s, 1H), 4.04 (s, 3H). **$^{13}$C NMR** (101 MHz, DMSO-$d_6$) δ 169.04, 165.68, 157.83, 155.99, 153.41, 152.99, 136.53, 127.56, 125.01, 123.77, 119.36, 116.89, 116.19, 108.83, 105.43, 56.97, 43.78.

**HRMS** (ESI-TOF) $[M+H]^+$ = $C_{17}H_{14}Cl_2FN_4O_2^+$ : calculated for 395.0478 ; Found 395.0389

$^{1}H$ NMR of N-(3-bromo-5-(trifluoromethyl)phenyl)-2 -chloroacetamide, **SO-67** in $CDCl_3$

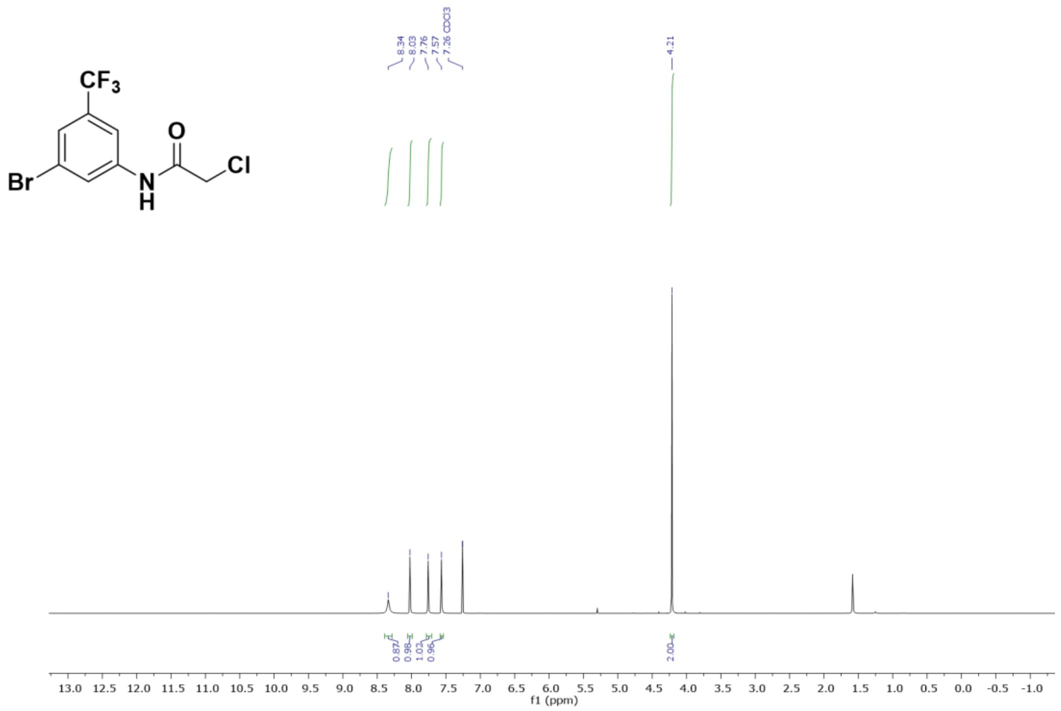


$^{13}C$ NMR of N-(3-bromo-5-(trifluoromethyl)phenyl)-2 -chloroacetamide, **SO-67** in $CDCl_3$

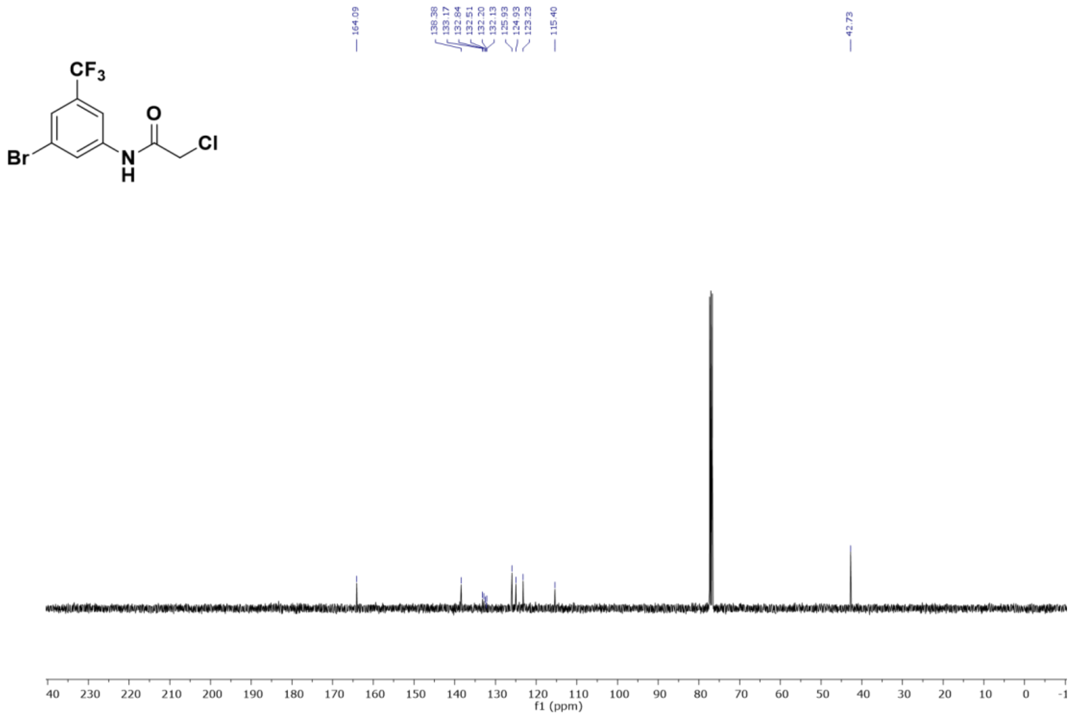

$^{19}F$ NMR of N-(3-bromo-5-(trifluoromethyl)phenyl)-2 -chloroacetamide, **SO-67** in $CDCl_3$

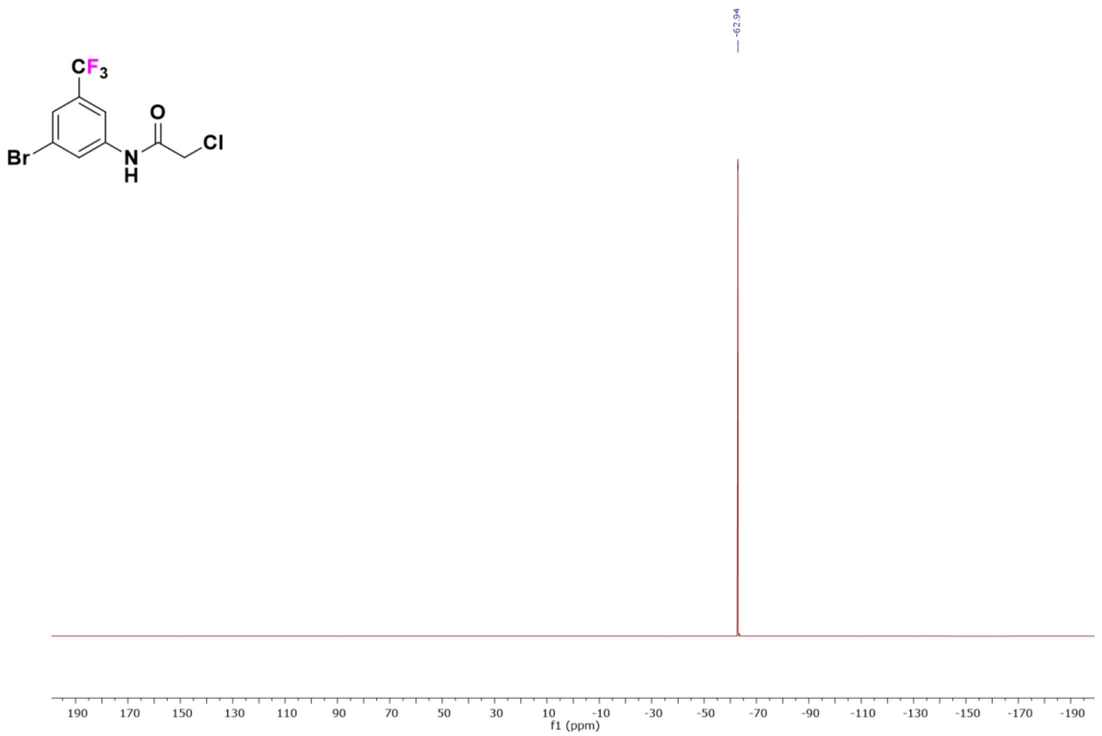

$^{1}H$ NMR of 2-chloro-1-(4-(2,3-dichlorophenyl)piperazin-1-yl)ethan-1-one**, SO-43** in $CDCl_3$

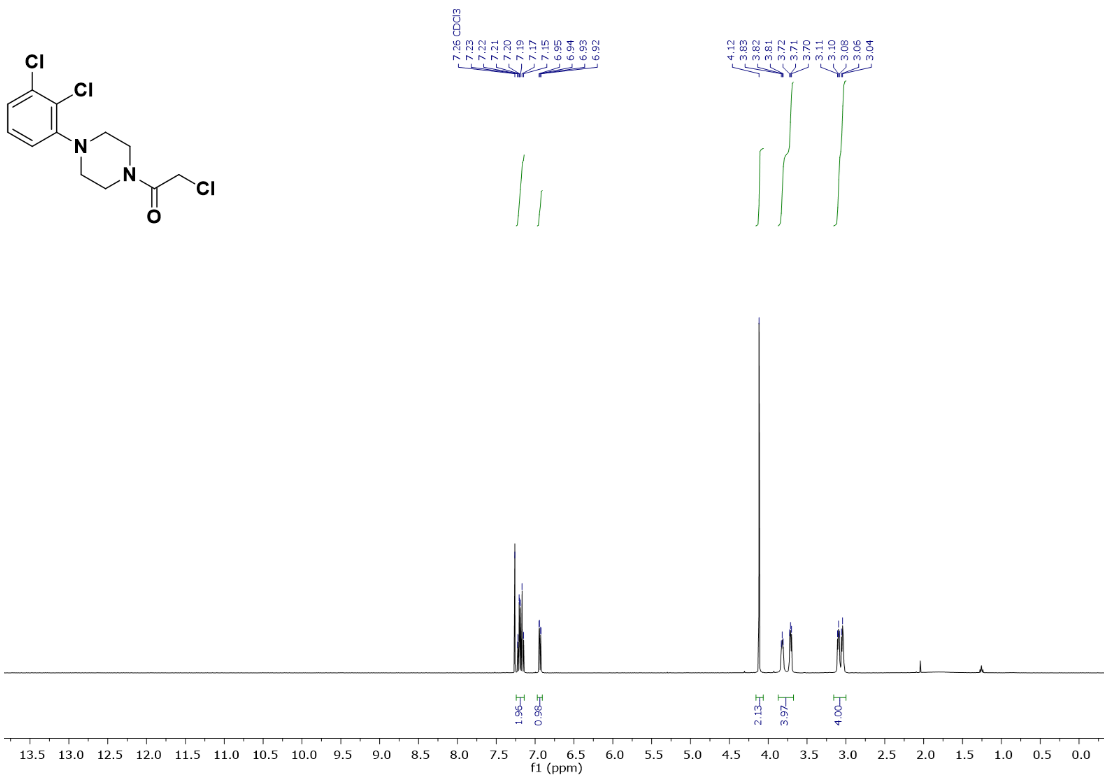

$^{13}C$ NMR of 2-chloro-1-(4-(2,3-dichlorophenyl)piperazin-1-yl)ethan-1-one**, SO-43** in $CDCl_3$

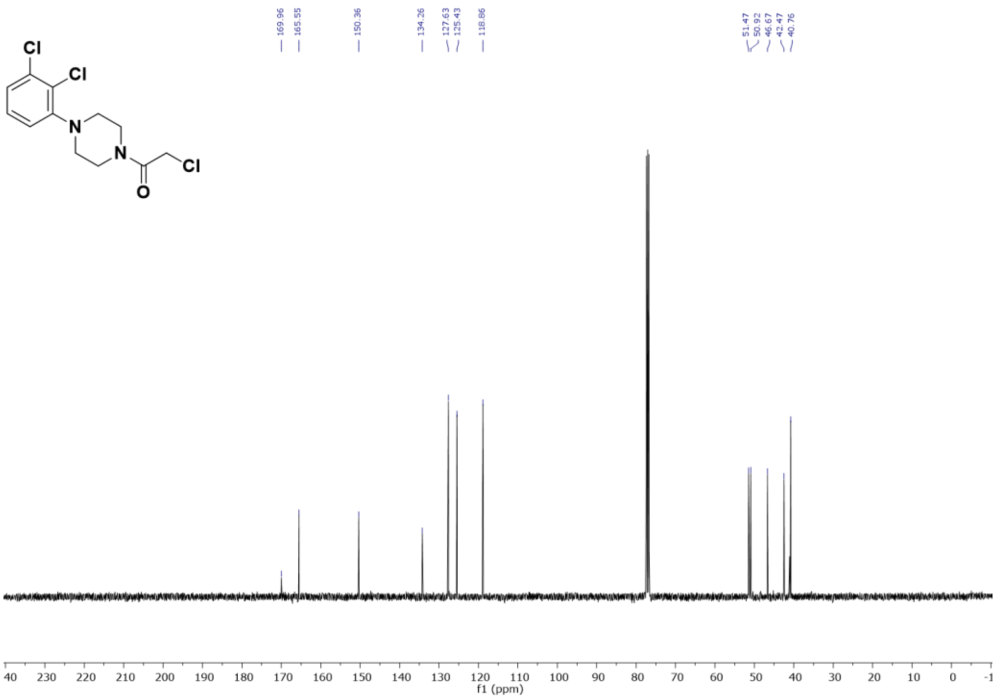


$^{1}H$ NMR of N-([1,1'-biphenyl]-2-yl)-2-chloroacetamide , **SO-90** in $CDCl_3$

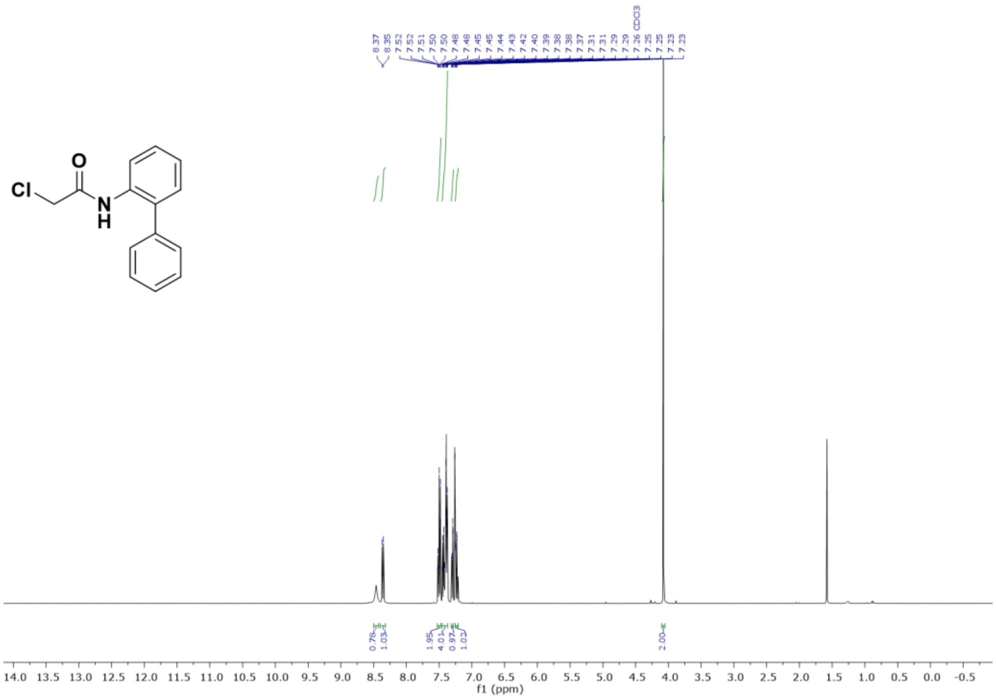

$^{13}C$ NMR of N-([1,1'-biphenyl]-2-yl)-2-chloroacetamide , **SO-90** in $CDCl_3$

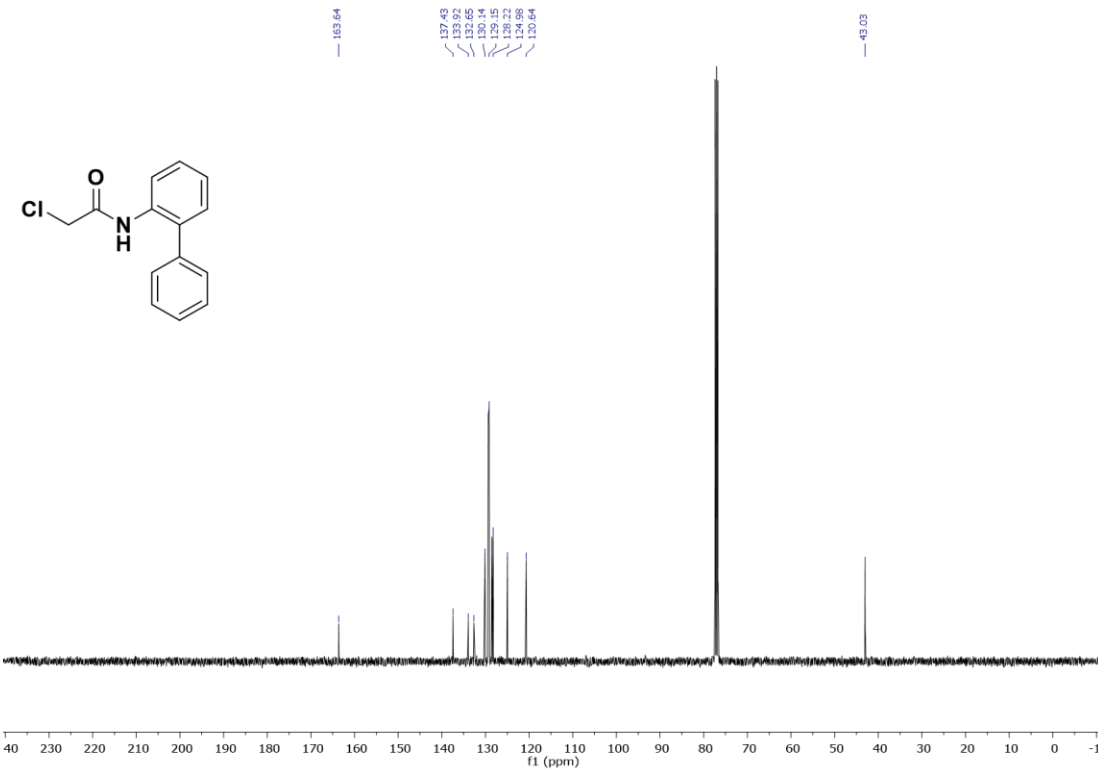


$^{1}H$ NMR of N-(3-chloro-4-fluorophenyl)-7-methoxy-6-nitroquinazolin-4-amine **(S-01)** in $CDCl_3$

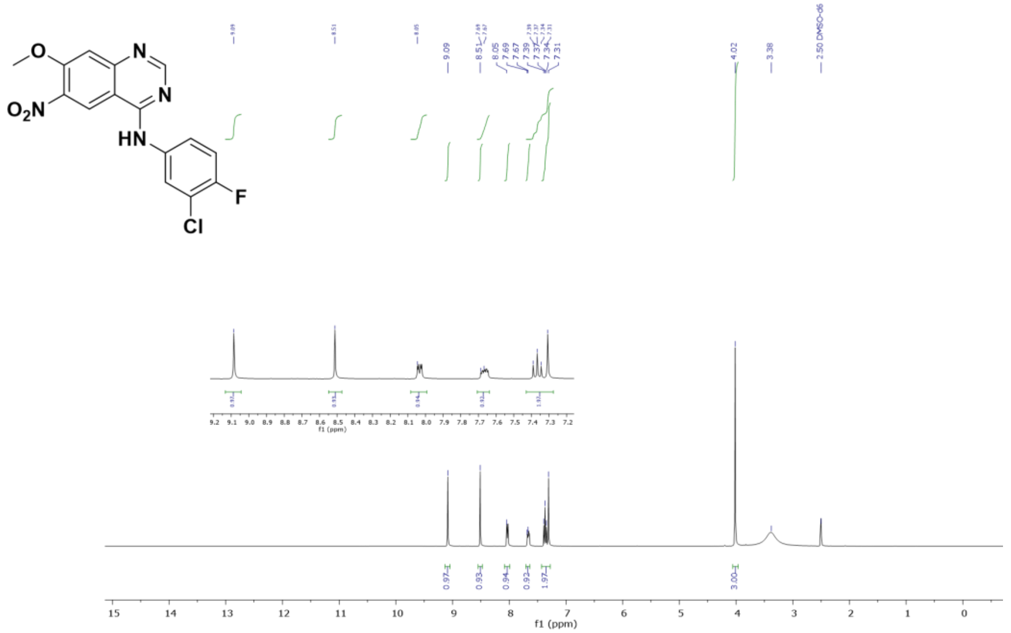

$^{13}C$ NMR of N-(3-chloro-4-fluorophenyl)-7-methoxy- 6-nitroquinazolin-4-amine **(S-01)** in $CDCl_3$

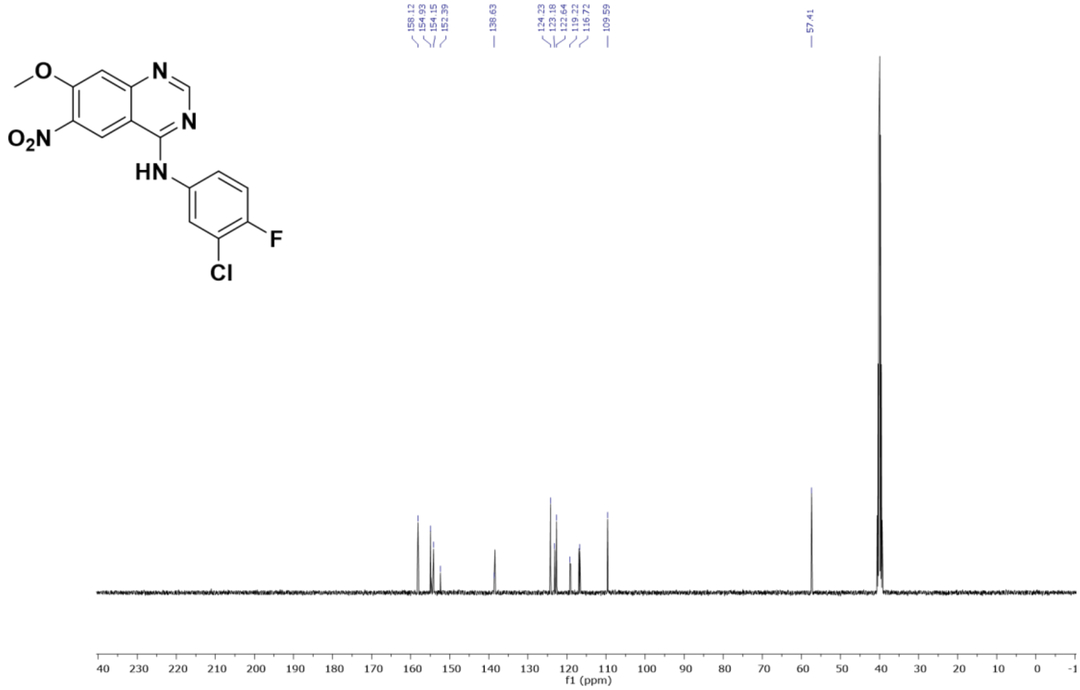


$^{1}H$ NMR of $N^4$-(3-chloro-4-fluorophenyl)-7-methoxyquinazoline- 4,6-diamine **(S-02) , in** DMSO-$d_6$

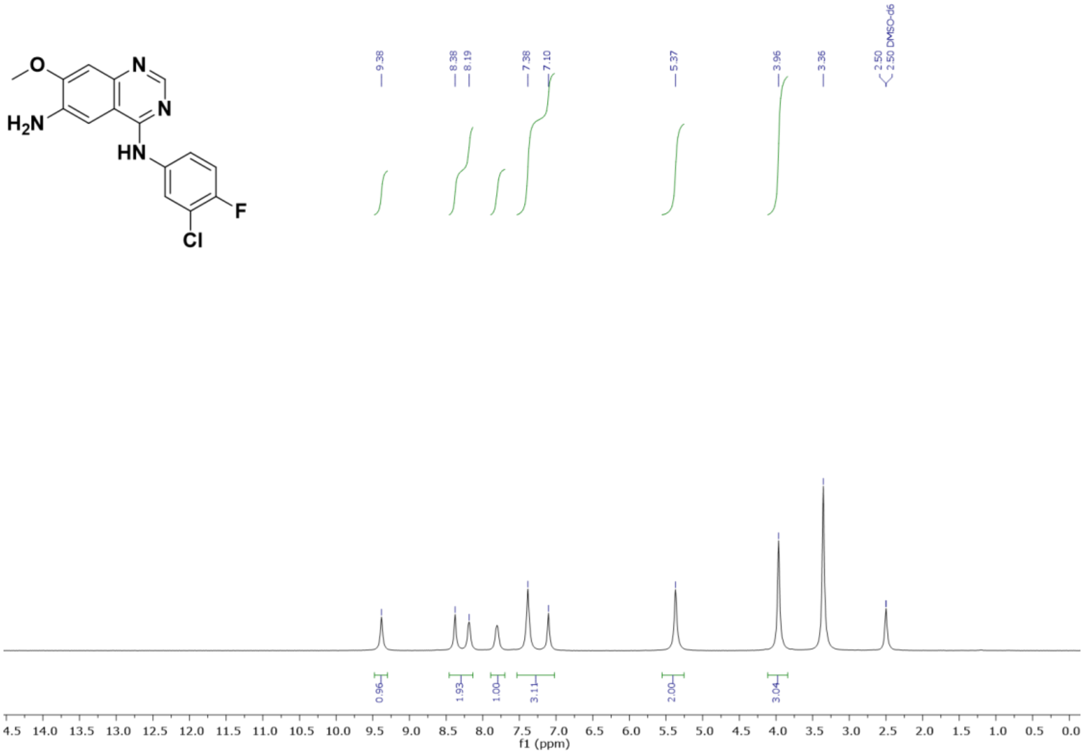

[13]C NMR of N[4]-(3-chloro-4-fluorophenyl)-7-methoxyquinazoline-4,6-diamine **(S-02) ,** in DMSO-*d*$_6$

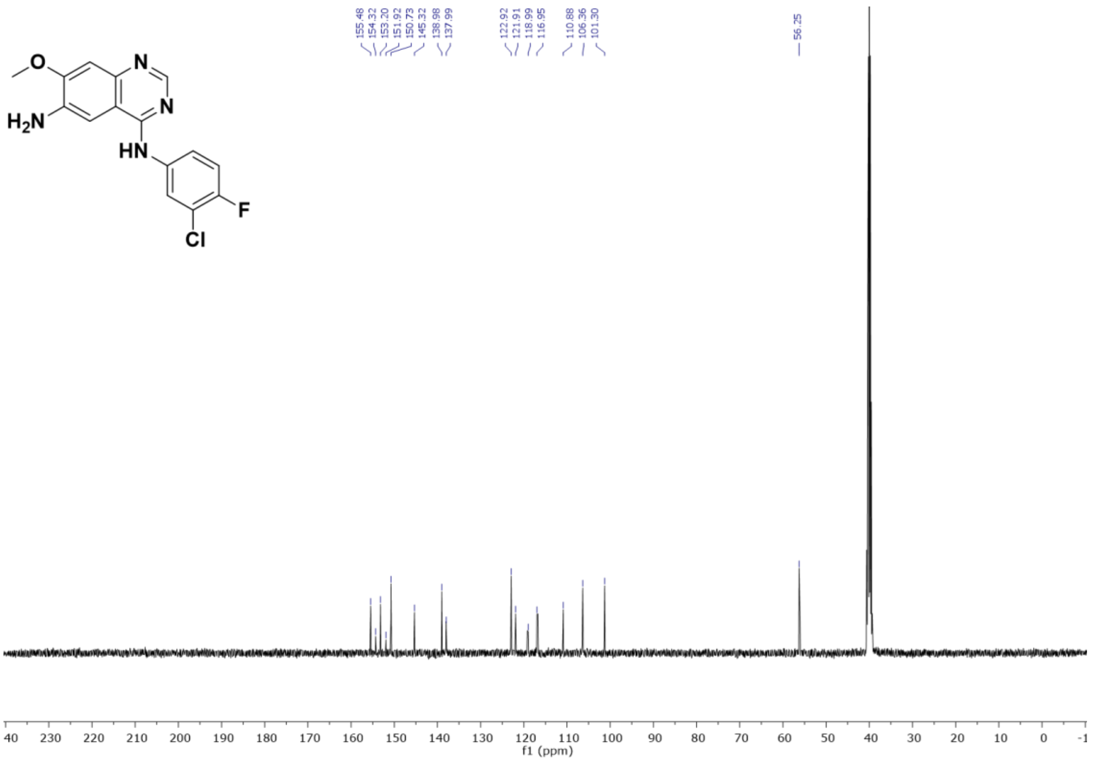


[1]H NMR of 2-chloro-N-(4-((3-chloro-4-fluorophenyl)amino)-7-methoxyquinazolin-6-yl) acetamide, **SO-105, in** DMSO-*d*$_6$

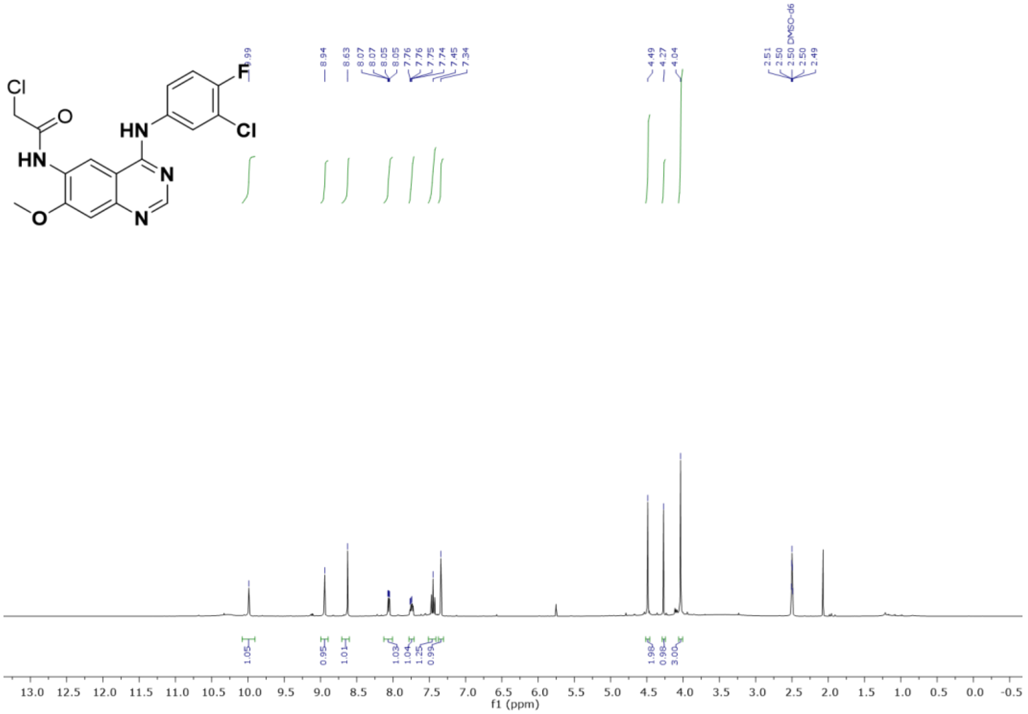

$^{13}C$ NMR of 2-chloro-N-(4-((3-chloro-4-fluorophenyl)amino)-7-methoxyquinazolin-6-yl) acetamide, **SO-105**, in DMSO-$d_6$

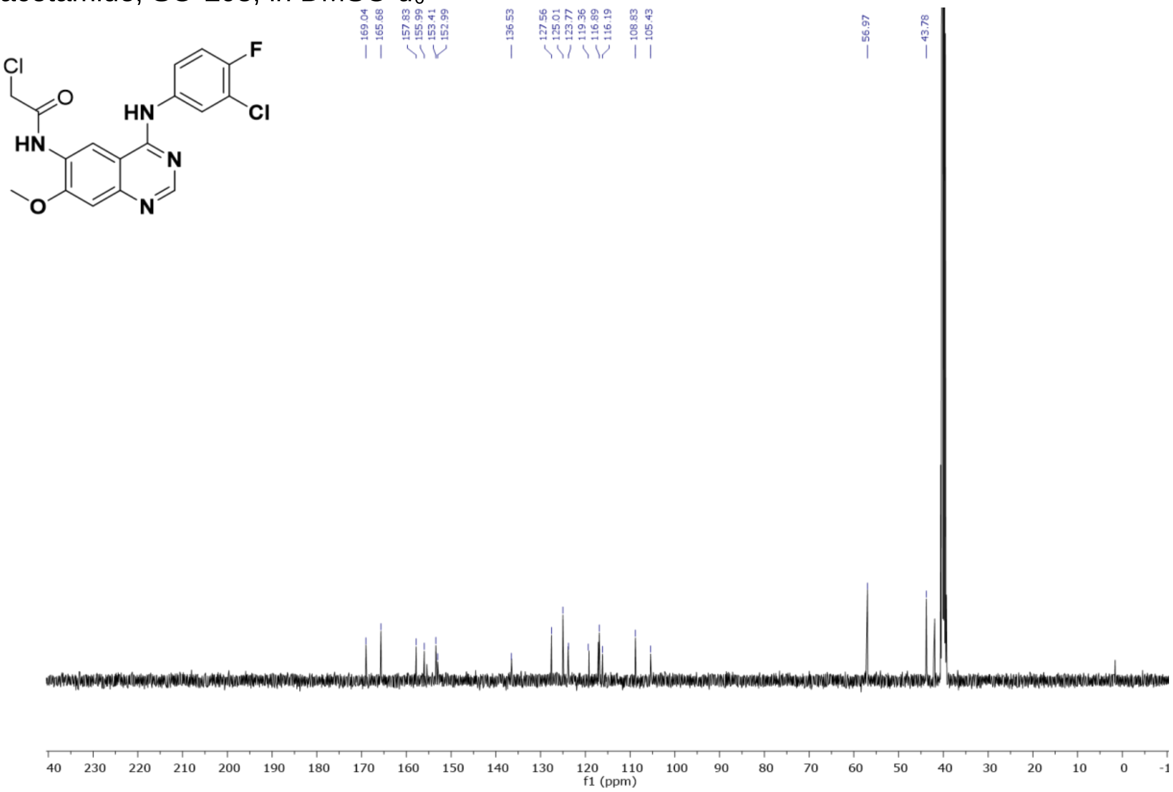


### (C) Running 2-stage search with FragPipe v20+

***Make sure the tools are up-to-date***

*FragPipe v20.0+, MSFragger version 3.8+, Philosopher version 5.0.0+*

1. Set up a search as normally done with the normal reference database and with experiment specific parameters (Percolator rescoring recommended).
2. Select ‘Print decoys’ in the Validation tab
3. In the last "Run” tab, there is a checkbox called "write sub mzML":

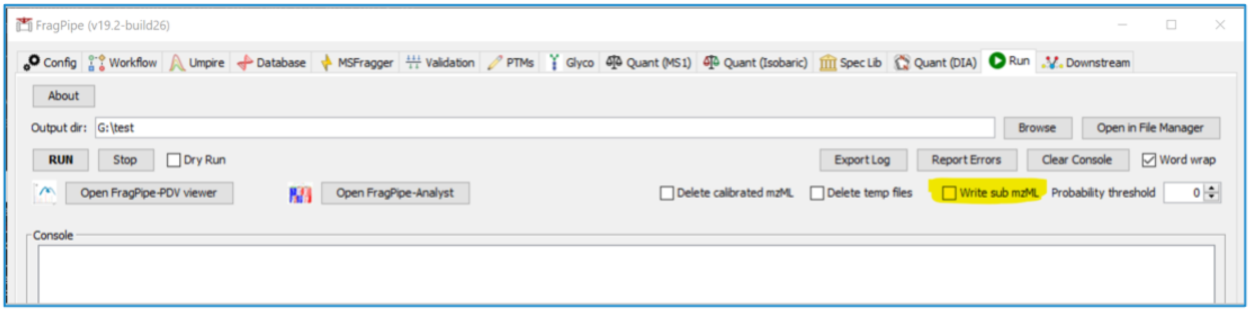


Check this box and leave the threshold set to 0.

*When it is enabled, FragPipe runs the search, and writes new mzML files with "_sub.mzML" as the file name suffix, "fragpipe-second-pass.workflow", and "fragpipe-files-second-pass.fp-*

*manifest" files to the specified result folder. In the first-pass search, FragPipe will write decoy PSMs to psm.tsv if they pass 1% FDR.*

*Both "fragpipe-second-pass.workflow" and "fragpipe-files-second-pass.fp-manifest" have adjusted parameters and mzML file paths that can be directly used in the second search.*

4. Name the result folder to distinguish it from the reference search (“first_pass”, "reference", “1”) and run the first search.
5. After the first search is finished, in your open FragPipe window, navigate to the workflow tab and load the “fragpipe-second-pass.workflow” located in your first results folder file by selecting the "custom" and clicking "load workflow". Select the fragpipe-second-pass.workflow which should then be automatically loaded.

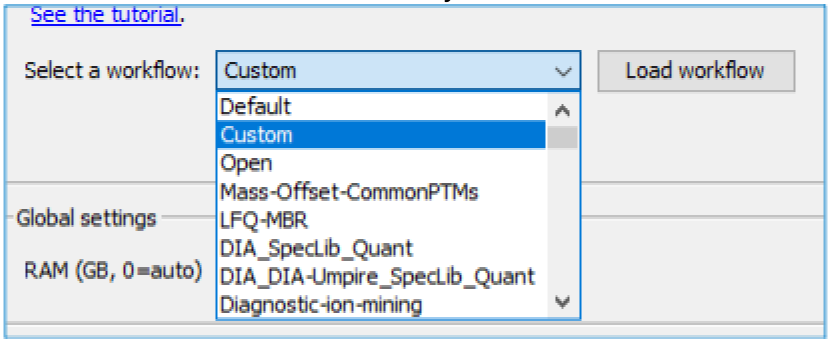


*Note if using Percolator: Percolator weights saving and loading is automatic. FragPipe lets Percolator save the weight files during the first-pass search. And then, in the second-pass, FragPipe gives Percolator the weight files if it can find it (the weight files are in the sub mzML files' folder)*

6. Clear all the .raw/mzML files you have loaded. Load the “fragpipe-files-second-pass.fp-manifest” by selecting load manifest and navigating to it in your first results folder. This should re-add your files with the “_sub.mzML” suffix.

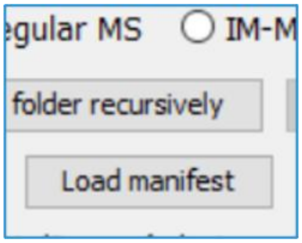


*Note: You should not remove/add any new modifications in the second search. The MSFragger parameters should be the same, just a new database and using _sub.mzML files*

7. Go to the Database tab and navigate to the variant-containing database for your cell line.
8. Specify the reverse header (ex. “REV”).

*Note for databases: For FASTA databases lacking header information such as simple Uniprot IDs (>PXXXX), additional processing/mapping is required to obtain variant IDs.*

9. Make sure the MSFragger parameters are the same as the first search.
10. Navigate to the “Run” tab, uncheck the write sub mzML box, and specify a new results folder to distinguish variant search (“second_pass” or "variant", “2”) and run the second search.

### (D) References


1. Obenchain, V. *et al.* VariantAnnotation: a Bioconductor package for exploration and annotation of genetic variants. *Bioinformatics* **30**, 2076–2078 (2014).
2. Dobin, A. *et al.* STAR: ultrafast universal RNA-seq aligner. *Bioinformatics* **29**, 15–21 (2013).
3. Li, H. *et al.* The Sequence Alignment/Map format and SAMtools. *Bioinformatics* **25**, 2078–2079 (2009).
4. Danecek, P. *et al.* Twelve years of SAMtools and BCFtools. *GigaScience* **10**, giab008 (2021).
5. Oikkonen, L. & Lise, S. Making the most of RNA-seq: Pre-processing sequencing data with Opossum for reliable SNP variant detection. *Wellcome Open Res.* **2**, 6 (2017).
6. Rimmer, A. *et al.* Integrating mapping-, assembly- and haplotype-based approaches for calling variants in clinical sequencing applications. *Nat. Genet.* **46**, 912–918 (2014).
7. McKenna, A. *et al.* The Genome Analysis Toolkit: A MapReduce framework for analyzing next-generation DNA sequencing data. *Genome Res.* **20**, 1297–1303 (2010).
8. Bray, N. L., Pimentel, H., Melsted, P. & Pachter, L. Near-optimal probabilistic RNA-seq quantification. *Nat. Biotechnol.* **34**, 525–527 (2016).
9. Love, M. I., Huber, W. & Anders, S. Moderated estimation of fold change and dispersion for RNA-seq data with DESeq2. *Genome Biol.* **15**, 550 (2014).
10. Lawrence, M. *et al.* Software for Computing and Annotating Genomic Ranges. *PLOS Comput. Biol.* **9**, e1003118 (2013).
11. Durinck, S. *et al.* BioMart and Bioconductor: a powerful link between biological databases and microarray data analysis. *Bioinformatics* **21**, 3439–3440 (2005).
12. UniProt Consortium, T. UniProt: the universal protein knowledgebase. *Nucleic Acids Res.* **46**, 2699 (2018).
13. Kong, A. T., Leprevost, F. V., Avtonomov, D. M., Mellacheruvu, D. & Nesvizhskii, A. I. MSFragger: Ultrafast and comprehensive peptide identification in mass spectrometry-based proteomics. *Nat. Methods* **14**, 513–520 (2017).
14. Yu, F., Haynes, S. E. & Nesvizhskii, A. I. IonQuant Enables Accurate and Sensitive Label-Free Quantification With FDR-Controlled Match-Between-Runs. *Mol. Cell. Proteomics* **20**, 100077 (2021).
15. Yu, F. *et al.* Fast Quantitative Analysis of timsTOF PASEF Data with MSFragger and IonQuant. *Mol. Cell. Proteomics* **19**, 1575–1585 (2020).
16. Leprevost, F. da V. *et al.* Philosopher: a versatile toolkit for shotgun proteomics data analysis. *Nat. Methods 2020 179* **17**, 869–870 (2020).
17. Keller, A., Nesvizhskii, A. I., Kolker, E. & Aebersold, R. Empirical Statistical Model To Estimate the Accuracy of Peptide Identifications Made by MS/MS and Database Search. *Anal. Chem.* **74**, 5383–5392 (2002).
18. Perez-Riverol, Y. *et al.* The PRIDE database and related tools and resources in 2019: Improving support for quantification data. *Nucleic Acids Res.* **47**, D442–D450 (2019).
19. Hatos, A. *et al.* DisProt: intrinsic protein disorder annotation in 2020. *Nucleic Acids Res.* **48**, D269–D276 (2020).
20. Quaglia, F. *et al.* DisProt in 2022: improved quality and accessibility of protein intrinsic disorder annotation. *Nucleic Acids Res.* **50**, D480–D487 (2022).
21. Piovesan, D. *et al.* DisProt 7.0: a major update of the database of disordered proteins. *Nucleic*

*Acids Res.* **45**, D219–D227 (2017).
22. Hornbeck, P. V., Chabra, I., Kornhauser, J. M., Skrzypek, E. & Zhang, B. PhosphoSite: A bioinformatics resource dedicated to physiological protein phosphorylation. *Proteomics* **4**, 1551–1561 (2004).
23. Rainer, J., Gatto, L. & Weichenberger, C. X. ensembldb: an R package to create and use Ensembl-based annotation resources. *Bioinformatics* **35**, 3151–3153 (2019).
